# The extrachromosomal circular DNAs of the rice blast pathogen *Magnaporthe oryzae* contain a wide variety of LTR retrotransposons, genes, and effectors

**DOI:** 10.1101/2021.10.12.464130

**Authors:** Pierre M. Joubert, Ksenia V. Krasileva

**Affiliations:** Department of Plant and Microbial Biology, University of California, Berkeley, CA 94720, USA

**Keywords:** Extrachromosomal circular DNA, fungal plant pathogen, LTR retrotransposons, rice blast

## Abstract

**Background:** One of the ways genomes respond to stress is by producing extrachromosomal circular DNAs (eccDNAs). EccDNAs can contain genes and dramatically increase their copy number. They can also reinsert into the genome, generating structural variation. They have been shown to provide a source of phenotypic and genotypic plasticity in several species. However, whole circularome studies have so far been limited to a few model organisms. Fungal plant pathogens are a serious threat to global food security in part because of their rapid adaptation to disease prevention strategies. Understanding the mechanisms fungal pathogens use to escape disease control is paramount to curbing their threat.

**Results:** We present a whole circularome sequencing study of the rice blast pathogen *Magnaporthe oryzae*. We find that *M. oryzae* has a highly diverse circularome containing many genes and showing evidence of large LTR retrotransposon activity. We find that genes enriched on eccDNAs in *M. oryzae* occur in genomic regions prone to presence-absence variation and that disease associated genes are frequently on eccDNAs. Finally, we find that a subset of genes is never present on eccDNAs in our data, which indicates that the presence of these genes on eccDNAs is selected against.

**Conclusions:** Our study paves the way to understanding how eccDNAs contribute to adaptation in *M. oryzae*. Our analysis also reveals how *M. oryzae* eccDNAs differ from those of other species and highlights the need for further comparative characterization of eccDNAs across species to gain a better understanding of these molecules.

## Background

Extrachromosomal circular DNAs (eccDNAs) are a broad and poorly understood category of molecules defined simply by the fact that they are circular and originate from chromosomal DNA. This group of molecules has been referred to by many names and includes many smaller categories of molecules such as episomes, double minutes, small polydisperse circular DNAs, and microDNAs. They form through several mechanisms including non-allelic homologous recombination (HR), double strand break repair, replication slippage, replication fork stalling, R-loop formation during transcription [1], and as a byproduct of LTR retrotransposon activity [2–4] (Fig. 1A). EccDNAs can accumulate in cells through autonomous replication [5–8], high rates of formation [9], or through retention in ageing cells [10]. EccDNAs can contain genes, and amplification of gene-containing eccDNAs has been linked to adaptation to copper [9] and nitrogen [5] stress in yeast, herbicide resistance in weeds [6], and drug resistance in cancer cells [11, 12]. EccDNA formation is thought to sometimes cause genomic deletions [5,13,14] and reinsertion of eccDNAs after their formation has also been thought to generate structural variation [15, 16]. Some evidence also indicates that eccDNAs could facilitate horizontal gene transfer [16]. Despite their potential as important facilitators of genetic and phenotypic plasticity and presence in all eukaryotes, research efforts, and especially whole circularome sequencing experiments, have been limited to model organisms and human cancer. Therefore, how these molecules behave across the tree of life and how different species could take advantage of these molecules to rapidly adapt to their environments have remained largely unknown.

**Fig. 1.**
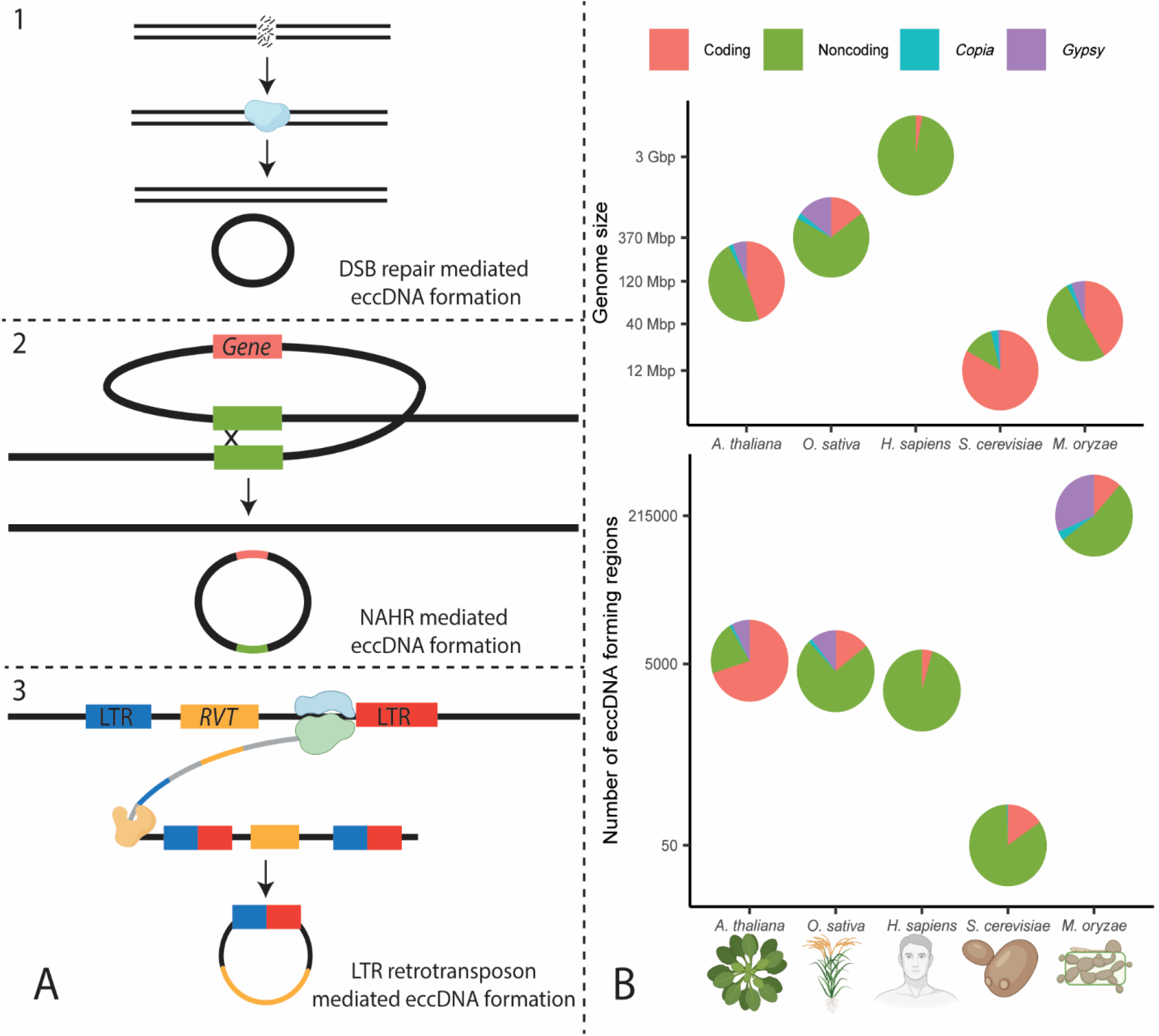
Comparison of eccDNA formation in *M. oryzae* and other organisms. **A.** Examples of mechanisms of extrachromosomal circular DNA (eccDNA) formation. **1.** eccDNA formation as a result of double strand break repair. The blue enzyme represents several different types of DNA repair mechanisms **2.** eccDNA formation as a result of nonallelic homologous recombination (NAHR). The green boxes represent homologous sequences. **3.** eccDNA formation as a result of LTR retrotransposon activity. The blue and green enzyme represents RNA polymerase, and the orange enzyme represents a reverse transcriptase (RVT). Rectangles that are partly blue and partly red represent hybrid LTRs formed from 5’ and 3’ LTRs during retrotransposition. DNA is drawn in black and RNA in gray. **B.** Comparison of genome size and number of eccDNA forming regions for *Arabidopsis thaliana* [17], *Oryza sativa* [18], *Homo sapiens* [13], *Saccharomyces cerevisiae* [19], and *Magnaporthe oryzae.* The number of eccDNA forming regions are shown as called by our pipeline in an average sample. Circularome data for *A. thaliana* and *O. sativa* leaf tissue, *H. sapiens* muscle tissue, and *S. cerevisiae* deletion collection samples are shown. The organism and protein icons were created with BioRender.com.

One of the greatest threats to food security is the devastation of crops by fungal plant pathogens. These pathogens secrete molecules known as effectors to modify host functions and cause disease [20]. The most promising solution to these diseases is the genetic modification of crops by introducing new disease resistance genes, often by allowing the crops to detect effectors and trigger immune responses [21]. Unfortunately, the deployment of disease resistant crops has often had only short-term impacts as some fungal pathogens have adapted to these defenses in very short time spans [22]. Similarly, fungicides are often used to mitigate the devastation caused by pathogens but fungi often evolve drug resistance [23]. A better understanding of how these pathogens adapt and overcome disease prevention efforts so quickly is vital to implementing future strategies. Sequencing and characterization of the genomes of fungal plant pathogens have implicated transposable elements [24], accessory chromosomes [25, 26], and horizontal gene transfer [27]. Additionally, the compartmentalized genome architectures of some of these pathogens, commonly referred to as the “two-speed” genome, is thought to facilitate adaptation to stress by harboring stress response genes and disease associated genes, including effectors, in rapidly evolving regions of their genomes that contain few genes and many repetitive elements [28]. Given the potential for eccDNAs to be a source of phenotypic and genotypic plasticity, we sought to characterize the circularome of one of these pathogens to identify if eccDNAs could play a role in the rapid adaptation of the fungal plant pathogen, *Magnaporthe oryzae* (syn. *Pyricularia oryzae*).

*M. oryzae*, is the causative agent of the rice blast disease [29], has been described as one of the most important fungal pathogens threatening agriculture [30] and is responsible for losses in rice crops equivalent to feeding 60 million people each year [31]. Its ease of culture as well as the importance of this pathogen for global food security have propelled it to being one of the most studied plant pathogens resulting in over three hundred sequenced genomes as well as transcriptomic, and epigenetic datasets in addition to genetic tools including CRISPR/Cas9 mediated genome editing [32]. The availability of these extensive genomic datasets makes *M. oryzae* a prime candidate for understanding the role eccDNAs may play in adaptation to stress in a fungal plant pathogen.

We present here our analysis of circularome sequencing data for *M. oryzae* and identify eccDNA forming regions in its genome. We describe the high diversity of eccDNA forming regions that we found in the rice blast pathogen and compare it to previously sequenced circularomes. We find that most of the *M. oryzae* circularome is made up of LTR retrotransposon sequences and that genes on eccDNAs tend to originate from regions of the genome prone to presence-absence variation. Additionally, our characterization of the genes found on eccDNAs shows that many genes are never found on eccDNAs under the conditions we tested and suggests that selection may shape which genes are found on these molecules. Finally, our analysis reveals that many disease-causing effectors are found on eccDNAs in the pathogen.

## Results

### Identification of eccDNA forming regions in *Magnaporthe oryzae*

To characterize the circularome of *M. oryzae*, eccDNAs were purified and sequenced from pure cultures of *M. oryzae* Guy11 using a protocol adapted from previously published methods [18]. Briefly, after total DNA extraction of 3 biological replicates, linear DNA was degraded from 3 technical replicates for each biological replicate using an exonuclease and the remaining circular DNA was amplified using rolling circle amplification (RCA). Depletion of linear DNA was verified with qPCR using markers to the *M*. *oryzae* actin gene (MGG_03982, Additional File 1: Fig. S1). This gene was used as a marker for linear DNA since increased copies of the ACT1 gene are thought to be deleterious in yeast [19, 33]. Isolated eccDNAs were then sequenced using both paired-end Illumina sequencing and PacBio circular consensus sequencing (CCS). In total, we sequenced 8 samples as one technical replicate failed quality checks during library preparation. Illumina sequencing yields were 6.5 Gbp per sample, on average, and PacBio sequencing yields were 8 Gbp (subreads) and 500 Mbp (CCS) per sample, on average.

To identify specific breakpoints indicating eccDNA formation in our Illumina sequencing data, we developed a pipeline inspired by previously published methods [13]. In circularome sequencing data, split mapping reads originate from sequencing circularization junctions of eccDNAs. Additionally, read pairs in the data that map in the opposite direction represent sequencing from paired-end sequencing fragments that span these circularization junctions. Our pipeline uses split reads in combination with opposite facing read pairs to find evidence of eccDNA formation (Fig. 2). This allowed us to identify, with high confidence, genomic sequences belonging to eccDNAs, which we will hereafter refer to as “eccDNA forming regions.” We will refer to split reads associated with these eccDNA forming regions simply as “junction split reads.” Our analysis was limited to these eccDNA forming regions, rather than the fully resolved structure of each eccDNA molecule, because of the complexity of eccDNAs as well as the techniques used into sequenced them in this study. For example, eccDNAs can sometimes contain multiple copies of the same sequence [34] and our use of RCA, which generates long DNA fragments containing hundreds of tandem repeats of each circular molecule [35], prevents determination of whether a sequence is repeated many times on an eccDNA molecule or just present once. Additionally, eccDNAs have also been shown to assemble with others, forming complex structures [36]. While our long-read PacBio sequencing may have been able to address this issue, our attempts at reference-free assembly of complete eccDNAs were unsuccessful, likely due to insufficient coverage of each molecule. While only eccDNA forming regions could be described in this study, these regions still enable a detailed description of the *M. oryzae* circularome. Across all 8 sequenced samples, our pipeline identified 1,719,878 eccDNA forming regions using Illumina paired-end sequencing data (Additional File 3). We validated 8 of these eccDNA forming regions using outward PCR and Sanger sequencing (Fig. 2 and Additional File 1: Fig. S2). These regions were chosen for validation as they fully contained genes of interest to the rest of the study, including well-known effectors.

**Fig 2.**
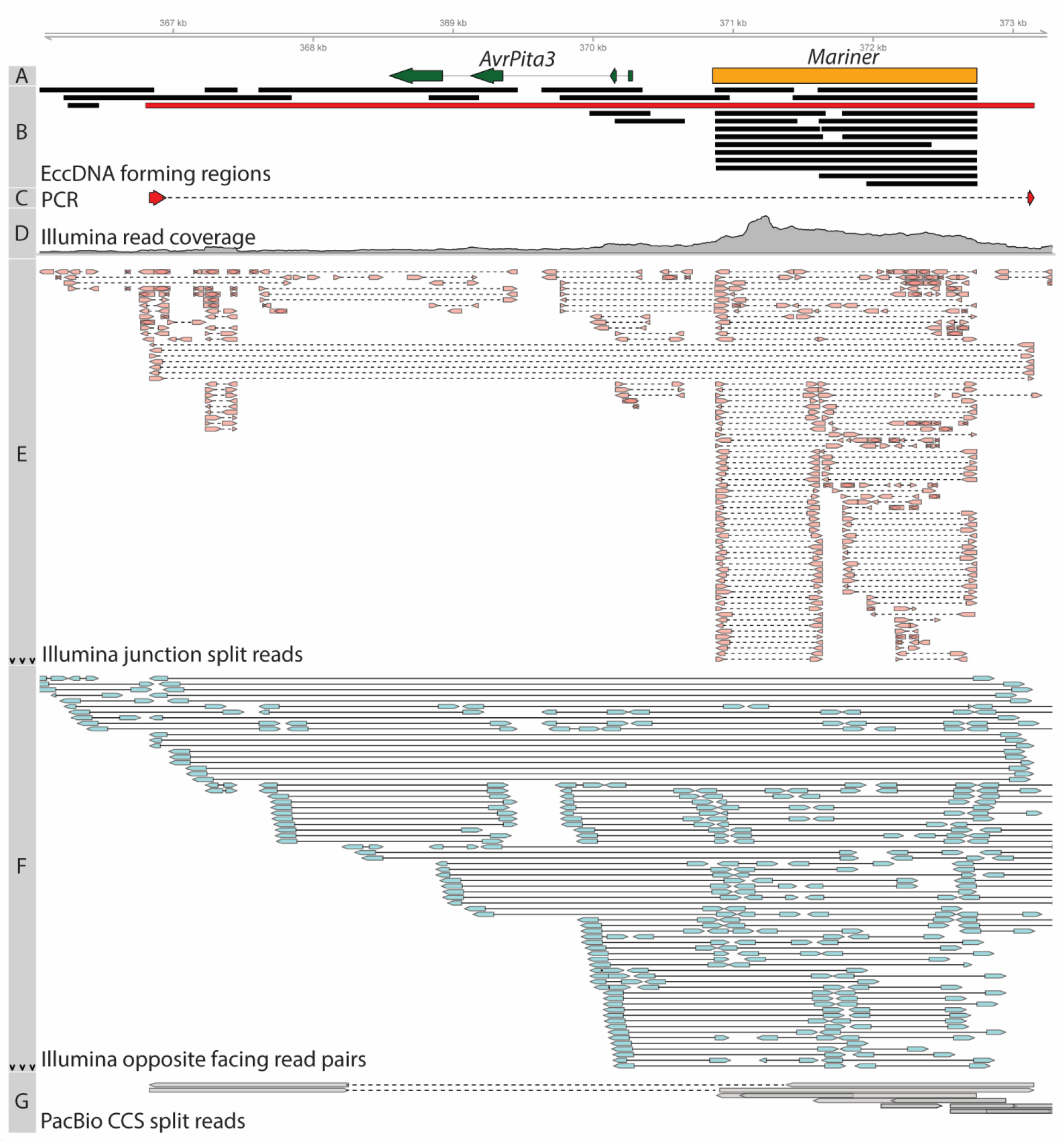
Summary of evidence supporting an eccDNA forming region of interest in the *M. oryzae* genome. **A.** Location of effector *AvrPita3* and *Mariner* transposon. **B.** Location of eccDNA forming regions. The eccDNA forming region in red was chosen for validation using outward PCR. This eccDNA forming region was considered to fully encompass *AvrPita3*. **C.** Sanger sequencing read generated from outward PCR (Additional File 1: Fig S2) that supports eccDNA forming region highlighted in red in track B. **D.** Overall Illumina sequencing read coverage. **E.** Junction split reads obtained from Illumina data. Split reads are joined by a dashed line. Black arrows indicate not all reads were shown in areas with high counts. **F.** Opposite facing read pairs obtained from Illumina data. Read pairs are joined by a solid line. Black arrows indicate that not all reads were shown in areas with high counts. **G.** Split reads obtained from PacBio CCS data. Overlapping arrows indicate single reads mapped to the same location more than once. Split reads are joined by a dashed line. All data was obtained from a single sequenced sample (biological replicate 1, technical replicate A).

To determine how similar our technical and biological replicates were to each other, we compared the coordinates of eccDNA forming regions found in each sample. Overall, we found little overlap in eccDNA forming regions between technical replicates (14.16%, 10.09%, and 23.77%, for biological replicates 1, 2 and 3, respectively) and between biological replicates (9.41%) when comparing the exact start and end coordinates of these regions (Additional File 1: Fig. S3). Rarefaction analysis showed that these differences could be, at least partially, attributed to under sequencing, though this data could also be evidence of many low copy number eccDNAs being produced by the *M. oryzae* genome (Additional File 1: Fig. S4). However, principal component analysis using the coverage of junction split reads throughout the genome showed that technical replicates were more likely to be similar to other technical replicates within the same biological replicate than across biological replicates in the content of their eccDNA forming regions (Additional File 1: Fig. S5). Additionally, while exact coordinates of eccDNA forming regions did not have much overlap between samples, considering eccDNA forming regions whose start and end coordinates were within 100 bp of each other in two different samples to be the same increased this overlap greatly between technical replicates (48.46%, 45.55%, and 58.29% for biological replicates 1, 2 and 3, respectively) and between biological replicates (42.89%) (Additional File 1: Fig. S6). We performed a permutation analysis to simulate random formation of eccDNAs throughout the genome to verify that this result was meaningful and observed little overlap between replicates in this simulated scenario when increasing our overlap tolerance up to 100bp (Additional File 1: Fig. S6). All together, these results, as well as others presented throughout this study suggested that while the exact breakpoints of eccDNA forming regions were not identical across samples, the genomic loci, or hotspots, of eccDNA formation were highly similar.

Likely due to the great number of different eccDNAs in *M.* oryzae, the coverage of our PacBio sequencing data was too low to enable *de novo* assembly of eccDNA molecules. Therefore, we used our long read data to infer eccDNA forming regions by mapping them to the *M. oryzae* Guy11 genome and comparing these regions to those called using our short read data. This was done using a similar pipeline to the Illumina data with less stringent criteria which was better adapted to the lower read depth of the long read data. Our long read data allowed us to identify 147,335 eccDNA forming regions across all samples (Additional File 4). We compared these eccDNA forming regions to those called using Illumina data, allowing for up to a 10 bp difference between breakpoints to account for mapping ambiguity, and found that, on average, 81.42% of eccDNA forming regions called using PacBio data for one sample were also found in our eccDNA forming regions called using Illumina reads in the same sample (Additional File 1: Fig. S7). We were able to attribute much of this discrepancy to our stringent criteria for calling eccDNA forming regions since simply looking for split reads in our Illumina data increased this rate to 90.36% (Additional File 1: Fig. S7). The remaining differences are likely due to Illumina reads not being long enough to properly be mapped as split reads in certain regions of the genome. Such strong overlap between eccDNA forming regions called by long reads and short reads demonstrates the robustness of our short read data analysis. Aside from this validation, we chose not to include the PacBio data in our final analyses due to the low read depth.

Next, we quantified the potential false positive rate of our pipeline that could have originated from any undigested genomic DNA in our samples by running the pipeline on previously published whole genome sequencing data from *M. oryzae* Guy11 [32,37,38]. Based off the number of eccDNA forming regions called from this data, we estimated this false positive rate to be approximately 3 junction split reads per million sequencing reads (Additional File 2: Table S1). In comparison, we found 41,873 junction split reads per million reads in our eccDNA enriched samples, on average, indicating a very low false positive rate from our pipeline. Additionally, we could not completely rule out the presence of eccDNAs in the whole genome sequencing samples we analyzed. This validation showed that any remaining linear DNA in our samples after linear DNA degradation were unlikely to be called as eccDNA forming regions by our pipeline.

Finally, we benchmarked our pipeline on previously published eccDNA data in human tissue [13] (Additional Files 5 and 6). We found that, on average, 74.62% of eccDNA forming regions called by our pipeline were also described in the published dataset (Additional File 1: Fig. S8A). This number was even higher for eccDNA forming regions associated with 10 or more junction split reads (85.63%). The small fraction of eccDNA forming regions called by our pipeline that did not appear in the published list could not be attributed to how our pipeline handled multi-mapping reads (Additional File 1: Fig. S8A, see Methods) and were likely due to differences in sequence data processing and different criteria for selecting split reads between the two studies [13]. However, the two lists significantly differed in the number of eccDNA forming regions identified, with our pipeline identifying substantially less (Additional File 1: Fig. S8B). This difference can be attributed to our stricter evidence to call eccDNA forming regions. In our method, eccDNA forming regions were only called if split reads mapped to the region. This is in contrast to other methods of calling eccDNA forming regions which rely at least partly on peaks in sequencing coverage [13,19,39]. This meant that our pipeline could not detect eccDNAs formed from HR between identical repeats which do not result in split reads. We chose this method for *M. oryzae* because it showed circularome sequencing coverage throughout the entire genome in our samples and very few clear coverage peaks, which indicates that many low copy number eccDNAs were present in our samples. The high degree of overlap between our called eccDNA forming regions and those described by Møller *et al.* makes us confident that the eccDNA forming regions we called using our pipeline are robust.

### The *M. oryzae* circularome is more diverse and contains more noncoding sequences than the circularomes of other organisms

We were first interested in comparing the circularome of *M. oryzae* to those of other previously characterized organisms. To compare these datasets across different organisms, we gathered sequencing data from several previous studies [13,17–19] and reanalyzed them using our pipeline (Additional Files 5-20). Our analysis revealed a very large number of eccDNA forming regions in *M. oryzae* compared to other previously sequenced organisms (Fig. 1B). We also looked at the percentage of the genome that was found in eccDNA forming regions and found that, while most organisms had 1-10% of their genome in eccDNA forming regions, our samples showed an average of 74.48% of the *M. oryzae* genome in eccDNA forming regions (Additional File 1: Fig. S9A). The difference in the number of eccDNA forming regions between organisms was still striking after normalizing for genome size and sequencing library size (Additional File 1: Fig. S9B). These results supported the idea that the low amount of overlap in eccDNA forming regions between our samples could be explained partly by the great number of eccDNAs produced by the *M. oryzae* genome. While the difference in the number of called eccDNA forming regions could be attributed to differences in the methods used for eccDNA purification (Additional File 2: Table S2), we extracted and sequenced eccDNAs from *Oryza sativa* and found similar levels of diversity to previously published samples (Additional File 1: Fig. S9B). We also found that *M. oryzae* had more eccDNA forming regions made up of noncoding sequences relative to the percentage of noncoding sequence in its genome than other organisms aside from *S. cerevisiae* (Fig. 1B, Additional File 1: Fig. S9C).

### LTR retrotransposon sequences make up most of the *M. oryzae* circularome

*Gypsy* and *Copia* LTR retrotransposons frequently generate eccDNAs through several mechanisms [2–4], so we looked for the presence of these sequences in the *M. oryzae* circularome. Our analysis revealed that 54.12% of the eccDNA forming regions we identified seemed to be composed of more than 90% LTR retrotransposon sequence indicating that these elements made up a large portion of the pathogen’s circularome despite only making up a small fraction of its genome (Fig. 1B, Additional File 1: Fig. S10). Further comparative analysis revealed that a much higher proportion of the *M. oryzae* circularome was made up of these LTR retrotransposon sequences than in other organisms (Fig. 1B, Additional File 1: Fig. S9D and S9E).

All six LTR retrotransposons identified in *M. oryzae* Guy11 formed eccDNAs (Fig. 3A). However, the elements *MAGGY*, *GYMAG1*, and *Copia1* made up the majority of the eccDNA sequencing data (Fig. 3B). When this data was normalized to the proportion of the genome made up by each transposon, *GYMAG1* stood out as making up a much greater percentage of the sequencing data than expected (Fig. 3C, Additional File 1: Fig. S11).

**Fig. 3.**
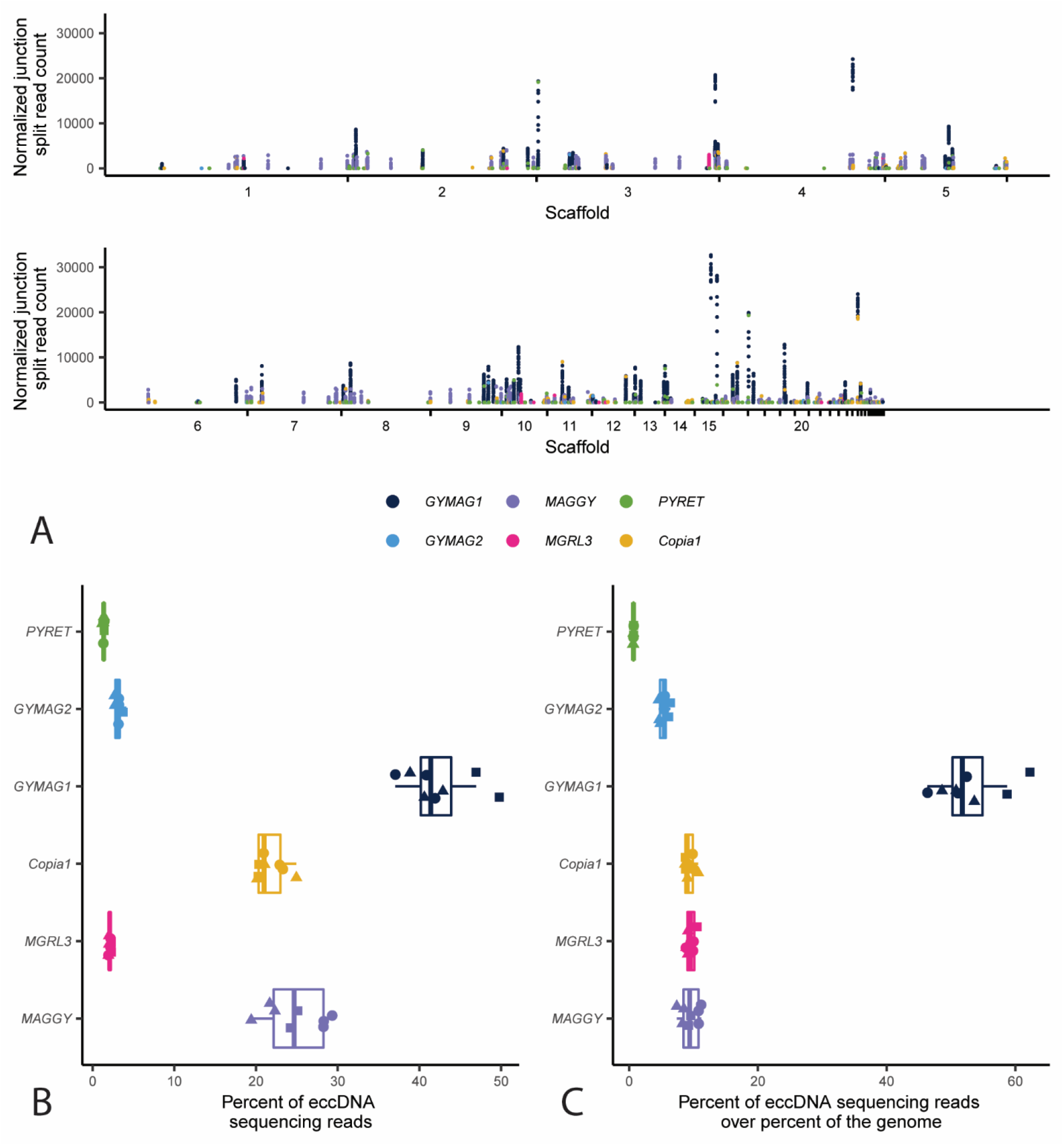
The majority of eccDNAs in *M. oryzae* are made up of LTR retrotransposons. **A.** Manhattan plot showing the number of junction split reads per million averaged across biological replicates for all 100 bp bins that overlap an LTR retrotransposon in the *M. oryzae* Guy11 genome. Each point represents one of these bins. **B.** Boxplot showing the percentage of sequencing reads that map to LTR retrotransposons. Each point represents one sample, and the shape of the points represent the biological replicate that sample was taken from. **C.** Boxplot showing the ratio of the percentage of sequencing reads that map to LTR retrotransposons to the percentage of the *M. oryzae* Guy11 genome that is made up by that retrotransposon. Each point represents one sample, and the shape of the points represent the biological replicate that sample was taken from.

### LTR retrotransposons in *M. oryzae* form eccDNAs through a variety of mechanisms

LTR retrotransposons can form eccDNAs through a variety of mechanisms [2–4]. EccDNA formation commonly occurs after transcription and reverse transcription of the transposon which results in a linear fragment of extrachromosomal DNA [40] (Fig. 1A). Then, the most common circularization mechanisms are nonhomologous end joining (NHEJ) of the two LTR ends to form eccDNAs containing 2 LTRs (scenario 1, Fig. 4A), autointegration of the retrotransposon forming single LTR eccDNAs of various lengths, depending on where in the internal sequence of the transposon the autointegration event happens (scenario 2, Fig. 4B), and HR between the two LTRs to forming single LTR eccDNAs (scenario 3, Fig. 4C). Finally, LTR retrotransposon sequences can also become part of eccDNAs by other eccDNA formation mechanisms that do not rely on retrotransposition activity, such as intrachromosomal HR between solo-LTRs or between multiple copies of the same transposon [4,5,19]. Given this diversity of mechanisms, we wanted to evaluate which of them contributed to eccDNA formation in *M. oryzae.* To do this, we first simulated the expected read coverage for each of the three active LTR eccDNA formation mechanisms under ideal conditions where only one mechanism of formation was occurring (Fig. 4A-C). Then, we measured the prevalence of scenarios 1 and 2 by identifying specific split read variants in our data. LTR eccDNAs formed through NHEJ result in split reads that map to one end of an LTR and the other which we will refer to as LTR-LTR split reads (Additional File 1: Fig. S12 and S14A). Autointegration results in split reads that map to one LTR and to the internal region of the transposon which we will refer to as LTR-internal split reads (Additional File 1: Fig. S13 and S14B). HR between two identical LTRs (scenario 3) would not result in a split read so we could not find this type of evidence in our data.

**Fig. 4.**
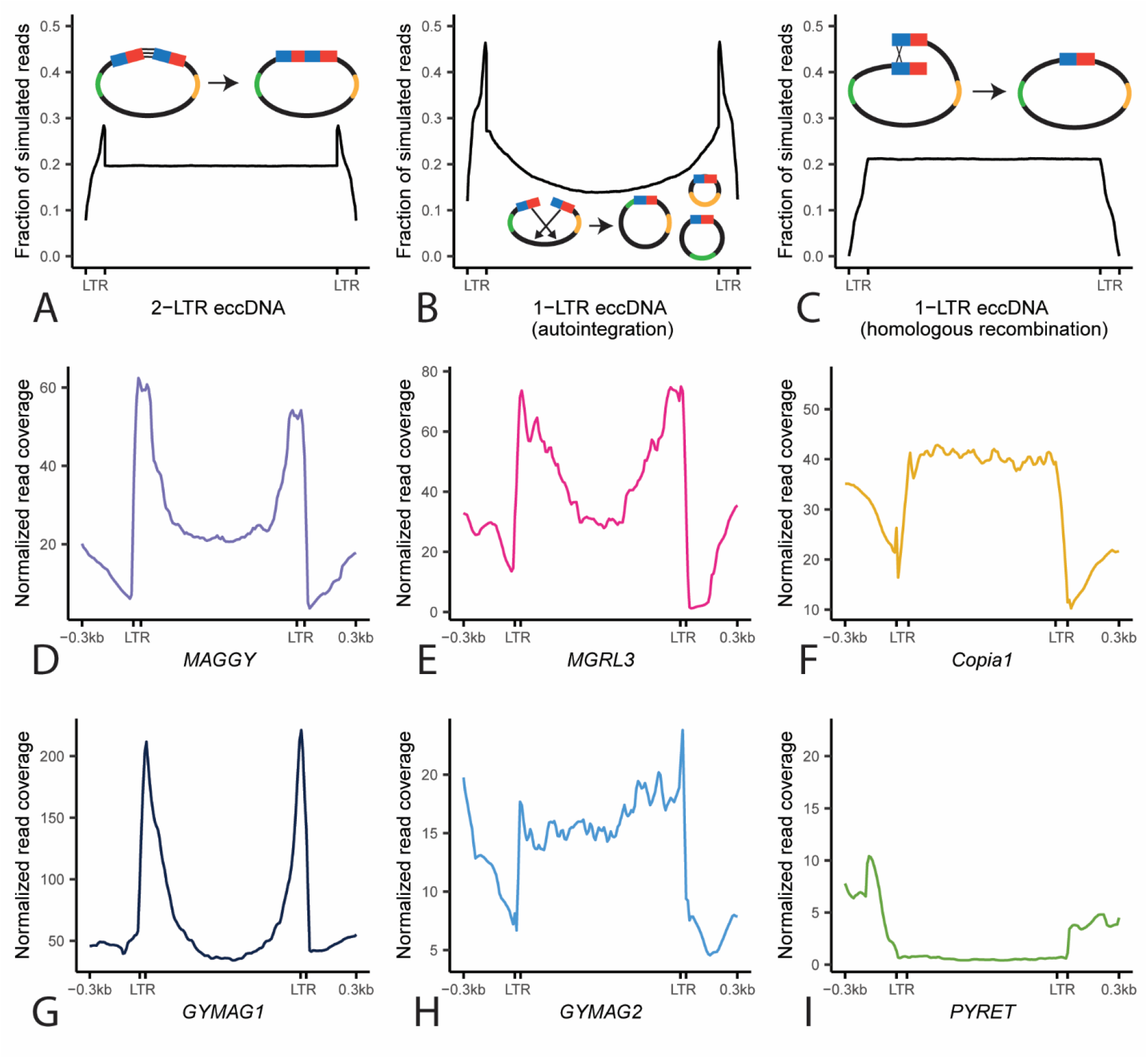
LTR retrotransposons in *M. oryzae* form eccDNAs through a variety of mechanisms. **A-C.** Profile plots showing expected sequencing read coverage for each LTR retrotransposon eccDNA formation scenario as well as graphical representations of the scenario. In the graphics, blue and red rectangles represent hybrid LTRs formed from 5’ and 3’ LTRs during retrotransposition and green and orange lines represent areas of the internal region of the retrotransposon with distinct sequences. **D-I.** Profile plots showing observed sequencing read coverage for each LTR retrotransposon found in the *M. oryzae* Guy11 genome.

Comparisons between simulated and observed read coverage plots revealed contributions of several eccDNA formation mechanisms that varied by transposable element. For *MAGGY*, our analysis indicated that it forms eccDNAs primarily through autointegration (Fig. 4D). This was supported by a high correlation between the number of sequencing reads and LTR-internal split reads (Additional File 1: Fig. S13A) and a low correlation between sequencing reads and LTR-LTR split reads (Additional File 1: Fig. S12A). The data also pointed to *MGRL3* and *GYMAG1* forming eccDNAs primarily through autointegration (Fig. 4E and 4G, Additional File 1: Fig. S12BD and S13BD). *Copia1*, on the other hand showed a clear pattern of read coverage corresponding to eccDNA formation through HR (Fig. 4F), though the high correlation between sequencing reads and LTR-internal split reads mapping to this element hinted that a small, but proportional, fraction of *Copia1* elements formed eccDNAs through autointegration (Additional File 1: Fig. S13C). In the case of *GYMAG2*, its sequencing read coverage resembled a pattern expected for LTR-eccDNAs formed through NHEJ (Fig. 4H). The large amount of LTR-LTR split reads per million mapped reads found corresponding to *GYMAG2* elements compared to other retrotransposons supported this inference (Additional File 1: Fig. S14A). *PYRET*’s distinct sequencing read coverage profile likely indicated that it mostly formed eccDNAs by other eccDNA formation mechanisms that do not rely on retrotransposition activity such as intrachromosomal HR (Fig. 4I). A low correlation between sequencing read coverage and both LTR-LTR split reads and LTR-internal split reads as well as the fragmented nature of *PYRET* elements, which is a sign of low recent retrotransposon activity, supported this inference (Additional File 1: Fig. S12F and S13F). Finally, to determine whether the results we obtained were caused by bias in the length and completeness of the retrotransposon sequences in the *M. oryzae* genome, we generated profile plots for each retrotransposon using previously generated whole genome sequencing data [32,37,38]. The results from this analysis ruled out this possibility (Additional File 1: Figure S15). In conclusion, it is clear that a variety of eccDNA formation mechanisms contributed to eccDNAs containing LTR retrotransposon sequences and that these mechanisms varied by element.

### MicroDNAs are distinct from other eccDNAs

MicroDNAs have previously been studied as a distinct set of molecules within the eccDNA category. Besides being small (less than 400bp), microDNAs are found to be enriched in genic regions, exons, 5’UTRs and CpG islands [14, 41]. We examined if microDNAs in *M. oryzae* showed these characteristics by analyzing eccDNA forming regions less than 400 bp in length with less than 10% LTR retrotransposon sequence across different organisms. Enrichment of microDNAs in CpG islands was the most consistent result across all organisms we analyzed, though this enrichment was not found in *M. oryzae* (Additional File 1: Fig. S16). Similarly, we found no enrichment of microDNAs in 5’UTRs in *M. oryzae*. We did however find a small enrichment of microDNAs in genic regions in *M. oryzae* as in many of the other sequenced organisms (Additional File 1: Fig. S16 and S17). In general, our analysis suggested that the previously described characteristics of microDNAs are not common across all organisms and sample types.

MicroDNAs also displayed distinct features from the remaining subset of non-LTR eccDNAs which we called large eccDNAs. Among other differences, we found that, unlike microDNAs, large eccDNAs tended to be enriched in intergenic regions (Additional File 1: Fig. S17 and S18). Additionally, eccDNAs are often associated with active transcription [1, 9], and we found a slight but significant correlation between expression and junction split reads for large eccDNAs but not for microDNAs (Additional File 1: Fig. S19).

In yeast, eccDNA amplification is thought to often occur with the help of autonomously replicating sequences (ARSs) which contain ARS consensus sequences (ACSs) [5,19,42]. In *M. oryzae*, we found that ACSs were enriched in large eccDNAs (permutation test, mean of expected: 5320.14 regions, observed: 6950 regions, p < 0.01, n = 100 replicates) but depleted in microDNAs (permutation test, mean of expected: 818.09 regions, observed: 714 regions, p < 0.01, n = 100 replicates). However, for both large eccDNAs and microDNAs, presence of an ACS in the eccDNA forming region did not result in an increased number of junction split reads (Additional File 1: Fig. S20). Finally, microDNAs have been found to be associated with chromatin marks and increased GC content [14, 41]. However, we did not find any of these enrichments in microDNAs or large eccDNAs in *M. oryzae* (Additional File 1: Fig. S21).

### Many genes are found encompassed by eccDNA forming regions

Many eccDNAs contain genes and these eccDNAs can provide genotypic and phenotypic plasticity in other organisms. In *M. oryzae* we found that, out of the 12,115 genes in Guy11, 9,866 were fully contained by an eccDNA forming region in at least one sample (for an example, see Fig. 2B). These genes included *TRF1* (MGG_04843) and *PTP2* (MGG_00912) which have been shown to be involved in fungicide resistance in *M. oryzae* [43, 44]. EccDNA forming regions containing these two genes were validated using outward PCR (Additional File 1: Fig. S2). However, not all genes were observed in eccDNA forming regions at the same frequency and their presence on eccDNAs was heterogenous across samples. To further understand what types of genes are enriched in eccDNA forming regions, we focused on a robust set of eccDNA-associated genes. To identify these genes, we first counted the number of times each gene was found fully contained by a junction split read in each sample. We referred to this count as the number of “encompassing split reads” for each gene. We then normalized this count to the number of junction split reads in each sample and averaged it across technical replicates for each biological replicate. Finally, we sorted the genes by their prevalence in each biological replicate and chose genes that were found in the top third of genes for this count in all three biological replicates. In total, using these metrics, we identified 558 eccDNA-associated genes shared across all biological replicates (Additional File 1: Fig. S22 and Additional File 21).

To identify biological processes enriched in eccDNA-associated genes, we performed gene ontology (GO) enrichment analysis. We found that eccDNA-associated genes were enriched for GO terms related to vesicle transport, mitosis, and the cytoskeleton among other terms (Fig. 6A, Additional File 1: Fig. S23 and Additional Files 22-24). We also explored whether eccDNA-associated genes showed differences in gene expression or other genomic features from other genes. However, we found no difference between eccDNA-associated genes and other genes in gene expression, GC content, or histone marks, aside from a significant difference in H3K36me3 (Additional File 1: Fig. S24 and S25).

**Fig. 5.**
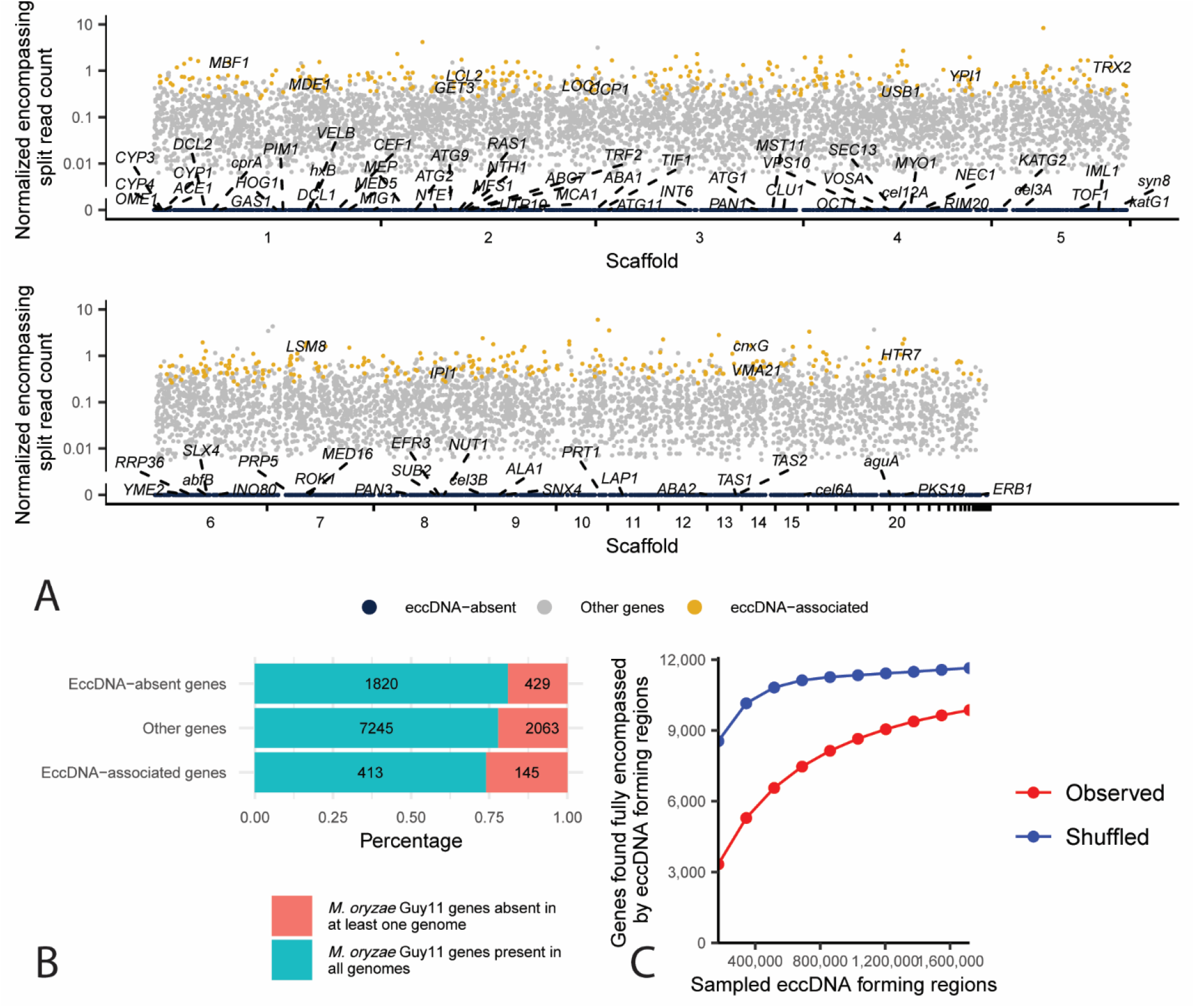
EccDNA forming regions contain most *M. oryzae* genes, but not all, and many are associated with presence-absence variation. **A.** Manhattan plot showing the number of encompassing split reads per million junction split reads averaged across biological replicates for each gene in the *M. oryzae* Guy11 genome. Each dot represents one gene. EccDNA-associated genes with known gene names are labeled according to their normalized encompassing split read count and position in the genome. EccDNA-absent genes with known gene names are labeled with lines pointing to their location in the genome. **B.** Stacked bar plot showing the percentage of eccDNA-absent genes, other genes, and eccDNA-associated genes in the *M. oryzae* Guy11 genome that had an ortholog in all other 162 *M. oryzae* genomes analyzed or not. Numbers indicate the number of genes in each category. **C.** Rarefaction analysis of the observed number of genes found fully encompassed by eccDNA forming regions at different subsamples of all found eccDNA forming regions, compared to the same number of randomly selected genomic regions.

**Fig. 6.**
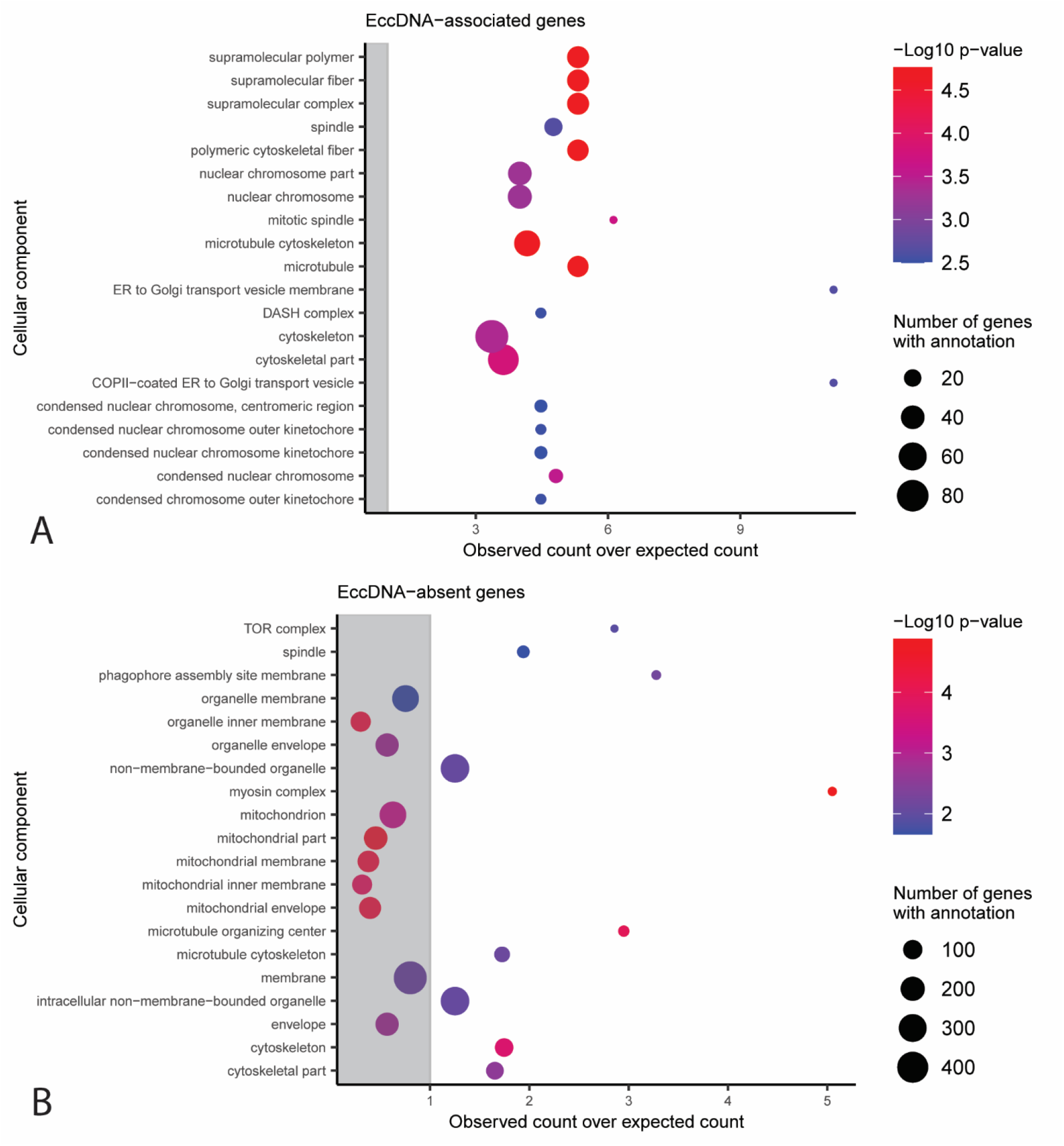
GO terms associated with eccDNA-associated and eccDNA-absent genes in *M. oryzae*. Functional categories in the cellular component Gene Ontology with an observed number of **A.** eccDNA-associated genes or **B.** eccDNA-absent genes that is significantly different from the expected number with correction for gene length bias. The y-axis shows the different functional categories, and the x-axis represents the observed number of genes divided by the expected number of genes in this group. Dots outside of the grey rectangle represent functional categories that are observed more often than expected. The size of dots indicates the total number of genes in the *M. oryzae* genome that belong to each functional category. Only the 20 categories with the largest -log10 p-values according to a Chi-square test are shown.

### EccDNA-associated genes are closer to gene sparse and repeat dense regions of the genome than other genes

Some plant pathogens are described as having “two-speed” genomes with housekeeping genes found close together in repeat-poor regions and environmentally responsive and disease-associated genes found in repeat-dense and gene-poor regions [28]. To determine if eccDNA-associated genes were enriched in either of these genomic contexts, we analyzed if eccDNA-associated genes were more distant from other genes than expected by chance (Fig. 7). We observed a significant difference (permutation test for difference of medians, p = 0.0117, n = 10,000 replicates) between the median distance to the nearest gene of eccDNA-associated genes (543 base pairs) and other genes (485 base pairs). We also observed a significant difference (permutation test for difference of medians, p = 0.0282, n = 10,000 replicates) between the median distance to the nearest genomic repeat of eccDNA-associated genes (663 base pairs) and other genes (769 base pairs, Additional File 1: Fig. S26). This difference in proximity was not observed for transposable elements, indicating that transposable elements alone were not responsible for this effect (Additional File 1: Fig. S27). The heterogeneity of eccDNAs and the mechanisms of their formation might be influencing this comparison. However, our data points to a link between genome architecture and eccDNA formation.

**Fig. 7.**
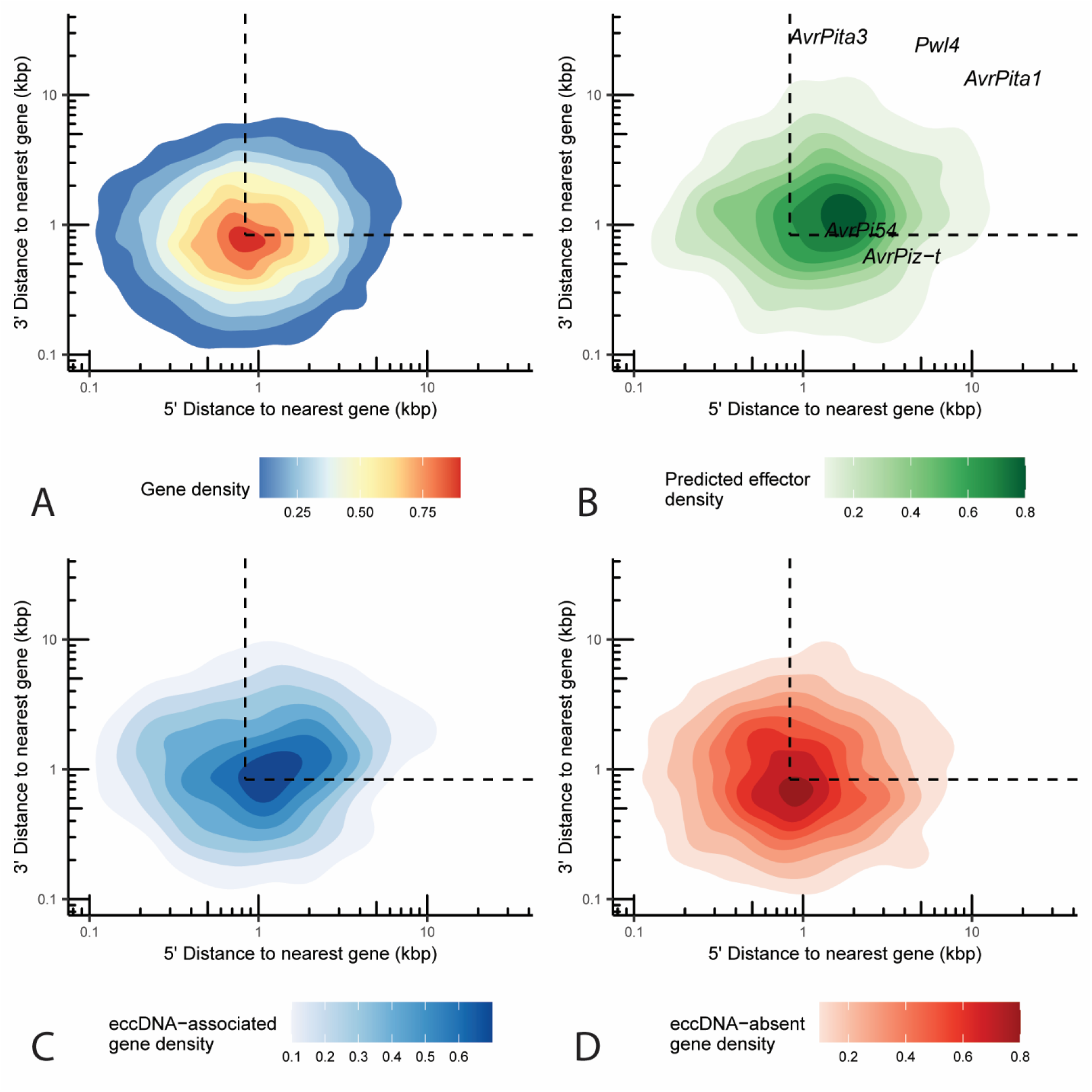
EccDNA-associtated genes are often found in gene sparse regions of the *M. oryzae* genome. Two-dimensional density plot representing the 5’ and 3’ distance to the nearest gene in the *M. oryzae* Guy11 genome in kilobase pairs for each **A.** gene, **B.** predicted effector, **C.** eccDNA-associated genes, and **D.** eccDNA-absent genes. Known effectors are shown as text in **B.** Dashed lines represent median 5’ and 3’ distance to nearest gene.

### EccDNA-associated genes are more prone to presence-absence variation than other genes

There is evidence of eccDNAs generating structural variation in other organisms [15, 16]. We therefore tested whether eccDNA formation is associated with genes prone to presence-absence variation in 162 rice-infecting *M. oryzae* isolates (Additional File 25). As expected from previous studies [45, 46], our analysis indicated that predicted effectors were more likely to experience presence-absence variation (Additional File 1: Fig. S28; X-squared = 146.33, df = 1, p-value < 2.2e-16). We also found that eccDNA-associated genes were more likely to be prone to presence-absence variation (Fig. 5B; X-squared = 16.262, df = 2, p-value = 2.95e-04). This result suggested that eccDNA formation and structural variation occur in similar regions of the genome but did not show whether they are directly linked. To see if a more direct link existed, we surveyed the genomes of the *M. oryzae* isolates for small deletions that completely or partially overlapped genes but did not disrupt neighboring genes. We were able to identify 257 such events (Additional File 26). However, none of these deletions matched our eccDNA forming regions and only 8 of them came within 50 bp. Our rarefaction analyses revealed that there is likely to be a much greater diversity of eccDNAs than what we were able to capture at the sequencing depth of this study, whether we considered samples individually or as a whole (Additional File 1: Fig. S4 and S29). Therefore, eccDNA formation that could have contributed to structural variation might have been missed due to either under sequencing or they could have been missing in the conditions tested in this study.

Similarly, we were interested in identifying any potential DNA translocations that may have occurred through an eccDNA intermediate. While we were able to successfully construct a bioinformatics pipeline that identified one previously described eccDNA-mediated translocation in wine yeast [16] (Additional File 1: Fig. S30), we were unable to identify any such examples in any of the *M. oryzae* genomes we analyzed despite including isolates infecting a variety of hosts in this analysis (306 genomes in total, Additional File 27).

Finally, since mini-chromosomes have been hypothesized as playing important roles in fungal plant pathogen evolution, we also sought to determine whether genes that were previously found on *M. oryzae* mini-chromosomes were associated with eccDNA formation but found no such effect (Additional File 1: Fig. S31).

### Many eccDNA-absent genes are myosin-complex related

Since most *M. oryzae* genes appeared in eccDNA forming regions in at least one sample, we were particularly interested in the 2,249 genes that never appeared fully encompassed by an eccDNA forming region in any of our technical or biological replicates, which we called eccDNA-absent (Additional File 21). We first verified that eccDNA-absent genes were not caused by insufficient sequencing coverage using rarefaction analysis. This analysis differed significantly from our previous ones (Additional File 1: Fig. S4 and S29). Here, we counted the number of genes found in eccDNA forming regions at various subsamples of eccDNA forming regions. This analysis revealed that our observations of eccDNA-absent genes were unlikely to be caused by the under sequencing we described previously as the number of genes found fully encompassed by eccDNA forming regions appeared to plateau at larger subsamples of eccDNA forming regions (Fig. 5C). Additionally, a permutation analysis showed that, given the high coverage of our data, we only expected to find 468 genes in this category by chance, which is far fewer than the 2,249 genes we observed (Fig. 5C).

We next explored whether gene expression or other genomic features could explain the observed eccDNA-absent genes. However, we found no strong differences between eccDNA-absent genes and other genes in gene expression, GC content, or histone marks (Additional File 1: Fig. S24 and S25). EccDNA-absent genes also did not differ from other genes in terms of their distance to the nearest gene, repeat or transposable element (Fig. 7, Additional File 1: Fig. S26 and S27).

Finally, we performed GO enrichment analysis on these genes and found, amongst many other enriched terms, that genes related to cytoskeletal proteins, and especially the myosin complex, were enriched within eccDNA-absent genes (Fig. 6B, Additional File 1: Fig. S32, and Additional Files 28-30). While genes related to the cytoskeleton were also enriched among eccDNA-associated genes, these were related to mitosis and microtubule polymerization, rather than the myosin complex (Fig. 6A, Additional File 1: Fig S23). This result is of particular interest given that the actin gene has been used in a previous study [19] as well as this one, as a marker for linear DNA due to its negative fitness effect at high copy numbers in yeast [33]. As expected, the *M. oryzae* actin gene (MGG_03982) was one of the eccDNA-absent genes, meaning it was never found in an eccDNA forming region in its entirety in any of our samples.

Furthermore, in agreement with our GO enrichment results, *MYO1* was also one of these eccDNA-absent genes. To validate our bioinformatics analysis, we tested whether we could amplify the full sequences of these genes from our eccDNA samples using PCR. In agreement with our findings, we were only able to amplify these sequences from our genomic DNA sample (Fig. S33). These results suggested that eccDNA formation is not random in *M. oryzae* and that certain groups of genes may be protected from eccDNA formation or maintenance of these eccDNAs in the cell.

### Effectors are enriched in eccDNA forming regions compared to other genes

Finally, we wanted to identify whether eccDNA forming regions contained disease-causing effectors. We found that many known *M. oryzae* effectors were encompassed by eccDNA forming regions in at least one sample. This included *AvrPita3*, *AvrPita1*, *AvrPi9*, *AvrPi54*, *AvrPiz-t*, and *Pwl4* (Fig. 2 and 8 and Additional File 21). We validated eccDNA forming regions containing these effectors using outward PCR (Additional File 1: Fig. S2). Additionally, we found that many predicted effectors were found in eccDNA forming regions (Fig. 8 and Additional File 21). We also found that many of these putative effectors were associated with larger numbers of encompassing split reads and found this difference to be statistically significant (Additional File 1: Fig S34; permutation test for difference in medians, p < 0.0001, n = 10,000 replicates). Effectors are often small genes and, given the often-small size of eccDNA forming regions in our data, which may have been caused by the bias of RCA towards small molecules [1, 47] (Additional File 1: Fig. S35), we felt that our analysis could be affected by this bias. To address this issue, we repeated our permutation test, comparing predicted effectors to a set of non-effectors of similar lengths and again found a significant difference in number of encompassing split reads (permutation test for difference in medians with correction for gene length distribution, p = 0.0206, n = 10,000 replicates). This result suggests that effectors are more likely to be found on eccDNAs than other genes in *M. oryzae* and that this effect is not simply due to their size. Additionally, a small proportion of effectors are found among our eccDNA-absent genes (Fig. 8). These candidates might be more evolutionarily stable and therefore useful as targets for disease resistance.

**Fig. 8.**
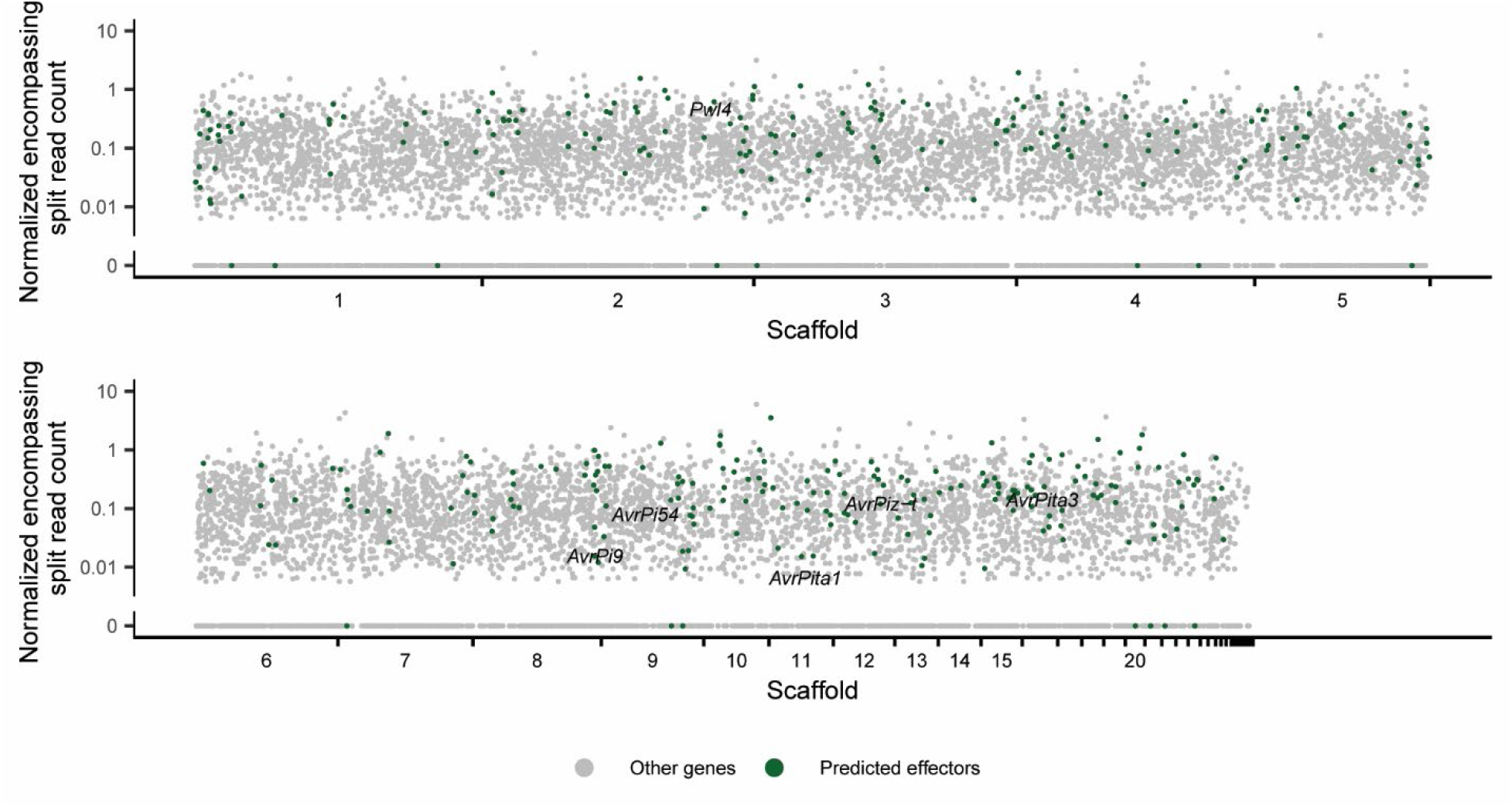
Effectors are enriched in eccDNAs in *M. oryzae*. Manhattan plot showing the number of encompassing split reads per million reads averaged across biological replicates for each gene in the *M. oryzae* Guy11 genome. Each dot represents one gene. Predicted effectors are shown in green and known effectors are shown as text.

## Discussion

EccDNAs have been shown to be a source of significant phenotypic [5,6,9,11,12] and genotypic [15, 48] plasticity that can help organisms adapt to stress. While eccDNAs have been extensively studied in human cancer [1], very few studies have attempted to study the circularome of other organisms, and even fewer have generated high quality whole circularome sequencing data. To expand our understanding of eccDNAs across the tree of life, we studied the circularome of the fungal plant pathogen *M. oryzae* and, through this analysis, developed many tools to analyze whole circularome sequencing data, which can often be difficult to interpret. These include a new pipeline to identify eccDNA forming regions and frameworks for comparing this data across organisms, identifying mechanisms of eccDNA formation of LTR retrotransposons, identifying gene sets enriched or depleted in eccDNAs, and identifying structural variants that may have been caused by eccDNAs. Our analysis also revealed that the circularome of *M. oryzae* contains a wide diversity of eccDNA forming regions that appeared to exceed those of other previously characterized organisms. This wide diversity likely contributed to under sequencing of our samples and a small overlap in exact eccDNA forming regions across samples. However, our analysis throughout this study showed that our samples clustered tightly together with regards to various features of the circularome, indicating that while exact eccDNA forming breakpoints were mostly not shared across samples, eccDNA formation hotspots were. We also found that eccDNA forming regions in *M. oryzae* were more commonly made up of LTR retrotransposons than other organisms. Though the results of our comparative analysis need to be verified using standardized protocols, these differences highlight the need to further characterize the circularome of other eukaryotes to obtain a better understanding of how they differ. Additionally, it is important to note that the data analyzed in this study only represent snapshots of the circularomes of the organisms described and could vary greatly across developmental stages and environmental stresses that were not included in these analyses. Further studies of eccDNAs across these different conditions are necessary to definitively describe and compare these molecules across organisms.

We analyzed the types of genes that were found on eccDNAs in *M. oryzae* and found that eccDNA-associated genes were often prone to presence-absence variation, hinting at a link between eccDNAs and genomic plasticity. However, we could not find direct evidence of gene deletions occurring through an eccDNA intermediate in *M. oryzae*. Similarly, we could not find any evidence of eccDNA-mediated translocations. These results could be due to our sequencing coverage and our bioinformatics pipelines not showing the full diversity of eccDNAs in *M. oryzae*. For example, our pipeline was unable to detect eccDNAs formed from HR between perfect repeats. Additionally, our scripts were able to identify an eccDNA-mediated translocation in wine yeasts but were limited to non-repetitive regions of the genome and may have missed some of these events in those regions in *M. oryzae*. Finally, it is possible that eccDNA-mediated translocations occur on a larger time scale than what we were able to sample within the *M. oryzae* species. However, it is likely that experimental approaches, including inducing the formation of specific eccDNAs, are necessary to determine whether these events lead to chromosomal deletions or rearrangements. On a genome-wide scale, single cell sequencing of the circularome as well as genomic DNA could also lead to a more precise view of eccDNA formation and structural variation as they occur in the cell during vegetative growth. These techniques will likely also need to be paired with amplification-free eccDNA sequencing protocols as well as high coverage, long read sequencing to fully resolve the structure of eccDNA molecules. Additionally, we found that eccDNA-associated genes presented characteristics associated with the gene-sparse, repeat-rich, and “fast” part of the plant pathogen genome where rapid adaptation to stress occurs [28]. The fact that eccDNA-associated genes were closer to repeats than other genes, but not transposons specifically, indicated that this effect was not simply caused by eccDNA formation by LTR retrotransposons. We also found that predicted effectors were enriched in eccDNA forming regions. These results show that eccDNA formation occurs in the same genomic contexts as rapid genome evolution in *M. oryzae* and could also point to eccDNAs directly playing a role in the plasticity of important genes like effectors.

We also identified a set of eccDNA-absent genes, which were never found fully encompassed by eccDNA forming regions under our experimental conditions. This observation was not explained by incomplete sequencing. Histone marks, expression and proximity to repetitive DNA did not appear to set these genes apart either. Though it is possible that other factors contribute to this phenomenon and directly prevent eccDNA formation in these regions, our data indicates that eccDNA formation in *M. oryzae* is not a random process and hints at selective pressure acting against cells that accumulate high copy numbers of these genes through eccDNA formation. This idea is supported by the absence of genes related to the myosin complex, which are deleterious at high copy numbers in other organisms.

Selective pressure during growth under stress could favor *M. oryzae* cells containing higher copy numbers of genes important for survival under these conditions as has been extensively shown in other organisms [5,6,9,11,12]. For example, we identified two genes associated with fungicide resistance in our eccDNA forming regions, which, if amplified, could lead to drug resistance, as previously observed [6,11,12]. Further experimentation and characterization of the *M. oryzae* circularome under stress is necessary to investigate if this eccDNA-mediated phenotypic plasticity is present in the plant pathogen. These experiments could also be used to assess how LTR retrotransposon activity changes in response to stress in *M. oryzae* and how the mechanisms of eccDNA formation that we described might be affected. We attempted to perform such experiments by sequencing *O. sativa* tissue infected by *M. oryzae* but found that *O. sativa* eccDNAs crowded out the circularome sequencing signal and prevented meaningful analysis, highlighting the need for a dedicated enrichment or single cell sequencing protocol. Additionally, analyzing the biological significance of the amplification of specific genes on eccDNAs, especially across treatments, may prove challenging and will require further tool development. For example, the same genes may be on eccDNAs of varying sizes and composition across samples. Multiple genes could also be on each eccDNA, further complicating the analysis. The complexity of eccDNAs combined with the limitations of current eccDNA sequencing techniques severely limits the analysis of circularome sequencing data, which is why we chose to focus our analysis on hotspots of eccDNA formation and groups of genes rather than individual genes. In the future, high coverage, long read sequencing of eccDNAs collected without amplification will likely be necessary to perform more thorough analyses of eccDNAs, and this type of study is likely to become the gold standard for the field once cost is no longer prohibitive.

## Conclusion

This study commences the characterization of the *M. oryzae* circularome and highlights its potential for generating phenotypic and genotypic plasticity. If eccDNAs were to facilitate these phenomena, they could become potential drug targets to prevent the rapid adaptation of the blast pathogen to environmental stress, fungicides, and resistant crop varieties. Furthermore, regions and genes prone to forming eccDNAs could be excluded as drug targets or as targets for engineered resistance in crops. On the other hand, we found 1,820 genes including several predicted effectors in the *M. oryzae* genome that were conserved in all other rice infecting isolates that we analyzed and that were in the eccDNA-absent group. These genes could be high potential targets for fungicide design or engineered resistance. Our study also describes the great diversity of eccDNAs and the enrichment of LTR retrotransposons in the *M. oryzae* circularome. These observations, in addition to the potential consequences of eccDNA formation, highlights the need to study these molecules in more organisms, including other fungal plant pathogens.

## Methods

### *M. oryzae* growth and DNA extraction

*M. oryzae* Guy11 was grown on Difco oatmeal agar plates for 21 days under constant light in a Percival Scientific Incubator Model CU-36L4 equipped with half fluorescent lights and half black lights. 1 cm^2^ of mycelium was scraped from the colony edge and used to start 3 liquid cultures (biological replicates) in 15 ml complete medium [49] (CM) in petri dishes. Liquid cultures were incubated without shaking for 3 days in the same growth chamber.

Total DNA extraction was performed according to a protocol from the Prof. Natalia Requena group at the Karlsruhe Institute of Technology. Briefly, mycelium grown in liquid culture was washed 3 times with water and then ground in liquid nitrogen. Ground mycelium was incubated in extraction buffer (0.1M Tris-HCl pH 7.5, 0.05 M EDTA, 1% SDS, 0.5 M NaCl) at 65°C for 30 minutes. 5M potassium acetate was then added to the samples which were then incubated on ice for 30 minutes. The supernatant was then washed with isopropanol and ethanol. Finally, the DNA pellet was resuspended in water and treated with RNase A (Thermo Scientific).

### *O. sativa* growth and DNA extraction

*O. sativa* samples were originally intended to serve as control samples to be compared to tissue infected by *M. oryzae* and therefore the methods below reflect this original intent. However, circularome sequencing data obtained from infected tissue was not included in this study as it included very little sequencing data that mapped to the *M. oryzae* Guy11 genome.

*O. sativa* cv. Nipponbare seeds were surface sterilized in 70% ethanol for 1 minute and 10% bleach for 10 minutes with thorough rinsing in sterile water after each before being placed on wet filter paper in a petri dish. The petri dish was wrapped in foil and placed at 4°C for 2 days to germinate. Germinated seedlings were planted in potting mix made up of 50% Turface and 50% Super Soil. Seedlings were grown for three weeks in a greenhouse under standard conditions. For three samples, the first true leaf was cut from one rice plant, its tip was removed, and it was then cut into two equal segments, approximately 10mm in length. This pair of segments was then placed on their abaxial surface on wet filter paper in a petri dish. Five hole-punches of filter paper soaked in 0.25% gelatin and 0.05% Tween-20 were then placed on each segment. The petri dishes were then placed in an airtight container with wet paper towels and placed on a windowsill for 7 days. Hole-punches were removed and non-chlorotic tissue in contact with hole-punches was ground in liquid nitrogen. DNA extraction was then performed using the Qiagen Plant DNeasy mini kit.

### Circular DNA enrichment

Total DNA obtained from DNA extractions (biological replicates) were then split into three samples (technical replicates) before circular DNA enrichment. This enrichment was performed according to a protocol from Lanciano *et al.* with a few modifications [18]. 5 µg of extracted DNA was used as input for circular DNA enrichment in *M. oryzae* and 750 ng of extracted DNA were used for *O. sativa*. To purify the samples and begin removing large linear DNA fragments, the samples were treated using a Zymo Research DNA Clean and Concentrator kit and standard protocols. Linear DNA digestion was then performed using Epicentre PlasmidSafe DNase and incubated at 37°C for 24 hours. DNase, ATP, and reaction buffer were then added to the samples every 24 hours while the incubation continued. In total, the reaction was allowed to proceed for 96 hours. Remaining DNA was then precipitated overnight at 4°C by adding 0.1 volume 3M sodium acetate, 2.5 volumes ethanol and 1 µl glycogen (20 mg/ml). Rolling circle amplification was then performed using the Illustra TempliPhi 100 Amplification Kit (GE Healthcare). Precipitated DNA was resuspended directly in 20 µl of the Illustra TempliPhi sample buffer and the amplification reaction was allowed to proceed for 24 hours at 30°C.

### Verification of circular DNA enrichment

In a separate experiment, 5 samples of *M. oryzae* mycelium were grown up in liquid culture and total DNA was extracted. Circular DNA enrichment was performed as before with some exceptions and without technical replicates. First, linear DNA digestion was only performed for 72 hours for 3 samples. Second, aliquots of the incubating samples were taken at 0 hours, 24 hours, 48 hours and 72 hours for these 3 samples, and 0 hours, 48 hours, 72 hours and 96 hours for the last 2 samples. qPCR was then used to verify linear DNA depletion in each sample using an Applied Biosystems QuantStudio 5 instrument and the QuantStudio Design and Analysis desktop software. Primers were used to amplify a portion of the *M. oryzae* actin gene (MGG_03982) along with Lightcycler 480 Sybr Green I master mix (Additional File 2: Table S3). Data from four qPCR technical replicates was obtained. Remaining linear DNA fraction in each sample at each timepoint was then calculated using the 2^-ΔΔCt^ method.

### Illumina library preparation and sequencing

Library preparation was performed by the QB3-Berkeley Functional Genomics Laboratory at UC Berkeley. DNA was fragmented with an S220 Focused-Ultrasonicator (Covaris), and libraries prepared using the KAPA Hyper Prep kit for DNA (Roche KK8504). Truncated universal stub adapters were ligated to DNA fragments, which were then extended via PCR using unique dual indexing primers into full length Illumina adapters. Library quality was checked on an Agilent Fragment Analyzer. Libraries were then transferred to the QB3-Berkeley Vincent J. Coates Genomics Sequencing Laboratory, also at UC Berkeley. Library molarity was measured via quantitative PCR with the KAPA Library Quantification Kit (Roche KK4824) on a BioRad CFX Connect thermal cycler. Libraries were then pooled by molarity and sequenced on an Illumina NovaSeq 6000 S4 flowcell for 2 x 150 cycles, targeting at least 10Gb per sample. FastQ files were generated and demultiplexed using Illumina bcl2fastq2 version 2.20 and default settings, on a server running CentOS Linux 7. One technical replicate did not pass quality control before library preparation and was omitted.

### PacBio library preparation and sequencing

Using a Covaris S220 Focused-Ultrasonicator, 2 ug of each DNA sample was sheared to an approximate fragment size of 5000 bp and purified using AMPure XP beads (Beckman Coulter). Library preparation was performed using the NEBNext Ultra DNA Library Prep Kit (kit number E7370L, New England Biolabs) and 8 cycles of PCR. Barcode sequences and barcodes assigned to each sample are described in Additional files 31 and 32. Libraries were then quality controlled using a Bioanalyzer high sensitivity DNA chip and the Agilent 2100 Bioanalyzer system. One technical replicate did not pass quality control before library preparation and was omitted. The samples were then submitted to Novogene (Tianjin, China) for PacBio sequencing which was performed on the PacBio Sequel platform using a 600-minute sequencing strategy and three SMRT cells.

### Inferring eccDNA forming regions from short read sequencing data

Illumina sequencing signal was analyzed using a custom pipeline inspired by previously published methods [13]. Illumina reads were first trimmed of Illumina TruSeq adapters using CutAdapt [50] version 2.4 with the nextseq-trim=20 option. Trimmed reads were then mapped to the *M. oryzae* Guy11 genome [37] and the 70-15 mitochondrial sequence [51] obtained from the Broad Institute (https://www.broadinstitute.org/scientific-community/science/projects/fungal-genome-initiative/magnaporthe-comparative-genomics-proj) using BWA-MEM [52] version 0.7.17-r1188 and the q and a options. Reads mapping to mitochondrial sequences were excluded. Uniquely mapped reads were then mined for split reads that mapped in the same orientation, had at least 20 bp of alignment on either side of the split, mapped to only two places in the genome, and where the start of the read mapped downstream from the end. This last filter sets these split reads apart from split reads that would indicate a deletion in the genome. Split reads for which one side of the split read mapped more than 50kbp away from the other or to a different scaffold than the other were excluded. Opposite facing read pairs were also obtained from uniquely mapped reads. Candidate eccDNA forming regions were then inferred by combining these two structural read variants. A split read that contained an opposite facing read pair that mapped no more than a combined 500 bp from the borders of the region contained within the two halves of the split read was considered a candidate eccDNA and a junction split read. The length distribution of these candidate eccDNA forming regions (Additional File 1: Fig. S35A) was then used to probabilistically infer candidate eccDNA forming regions from multi-mapping reads (Additional File 1: Fig. S35B). For each multi-mapping split read, a list of potential combinations of alignments that satisfied the previously described criteria for split reads was generated and one of these combinations was chosen at random, weighted by their length according to the generated length distribution. The chosen combinations were then used to infer additional candidate eccDNA forming regions by combining these with opposite facing read pairs as before, except this time obtained from unique and multi-mapping reads.

Each candidate eccDNA forming region was then validated by verifying that the region had over 95% read coverage and at least two junction split reads with the exact same coordinates. Candidate eccDNA forming regions that did not pass these criteria were considered low quality and were not included in the analysis.

### Inferring eccDNA forming regions from long read sequencing data

Circular consensus sequences (CCS) were first called from PacBio data using ccs version 3.4.1 (https://ccs.how/). Demultiplexing was then performed using lima version 1.9.0 (https://lima.how/) and sequences of barcodes used for library preparation (Additional Files 31 and 32). CCSs were then mapped to the *M. oryzae* Guy11 genome using minimap2 [53] version 2.18-r1015. Only uniquely mapped reads were kept for analysis. We then identified eccDNA forming regions by looking for split reads that either mapped to the same orientation to the same exact region multiple times or pairs of split alignments that were less than 50 kb apart, mapped in the same orientation and oriented properly so that they were indicative of a circular junction rather than a deletion.

### Outward PCR validation of eccDNA forming regions and PCR validation of eccDNA-absent genes

Outward facing primers were designed to 8 eccDNA forming regions of interest to validate their presence in our eccDNA sequencing samples. Primers were designed to amplify the junction of each eccDNA but not result in a product of the same size when used on genomic DNA (Additional File 2: Table S3). Primer3 [54] was used for primer design. The oligonucleutides were then synthesized by Integrated DNA technologies. PCR was performed using New England Biolab’s Phusion High-Fidelity DNA polymerase on *M. oryzae* Guy11 genomic DNA and rolling circle amplification products for the sample each eccDNA forming region was found in. 5ng DNA of each sample was used per 50 µl PCR reaction as well as 5X Phusion HF buffer, 10 mM dNTPs, 10 µM forward primer, 10 µM reverse primer and 1 unit of Phusion DNA polymerase. PCR conditions were as follows: initial denaturation at 98°C for 30 seconds, 35 cycles of denaturation at 98°C for 10 seconds, annealing at 64°C or 65°C for 30 seconds, extension at 72°C for 10 seconds, and a final extension at 72°C for 5 minutes. PCR products were run on a 2% agarose gel to check for amplification. Bands of the expected size were extracted from electrophoresis gels using Zymo Research’s Zymoclean Gel DNA Recovery Kit. Sanger sequencing was performed by the UC Berkeley DNA Sequencing Facility and Sanger sequences were examined for matches to corresponding eccDNA forming regions using BLASTN [55] version 2.2.9 and manual inspection.

PCR validation of eccDNA-absent genes was performed using similar methods. Primers were designed to amplify the entire annotated gene region of *MYO1* and the actin gene (MGG_03982) and a small segment of the *MAGGY* LTR retrotransposon from genomic DNA. 2ng DNA of each sample was used per 20 µl PCR reaction as well as 5X Phusion HF buffer, 10 mM dNTPs, 10 µM forward primer, 10 µM reverse primer and 0.4 units of Phusion DNA polymerase. PCR conditions were as follows: initial denaturation at 98°C for 30 seconds, 25 cycles of denaturation at 98°C for 10 seconds, annealing at 64°C or 65°C for 30 seconds, extension at 72°C for 5, 60 or 120 seconds, and a final extension at 72°C for 5 minutes. PCR products were run on a 1% agarose gel to check for amplification.

### Comparing eccDNA forming regions inferred from Illumina data and eccDNA forming regions inferred from PacBio data

EccDNA forming regions called using Illumina data and PacBio data were found to be identical if their start and end coordinates were within 10 bp of each other to account for mapping errors. EccDNA forming regions were then called with less stringent requirements to verify if any of the missing eccDNA forming regions were being filtered out somewhere in the pipeline. In this test, all uniquely mapped split reads that had 10 or more bp overlap on either side, were properly oriented, and were less than 50kb apart were considered eccDNA forming regions.

### Benchmarking eccDNA forming regions called using our pipeline on previously published data

EccDNA forming regions called using our pipeline were compared to eccDNA forming regions previously published for *H. sapiens* [13]. EccDNA forming regions were found to be identical if their start and end coordinates were within 10 bp of each other. EccDNA forming regions described as low quality by the authors were excluded from the published dataset before comparison. High coverage eccDNA forming regions were chosen for comparison if they had more than 10 associated junction split reads. Finally, multi-mapping reads were excluded from the pipeline to identify eccDNA forming regions called using only uniquely mapped reads.

### Comparing eccDNA sequencing samples to each other

Overlaps in eccDNA forming regions between samples were first calculated based off the exact coordinates of the eccDNA forming regions and Venn diagrams based off these overlaps were generated using the ggVennDiagram R package [56] version 1.2.0. EccDNA forming regions found in all technical replicates taken from each biological replicate were first combined before looking for overlaps between biological replicates. Overlaps were then calculated with various levels of tolerance for the start and end coordinates of the eccDNA forming regions so that regions in one sample that were within 10, 100, or 1000 bp from the start and end coordinates of a region in another sample were considered to be found in both samples. Rarefaction analysis for eccDNA forming regions in all samples was performed by sampling mapped eccDNA sequencing reads at random in increasing 10% intervals. For each subsample, eccDNA forming regions were called as previously described and counted. Principal component analysis of read coverage was performed by first calculating junction split read coverage for all 10kbp windows in the genome for each sample. These values were then normalized to the total number of junction split reads in each sample. The matrix of normalized junction split read coverage for all samples was then processed using the prcomp function in R version 3.6.1 with the scale = TRUE option, and the first 6 principal components were plotted using the ggbiplot R package [57] version 0.55.

### Gene and effector annotation

The *M. oryzae* Guy11 genome along with 162 other rice-infecting *M. oryzae* genomes (Additional File 25) were annotated using the FunGAP [58] version 1.1.0 annotation pipeline. For all genomes, RNAseq data (SRR8842990) obtained from GEO accession GSE129291 was used along with the proteomes of *M. oryzae* 70-15, P131 and MZ5-1-6 taken from GenBank (accessions GCA_000002495.2, GCA_000292605.1, and GCA_004346965.1, respectively). The ‘sordariomycetes_odb10’ option was used for the busco_dataset option and the ‘magnaporthe_grisea’ option was used for the augustus_species option. For repeat masking, a transposable element library generated by combining the RepBase [59] fngrep version 25.10 with a *de novo* repeat library generated by RepeatModeler [60] version 2.0.1 run on the *M. oryzae* Guy11 genome with the LTRStruct option was used for all genomes. Genes in *M*. *oryzae* Guy11 were assigned names according to the gene names listed on UniProtKB for *M. oryzae* 70-15 accessed in October 2021. To make this assignment, *M. oryzae* Guy11 proteins were aligned to the *M. oryzae* 70-15 proteome using BLASTP [55] version 2.7.1+ and hits with greater than 80% sequence identity and that spanned more than 80% of the length of both the *M. oryzae* Guy11 protein and the *M. oryzae* 70-15 protein were assigned names.

Effectors were predicted among *M. oryzae* Guy11 genes by first selecting genes with signal peptides which were predicted using SignalP [61] version 5.0b Darwin x86_64. Genes with predicted transmembrane domains from TMHMM [62] version 2.0c were then excluded. Finally, EffectorP [63] version 2.0 was used to predict effectors from this secreted gene set. Previously well-characterized effectors were identified using previously published protein sequences [64] and DIAMOND [65] version 2.0.9.147.

### High quality LTR-retrotransposon annotations in *M. oryzae*

High quality, full length, consensus sequences for known *Gypsy* elements in *M. oryzae* (*MAGGY*, *GYMAG1*, *GYMAG2*, *PYRET*, *MGRL3*) and one *Copia* element (*Copia1*) were generated using the WICKERsoft [66] suite of tools. Reference sequences from other genomes for each element were obtained from the RepBase [59] fngrep version 25.10 library. The *M. oryzae* Guy11 genome was then scanned for the presence of these sequences using BLASTN [55] version 2.2.9 and then filtered to hits with 90% sequence identity and that contained 90% of the sequence length. Hits for each reference sequence were then extended to include 500 base pairs of genomic sequence upstream and downstream of the hit. A multiple sequence alignment of hits for each reference sequence was then generated using ClustalW [67] version 1.83 and boundaries were visually inspected and trimmed.

Consensus sequences for each element were then generated from these multiple sequence alignments. These consensus sequences were split into LTR and internal regions by self-alignment using the BLASTN [55] webserver in August 2020 to identify LTRs. These consensus sequences are available in Additional File 33. Finally, the locations of these elements in *M. oryzae* Guy11 genome were annotated with RepeatMasker [68] version 4.1.1 with the -cutoff 250, -nolow, -no_is, and -norna options to identify their locations in the *M. oryzae* Guy11 genome. For read coverage plots as well as histone and GC content plots, full length LTR retrotransposon copies were required. These were identified by using the original full length consensus sequences with RepeatMasker as before and then filtering to hits greater than 3000 bp in length and greater than 90% sequence identity.

### Comparative analysis of eccDNA forming regions

Analysis of eccDNA forming regions in organisms other than *M. oryzae* were performed as described above for Illumina sequencing data using previously published genome, gene annotation and transposable element annotation files (Additional File 34). However, unlike the other data used in this study, the sequencing data in the *S. cerevisiae* dataset was single-end and therefore opposite facing read pairs could not be used to infer eccDNA forming regions. Instead, only eccDNA forming regions with three overlapping junction split reads were used for analysis. For all organisms, reads mapping to unplaced scaffolds and organellar genomes were removed after mapping as described above for the *M. oryzae* mitochondrial genome. These scaffolds were also removed from genome size, number of coding base pairs, and number of LTR retrotransposon base pairs calculations for comparative analysis. To calculate the percent of the genome that was covered in each sample, the genomecov command of the BEDtools [69] suite versions 2.28.0 was used with the -d option along with the coordinates of eccDNA forming regions for each sample. Any base pair with a coverage value greater than zero was counted as being a portion of the genome in an eccDNA forming region.

### Characterization of eccDNA formation by LTR retrotransposons

To generate the Manhattan plot, junction split reads were filtered by selecting regions that were made up of 90% LTR retrotransposon sequences. Junction split read coverage was then calculated for each 100 bp window in the genome. Coverage values were then normalized to the total number of LTR eccDNA junction split reads per sample. These coverage values were then averaged across technical replicates for each biological replicates, and then averaged across biological replicates. Finally, only 100 bp bins that overlapped at least 50 bp with an LTR retrotransposon were plotted in Fig 3A. For Additional File 1: Fig. S10, only bins with coverage greater than 0 were plotted.

To simulate expected read coverage for different types of LTR eccDNAs, the *Copia1* consensus sequence was taken as a reference, though the *MAGGY* consensus sequence yielded identical results. Simulated DNA sequences were then generated for each type of LTR eccDNA. The expected 2-LTR circular sequence generated by NHEJ (scenario 1, Fig. 4A) was simply made up of two LTR sequences and the internal sequence, and the expected 1-LTR circle sequence generated by HR (scenario 3, Fig. 4C) was made up of one LTR sequence and the internal sequence. These sequences were shuffled 1000 times to generate 1000 sequences starting at various points of the expected circularized sequence. For the 1-LTR circle sequence generated by autointegration (scenario 2, Fig. 4B), the random autointegration events were simulated by choosing a random length segment of the internal sequence starting with its start or end, adding the LTR sequence to this sequence, and randomly shuffling the sequence to simulate a circular sequence. This process was repeated 1000 times to generate 1000 sequences. Finally, for each scenario, Illumina reads were simulated to reach 2000x coverage for each of the simulated sequences using ART Illumina [70] version 4.5.8 and the following parameters: 150 bp read length, 450 bp mean insert size, 50 bp insert size standard deviation, HiSeqX TruSeq. Reads were mapped to the simulated sequences using BWA-MEM [52] version 0.7.17-r1188 with default settings and coverage for each base pair was calculated.

To generate observed coverage for each element, sequencing read coverage across the genome was calculated for all 10 base pair windows in the *M. oryzae* Guy11 genome for each sample. Coverage values were then normalized to the total number of mapped sequencing reads in each sample. These coverage values were then averaged across technical replicates for each biological replicates, and then averaged across biological replicates. Finally, profile plot data was generated for full length, high confidence sequences for each LTR retrotransposon using computeMatrix scale-regions and plotProfile of the DeepTools [71] suite of tools version 3.5.1 using full length, high confidence LTR retrotransposon sequences. Profile plots were also generated using previously published whole genome sequencing data by averaging sequencing coverage across all three samples [32,37,38].

### Identification of split reads associated with eccDNA formation from LTR retrotransposons

Split reads were first identified as any read that mapped to only two places in the genome with at least 20 base pairs of alignment on either side. LTR-LTR split reads were then selected from these split reads for each LTR retrotransposon if both sides of the split read had any overlap with any copy of that retrotransposon’s LTR in the genome. LTR-internal split reads were selected if one side of the split read had any overlap with any copy of the retrotransposon’s LTR in the genome and the other side had any overlap with any copy of the retrotransposon’s internal region in the genome. Read coverage, LTR-LTR split read coverage, and LTR-internal coverage was then calculated for each annotation of each LTR retrotransposon. Coverage values were then normalized to the total number of mapped sequencing reads in each sample. These coverage values were then averaged across technical replicates for each biological replicates, and then averaged across biological replicates.

### Comparison of microDNAs and large eccDNAs across organisms

Genome, gene annotation, and transposable element annotation files for each organism used for this analysis were as previously described (Additional File 34). Again, organellar genomes as well as unplaced contigs were filtered out of these files before analysis. Introns and UTRs were added to gene annotation files that were missing these elements using the ‘agat_convert_sp_gff2gtf.pl’ and ‘agat_sp_add_introns.pl’ commands from the AGAT toolkit version 0.6.2 (https://github.com/NBISweden/AGAT). Cpgplot of EMBOSS [72] version 6.6.0.0 was used to annotate CpG islands in each genome. Upstream and downstream regions were defined as being 2000 base pairs upstream from the transcription start site and downstream from the transcription end site, respectively. Genic regions were defined as being made up of all sequences between transcription start and end sites and intergenic regions were the opposite. Junction split reads were counted as being from a specific region if they overlapped to any extent within that region.

The observed percentage of junction split reads overlapping with each region type was calculated for each sample for each organism and an average of these percentages was calculated. The junction split reads of each sample were then shuffled across the genome 10 times, excluding LTR retrotransposon locations, and an expected percentage for each region was calculated, averaged across all permutations, then averaged across all samples for each organism. Finally, the log_2_ of the fold enrichment was calculated by taking the log_2_ of the observed average percentage over the expected average percentage.

### Correlation of expression and eccDNA formation

Previously published RNAseq data from *M. oryzae* Guy11 grown in liquid culture in rich medium was obtained [73] (Additional File 35). The data was mapped to the *M. oryzae* Guy11 genome using STAR [74] version 2.7.1a with the quantMode GeneCounts option. Read counts per gene were then divided by library size and multiplied by the length of each gene in order to obtain reads per kilobase million (RPKMs). RPKMs per gene were then averaged across all samples.

Junction split read counts per gene used to analyze the correlation of expression and eccDNA formation were generated for each gene by counting the number of junction split reads that intersect the gene to any extent. Counts per gene were first assessed for each sample and normalized to the number of junction split reads in that sample. Normalized counts were then averaged across technical replicates for each biological replicate. Average counts per biological replicate were then averaged to obtain the final result.

To compare gene content and eccDNA formation, the *M. oryzae* genome was divided into 100kbp bins and the number of genes per bin was then calculated. Junction split reads per bin were calculated for each sample using the same method. Junction split read per bin values were then normalized to the total number of junction split reads in each sample. These values were then averaged across technical replicates for each biological replicate, and then averaged across biological replicates.

### ARS consensus sequence enrichment analysis

The published ACS sequence profile [42] was used to identify ACSs in eccDNA forming regions using the FIMO [75] software version 4.12.0. Only hits scoring greater than 17 were kept. In order to test for enrichment of these sequences, an expected distribution of ACS sequences was generated by randomly shuffling eccDNA forming regions across the *M. oryzae* Guy11 genome, excluding regions containing LTR retrotransposons. The observed number of ACS sequences in eccDNA forming regions was then compared to the expected distribution to generate a p-value.

### Histone mark and GC content profile plots

Previously published ChIPSeq data for H3K27me3, H3K27ac, H3K36me3 and loading controls was obtained [73]. Sequencing reads for each technical replicate were combined before reads for each treatment for each biological replicate were mapped to the *M. oryzae* Guy11 genome using BWA-MEM [52] version 0.7.17-r1188 with default settings. The bamCompare command from the DeepTools [71] suite of tools version 3.5.1 with the scaleFactorsMethod readCount option was then used to compare the signal from each treatment to the loading control for each biological replicate. computeMatrix scale-regions was then used in conjunction with the plotProfile command to generate processed data for profile plots. After verifying that all biological replicates resulted in similar profile plots, only the first biological replicate was chosen for presentation.

To generate tracks used for profile plots, a few different strategies were used. GC content profile plots were generated by calculating GC percentage for 50 base pair windows throughout the genome. Profile plot data was then generated using computeMatrix scale-regions and plotProfile commands as before. Methylated and acetylated genes were determined using the methylation and acetylation peaks published by Zhang *et al.* [73]. Marked genes were called when at least 50% of the gene overlapped with a peak. Large eccDNAs, microDNAs, and LTR-eccDNAs from all *M. oryzae* Guy11 samples were combined into a single list which was filtered for duplicates and used for the corresponding tracks in the profile plots. The genome baseline track was generated by combining all of these eccDNA forming regions and shuffling them randomly across the genome. Finally, the full length, high quality LTR-retrotransposon annotations described above were used for LTR retrotransposon tracks. The same approach was used for generating profile plots to compare histone marks and GC content for eccDNA-associated and eccDNA-absent genes.

### Identification of eccDNA-associated and eccDNA-absent genes

Encompassing split read counts per gene for determining eccDNA-associated and eccDNA-absent genes were generated for each gene by counting the junction split reads that fully encompass the gene using the intersect command of the BEDTools [69] suite version 2.28.0 with the -f 1 option. This count was normalized to the total number of junction split reads in each sample, then averaged across technical replicates for each biological replicate. Genes with a count of zero were removed from each biological replicate before being sorted by this count. Genes in the top third for this count were compared between biological reps using the ggVennDiagram R package [56] version 1.2.0. This count was averaged across biological replicates to obtain the encompassing split read count per gene for visualizations in Fig. 5 and Fig. 8 and for comparison between predicted effectors and other genes (Additional File 1: Fig. S34).

### GO enrichment analysis

GO terms were first assigned to annotated *M. oryzae* Guy11 genes using the PANNZER2 [76] webserver on August 17^th^, 2020. Annotated GO terms were then filtered to annotations with a positive predictive value greater than 0.6. The topGO [77] R package version 2.36.0 was then used to parse assigned GO terms and reduce the gene list to a list of feasible genes for analysis. Either eccDNA-associated or eccDNA-absent were assigned as significant genes, and the number of these genes belonging to each GO term was used as the observed value for the enrichment analysis. A kernel density function was then generated using the gene lengths of the significant gene set. The same number of genes as the significant gene set were then sampled at random from the feasible gene set using weighted random selection with weights obtained from the kernel density function. This random sampling was repeated 100 times and the average of the number of genes belonging to each GO term was used as the expected value for the enrichment analysis. Finally, the Chi-square statistic was then computed comparing observed and expected values to test for enrichment or depletion of each GO term.

### Gene presence absence variation

In order to identify genes prone to presence absence variation in the *M. oryzae* Guy11 genome, OrthoFinder [78] version 2.5.1 with default settings was used on all of the *M. oryzae* proteomes and the *Neurospora crassa* proteome obtained from GenBank (accession GCA_000182925.2). Then, for each *M. oryzae* genome, we queried whether each gene annotated in the *M. oryzae* Guy11 genome had an ortholog identified by OrthoFinder in that genome. Finally, the absence of genes without orthologues was confirmed using BLASTN [55] version 2.7.1+.

Small, genic deletions were identified using orthologs identified by OrthoFinder [78] version 2.5.1 as before. For each genome, we looked for genes in the *M. oryzae* Guy11 genome that had no ortholog in that genome, but that were flanked by two genes with orthologs in that genome. One-to-many, many-to-many, and many-to-one orthologs were excluded from this analysis. Candidate gene deletions were validated using alignments performed using the nucmer and mummerplot commands of the MUMmer [79] suite of tools version 4.0.0rc1 to verify that a DNA deletion truly existed, and that this deletion overlapped the gene of interest.

### Identification of eccDNA-mediated translocations

Identification of translocations with a potential eccDNA intermediate was done by first aligning two genomes using the nucmer command of the MUMmer [79] suite of tools version 4.0.0rc1 with the maxmatch option. The nucmer output was then parsed to look for portions of the reference genome that had an upstream region that aligned to one query scaffold, followed by two separate adjacent alignments to another query scaffold, followed by a downstream region that aligned to the original query scaffold. We also required that the two adjacent alignments in the center of the region were to adjacent regions in the query scaffold but their order was reversed compared to the reference. Candidate eccDNA-mediated translocations were verified manually by inspecting alignment plots generated using the mummerplot command. The *S. cerevisiae* EC1118 (GCA_000218975.1) and M22 genomes (GCA_000182075.2) obtained from GenBank were used to verify the ability of our pipeline to detect these translocation events. The *M. oryzae* Guy11 genome was then compared to 306 *M. oryzae* genomes (Additional File 27) to look for these events in the *M. oryzae* species. Before alignment, transposable elements were masked from these *M. oryzae* genomes using RepeatMasker [68] version 4.1.1 with the -cutoff 250, -nolow, -no_is, and -norna options, as well as a transposable elements library generated by combining the RepBase [59] fngrep version 25.10 with the de novo repeat library generated by RepeatModeler [60] version 2.0.1 run on the *M. oryzae* Guy11 genome with default settings aside from the LTRStruct argument.

### Minichromosome genes and eccDNAs

Scaffolds corresponding to minichromosomes in the *M. oryzae* FR13 (GCA_900474655.3), CD156 (GCA_900474475.3), and US71 (GCA_900474175.3) genomes were extracted according to previously published data [80]. Exonerate [81] version 2.4.0 was then used with the protein2genome model to identify genes in the *M. oryzae* Guy11 genome that were found on minichromosomes in these other isolates. Hits with greater than 70% sequence identity to any minichromosome scaffold were identified as genes found on minichromosomes. Encompassing split reads were then counted for all genes. This count was normalized to total number of junction split reads in each sample, then averaged across technical replicates for each biological replicate, then averaged across biological replicates. Finally, normalized encompassing split read counts for genes found on minichromosomes were compared to genes not found on minichromosomes.

### Rarefaction analysis for eccDNA-absent genes and unique eccDNA forming regions

Rarefaction analysis for genes found fully encompassed by eccDNA forming regions were performed by first sampling eccDNA forming regions from all samples at random in increasing 10% intervals. For each subsample, the number of genes found fully encompassed by eccDNA forming regions was determined as before. Next, eccDNA forming regions were shuffled across the genome and sampled at random in increasing 10% intervals. Again, the number of genes found fully encompassed by eccDNA forming regions was determined for each sample. This analysis was performed 100 times with similar results as those represented in Fig. 5C. A similar approach was used for rarefaction analysis of eccDNA forming regions but the number of unique microDNAs, large eccDNAs and LTR-eccDNAs were counted at each subsample instead.

### Data processing and analysis

Data processing was performed in a RedHat Enterprise Linux environment with GNU bash version 4.2.46(20)-release. GNU coreutils version 8.22, GNU grep version 2.20, GNU sed version 4.2.2, gzip version 1.5, and GNU awk version 4.0.2 were all used for file processing and handling. Conda version 4.8.2 (https://docs.conda.io/en/latest/) was used to facilitate installation of software and packages. Code parallelization was performed with GNU parallel [82] version 20180322. Previously published data was downloaded using curl version 7.65.3 (https://curl.se/) and sra-tools version 2.10.4 (https://github.com/ncbi/sra-tools). Image file processing was performed with the help of ghostscript version 9.25 (https://ghostscript.com/) and imagemagick version 7.0.4-7 (https://imagemagick.org/index.php). BED format files were processed using bedtools [69] version 2.28.0 and bedGraphToBigWig version 4 (https://www.encodeproject.org/software/bedgraphtobigwig/). SAM and BAM format files were processed with SAMtools [83] version 1.8 and Picard version 2.9.0 (https://broadinstitute.github.io/picard/).

Data processing was also facilitated by custom Python scripts written in Python version 3.7.4 with the help of the pandas [84] version 0.25.1 and numpy [85] version 1.17.2 modules. The scipy [86] version 1.4.1 and more-intertools version 7.2.0 (https://more-itertools.readthedocs.io/) modules were also used.

Data analysis and statistical analyses were performed in R version 3.6.1. Data handling was processed using data.table [87] version 1.13.6, tidyr [88] version 1.1.3, reshape2 [89] version 1.4.4, and dplyr [90] version 1.0.4 packages. Plotting was performed using the ggplot2 [91] version 3.3.5 package, with help from RColorBrewer [92] version 1.1.2, scales [93] version 1.1.1, cowplot [94] version 1.1.1, ggprepel [95] version 0.9.1 and ggpubr [96] version 0.4.0 packages. The Gviz [97] version 1.28.3 was used for BAM file visualization. Tables were made using gt [98] version 0.3.1.

## Declarations

### Availability of Data and Materials

The datasets supporting the conclusions of this article are available in the NCBI’s Sequence Read Archive repository. Illumina circularome sequencing data for *M. oryzae* was submitted under BioProject accession PRJNA768097. PacBio circularome sequencing data for *M. oryzae* was submitted under BioProject accession PRJNA556909. Illumina circularome sequencing data for *O. sativa* was submitted under BioProject accession PRJNA768410. Additional datasets supporting the conclusions of this article are available on Zenodo. Genomes and annotation files used for comparative circularome are available under the DOI 10.5281/zenodo.5544950. Annotated genes and predicted proteins for rice-infecting *M. oryzae* isolates are also available under the DOI 10.5281/zenodo.5542597. Outputs from OrthoFinder2 run on rice-infecting *M. oryzae* proteomes are also available under the DOI 10.5281/zenodo.5544260. Finally, all files used for statistical analysis and plotting are available under the DOI 10.5281/zenodo.7114261.

Code for the pipeline used to call eccDNA forming regions for Illumina sequencing data is available in a maintained GitHub repository (https://github.com/pierrj/ecc_caller). All other code used for raw data processing, data analysis, and figure generation is available in a GitHub repository (https://github.com/pierrj/moryzae_eccdnas_manuscript_code_final).

### Authors’ contributions

PMJ and KVK conceptualized and designed the study. PMJ collected and analyzed the data. PMJ wrote the original draft manuscript. PMJ and KVK reviewed and edited the manuscript. Both authors read and approved the final manuscript.

## Acknowledgements

We thank Snighda Poddar for providing the *M. oryzae* Guy11 isolate and for advice for culturing the pathogen. We thank Ursula Oggenfuss for advice on using WICKERsoft for generating LTR retrotransposon consensus sequences. We also thank the Krasileva lab for feedback on manuscript preparation. This research used the Savio computational cluster resource provided by the Berkeley Research Computing program at the University of California, Berkeley (supported by the UC Berkeley Chancellor, Vice Chancellor for Research, and Chief Information Officer). We also thank Novogene (Tianjin, China) for technical support.

## Supplementary information

Additional File 1: Supplementary Figures.

**Fig. S1.** Degradation of linear DNA using exonuclease treatment.

**Fig. S2.** Outward PCR validation of eccDNA forming regions.

**Fig. S3.** Overlap in exact break points of eccDNA forming regions across samples.

**Fig. S4.** Rarefaction analysis of sequencing coverage and eccDNA forming regions across all samples.

**Fig. S5.** Principal components analysis of sequencing coverage between samples.

**Fig. S6.** Overlap in eccDNA forming regions across samples, with increasing tolerance for start and end coordinates.

**Fig. S7.** Overlap between eccDNA forming regions called using PacBio sequencing data and Illumina sequencing data.

**Fig. S8.** Comparison between eccDNA forming regions in human samples called in this manuscript and in the original publication.

**Fig. S9.** Comparison of eccDNA forming regions between *M. oryzae* and other previously studied organisms.

**Fig. S10.** EccDNA forming regions composed of more than 90% LTR retrotransposon sequence in *M. oryzae*.

**Fig. S11.** Percentage of the *M. oryzae* Guy11 genome made up of each LTR retrotransposon.

**Fig. S12.** Correlation between number of LTR-LTR split reads and sequencing reads in eccDNA sequencing samples for each LTR retrotransposon in *M. oryzae*.

**Fig. S13.** Correlation between number of LTR-internal split reads and sequencing reads in eccDNA sequencing samples for each LTR-retrotransposon in *M. oryzae*.

**Fig. S14.** Number of LTR-LTR split reads and LTR-internal split reads in eccDNA sequencing samples for each LTR retrotransposon in *M. oryzae*.

**Fig. S15.** Expected read coverage for LTR retrotransposons in *M. oryzae*.

**Fig. S16.** MicroDNA enrichment and depletion in the genomes of various organisms.

**Fig. S17.** Enrichment and depletion of microDNAs and large eccDNAs across various genomic regions in *M. oryzae*.

**Fig. S18.** Correlation between gene count and junction split read count across the *M. oryzae* genome.

**Fig. S19.** Correlation between junction split read count and expression for *M. oryzae* genes.

**Fig. S20.** Comparison of junction split read counts between eccDNA forming regions with and without an ACS.

**Fig. S21.** GC content and chromatin marks of eccDNA forming regions in *M. oryzae*.

**Fig. S22.** Overlap between genes enriched on eccDNAs in biological replicates.

**Fig. S23.** GO terms associated with eccDNA-associated genes.

**Fig. S24.** GC content and chromatin marks of eccDNA-associated and eccDNA-absent genes in *M. oryzae*.

**Fig. S25.** Comparison of expression data between eccDNA-associated genes and eccDNA-absent genes in *M. oryzae*.

**Fig. S26.** Proximity of *M. oryzae* genes to repeats.

**Fig. S27.** Proximity of *M. oryzae* genes to TEs.

**Fig. S28.** Predicted effectors are prone to presence-absence variation in *M. oryzae*.

**Fig. S29.** Rarefaction curves for eccDNA forming regions in *M. oryzae*.

**Fig. S30.** Example of an eccDNA-mediated translocation in wine yeasts.

**Fig. S31.** Comparison of encompassing split read counts between genes found on mini-chromosomes in *M. oryzae* and other genes.

**Fig. S32.** GO terms associated with eccDNA-absent genes.

**Fig. S33.** PCR validation of eccDNA-absent genes.

**Fig. S34.** Effectors are enriched in eccDNAs in M. oryzae.

**Fig. S35.** Lengths of eccDNA forming regions in *M. oryzae*.

Additional File 2: Supplementary Tables.

**Table S1.** Number of eccDNA forming regions called using whole genome sequencing data.

**Table S2.** Summary of protocols used to extract eccDNAs in studies analyzed in this manuscript.

**Table S3.** Primers used for qPCR validation of linear DNA degradation and outward PCR validation of eccDNA forming regions.

Additional File 3: List of eccDNA forming regions called using Illumina circularome sequencing data for *M. oryzae* in this study.

The first column describes the sample the eccDNA forming region was called with, the next three columns represent the genomic coordinates of the eccDNA forming region, and the last column represents the number of junction split reads used to call the eccDNA forming region.

Additional File 4: List of eccDNA forming regions called using PacBio circularome sequencing data for *M. oryzae* in this study.

The first column describes the sample the eccDNA forming region was called with, the next three columns represent the genomic coordinates of the eccDNA forming region, and the last column represents the number of junction split reads used to call the eccDNA forming region.

Additional File 5: List of eccDNA forming regions called using Illumina circularome sequencing data for *H. sapiens* muscle tissue published by Møller *et al.* [13].

The first column describes the sample the eccDNA was called with, the next three columns represent the genomic coordinates of the eccDNA forming region, and the last column represents the number of junction split reads used to call the eccDNA forming region.

Additional File 6: List of eccDNA forming regions called using Illumina circularome sequencing data for *H. sapiens* leukocytes published by Møller *et al.* [13].

The first column describes the sample the eccDNA was called with, the next three columns represent the genomic coordinates of the eccDNA forming region, and the last column represents the number of junction split reads used to call the eccDNA forming region.

Additional File 7: List of eccDNA forming regions called using Illumina circularome sequencing data for *O. sativa* in this study.

The first column describes the sample the eccDNA forming region was called with, the next three columns represent the genomic coordinates of the eccDNA forming region, and the last column represents the number of junction split reads used to call the eccDNA forming region.

Additional File 8: List of eccDNA forming regions called using Illumina circularome sequencing data for *O. sativa* leaf tissue published by Lanciano *et al.* [18].

The first column describes the sample the eccDNA was called with, the next three columns represent the genomic coordinates of the eccDNA forming region, and the last column represents the number of junction split reads used to call the eccDNA forming region.

Additional File 9: List of eccDNA forming regions called using Illumina circularome sequencing data for *O. sativa* seed tissue published by Lanciano *et al.* [18].

The first column describes the sample the eccDNA was called with, the next three columns represent the genomic coordinates of the eccDNA forming region, and the last column represents the number of junction split reads used to call the eccDNA forming region.

Additional File 10: List of eccDNA forming regions called using Illumina circularome sequencing data for *O. sativa* callus tissue published by Lanciano *et al.* [18].

The first column describes the sample the eccDNA was called with, the next three columns represent the genomic coordinates of the eccDNA forming region, and the last column represents the number of junction split reads used to call the eccDNA forming region.

Additional File 11: List of eccDNA forming regions called using Illumina circularome sequencing data for *A. thaliana* WT tissue published by Lanciano *et al.* [18].

The first column describes the sample the eccDNA was called with, the next three columns represent the genomic coordinates of the eccDNA forming region, and the last column represents the number of junction split reads used to call the eccDNA forming region.

Additional File 12: List of eccDNA forming regions called using Illumina circularome sequencing data for *A. thaliana epi12* mutant tissue published by Lanciano *et al.* [18].

The first column describes the sample the eccDNA was called with, the next three columns represent the genomic coordinates of the eccDNA forming region, and the last column represents the number of junction split reads used to call the eccDNA forming region.

Additional File 13: List of eccDNA forming regions called using Illumina circularome sequencing data for *A. thaliana* leaf tissue published by Wang *et al.* [17].

The first column describes the sample the eccDNA was called with, the next three columns represent the genomic coordinates of the eccDNA forming region, and the last column represents the number of junction split reads used to call the eccDNA forming region.

Additional File 14: List of eccDNA forming regions called using Illumina circularome sequencing data for *A. thaliana* root tissue published by Wang *et al.* [17].

The first column describes the sample the eccDNA was called with, the next three columns represent the genomic coordinates of the eccDNA forming region, and the last column represents the number of junction split reads used to call the eccDNA forming region.

Additional File 15: List of eccDNA forming regions called using Illumina circularome sequencing data for *A. thaliana* stem tissue published by Wang *et al.* [17].

The first column describes the sample the eccDNA was called with, the next three columns represent the genomic coordinates of the eccDNA forming region, and the last column represents the number of junction split reads used to call the eccDNA forming region.

Additional File 16: List of eccDNA forming regions called using Illumina circularome sequencing data for *A. thaliana* flower tissue published by Wang *et al.* [17].

The first column describes the sample the eccDNA was called with, the next three columns represent the genomic coordinates of the eccDNA forming region, and the last column represents the number of junction split reads used to call the eccDNA forming region.

Additional File 17: List of eccDNA forming regions called using Illumina circularome sequencing data for *S. cerevisiae* WT cells published by Møller *et al.* [13].

The first column describes the sample the eccDNA was called with, the next three columns represent the genomic coordinates of the eccDNA forming region, and the last column represents the number of junction split reads used to call the eccDNA forming region.

Additional File 18: List of eccDNA forming regions called using Illumina circularome sequencing data for *S. cerevisiae* GAP1^circle^ cells published by Møller *et al.* [19].

The first column describes the sample the eccDNA was called with, the next three columns represent the genomic coordinates of the eccDNA forming region, and the last column represents the number of junction split reads used to call the eccDNA forming region.

Additional File 19: List of eccDNA forming regions called using Illumina circularome sequencing data for *S. cerevisiae* cells from the deletion collection published by Møller *et al.* [19].

The first column describes the sample the eccDNA was called with, the next three columns represent the genomic coordinates of the eccDNA forming region, and the last column represents the number of junction split reads used to call the eccDNA forming region.

Additional File 20: List of eccDNA forming regions called using Illumina circularome sequencing data for *S. cerevisiae* cells from the deletion collection treated with zeocin published by Møller *et al.* [19].

The first column describes the sample the eccDNA was called with, the next three columns represent the genomic coordinates of the eccDNA forming region, and the last column represents the number of junction split reads used to call the eccDNA forming region.

Additional File 21: List of genes annotated in the *M. oryzae* Guy11 genome along with other information discussed in this study for each gene.

The first three columns describe the genomic coordinates of the gene, the fourth column is the gene’s ID, the fifth column describes whether the gene was predicted to be an effector, the sixth column lists its name if it is a known effector, the seventh column lists the name of the protein in the *M. oryzae* 70-15 proteome, the eighth column describes whether it is an eccDNA-associated or eccDNA-absent gene, and the last column describes whether this gene was kept in all rice-infecting *M. oryzae* genomes analyzed.

Additional File 22: Enriched GO terms in the cellular components ontology for eccDNA-associated genes.

The first column lists the GO term, the second column lists the number of genes annotated with each term, the third column lists the number of genes observed in the eccDNA-associated category, the fourth column list the number of genes expected in that category, the fifth column shows is a description of the go term, the sixth column lists the Chi-square value for that GO term, and the final column lists the ratio of the observed number of genes in the eccDNA-associated category divided by the expected number of genes in that category.

Additional File 23: Enriched GO terms in the molecular function ontology for eccDNA-associated genes.

The first column lists the GO term, the second column lists the number of genes annotated with each term, the third column lists the number of genes observed in the eccDNA-associated category, the fourth column list the number of genes expected in that category, the fifth column shows is a description of the go term, the sixth column lists the Chi-square value for that GO term, and the final column lists the ratio of the observed number of genes in the eccDNA-associated category divided by the expected number of genes in that category.

Additional File 24: Enriched GO terms in the biological pathway ontology for eccDNA-associated genes.

The first column lists the GO term, the second column lists the number of genes annotated with each term, the third column lists the number of genes observed in the eccDNA-associated category, the fourth column list the number of genes expected in that category, the fifth column shows is a description of the go term, the sixth column lists the Chi-square value for that GO term, and the final column lists the ratio of the observed number of genes in the eccDNA-associated category divided by the expected number of genes in that category.

Additional File 25: List of GenBank accessions for the genomes of rice-infecting *M. oryzae* isolates used in this study for gene annotation.

Additional File 26: List of small, genic deletions identified in the *M. oryzae* Guy11 genome.

The first three columns describe genomic coordinates of the deletion, the fourth column is the missing gene’s ID, and the last column is the name of the genome where the deletion is present.

Additional File 27: List of GenBank accessions for the genomes of *M. oryzae* used in this study to search for eccDNA-mediated translocations.

Additional File 28: Enriched GO terms in the cellular components ontology for eccDNA-absent genes.

The first column lists the GO term, the second column lists the number of genes annotated with each term, the third column lists the number of genes observed in the eccDNA-absent category, the fourth column list the number of genes expected in that category, the fifth column shows is a description of the go term, the sixth column lists the Chi-square value for that GO term, and the final column lists the ratio of the observed number of genes in the eccDNA-associated category divided by the expected number of genes in that category.

Additional File 29: Enriched GO terms in the molecular function ontology for eccDNA-absent genes.

The first column lists the GO term, the second column lists the number of genes annotated with each term, the third column lists the number of genes observed in the eccDNA-absent category, the fourth column list the number of genes expected in that category, the fifth column shows is a description of the go term, the sixth column lists the Chi-square value for that GO term, and the final column lists the ratio of the observed number of genes in the eccDNA-associated category divided by the expected number of genes in that category.

Additional File 30: Enriched GO terms in the biological pathway ontology for eccDNA-absent genes.

The first column lists the GO term, the second column lists the number of genes annotated with each term, the third column lists the number of genes observed in the eccDNA-absent category, the fourth column list the number of genes expected in that category, the fifth column shows is a description of the go term, the sixth column lists the Chi-square value for that GO term, and the final column lists the ratio of the observed number of genes in the eccDNA-associated category divided by the expected number of genes in that category.

Additional File 31: List showing names of barcodes used for each PacBio sequencing sample.

Additional File 32: Sequences of barcodes used for library preparation of PacBio sequencing samples in FASTA format.

Additional File 33: Consensus sequences of LTR retrotransposons in the *M. oryzae* Guy11 genome in FASTA format.

Additional File 34: Genome, gene annotation, and transposable element annotation files used for comparative circularome analysis.

Additional File 35: List of SRA accessions for RNAseq data used in this study.

