## Additional File 1 for "The extrachromosomal circular DNAs of the rice blast pathogen *Magnaporthe oryzae* contain a wide variety of LTR retrotransposons, genes, and effectors"

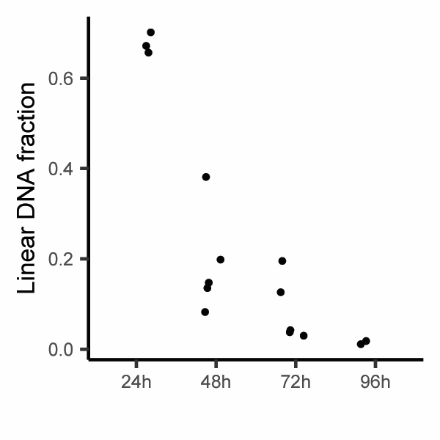

**Fig. S1.** Degradation of linear DNA using exonuclease treatment. Scatter plot showing the effect of exonuclease treatment on linear DNA fraction of total extracted DNA from *M. oryzae* tissue samples. Each dot represents one biological replicate averaged across four qPCR replicates.

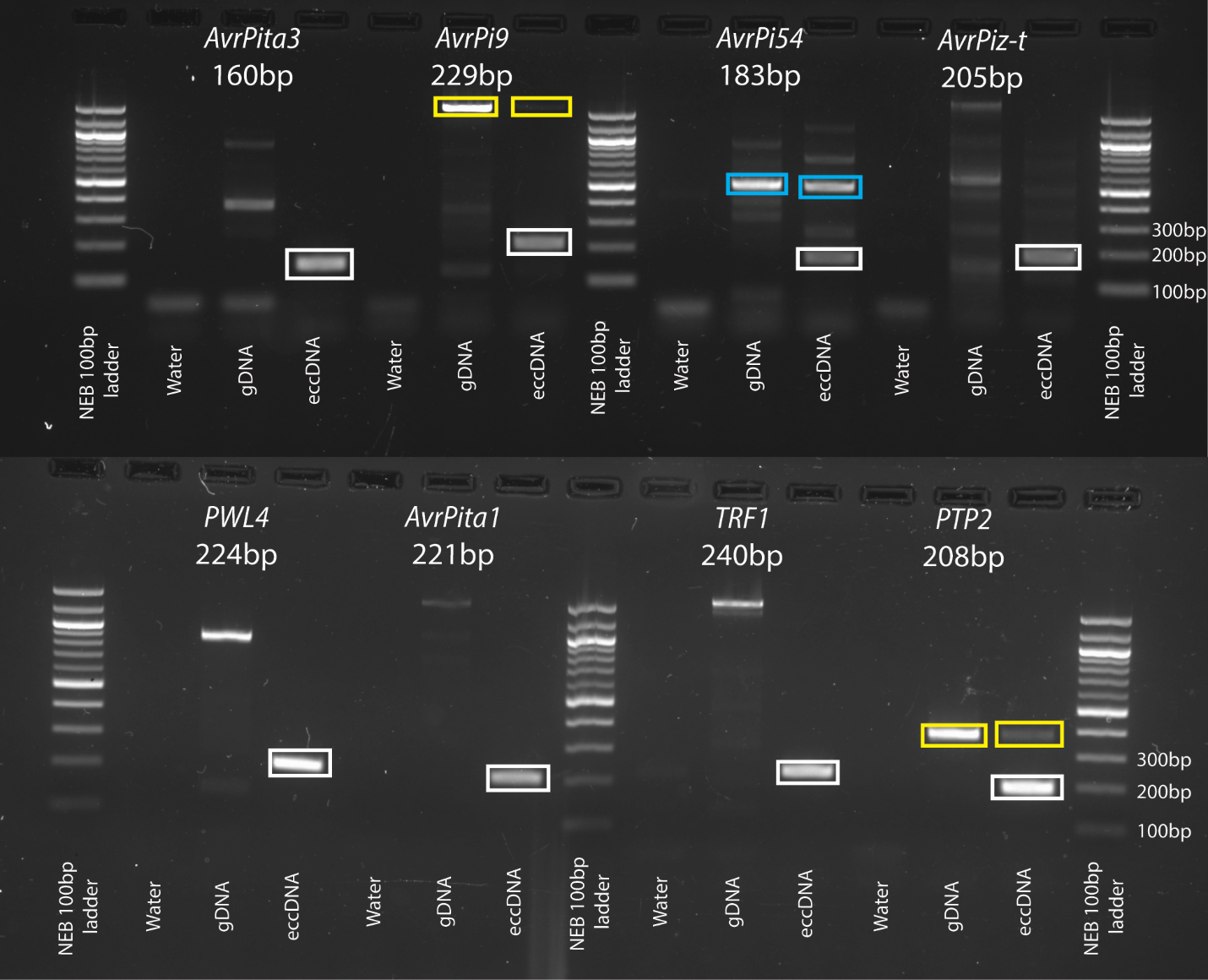

**Fig. S2.** Outward PCR validation of eccDNA forming regions. Genes of interest found in eccDNA forming regions are listed for each group of three samples. One primer set was used per group and the expected product size is written below the gene name. All samples for each product were from the same PCR reaction. All boxes indicate PCR products that were Sanger sequenced. White boxes indicate PCR products that matched the expected eccDNA junctions. Yellow boxes indicate PCR products that originated from continuous sequences of DNA present in both the genomic DNA and on a high confidence eccDNA forming region found in the eccDNA sample. Blue boxes indicate PCR products with different sequences. PCR for *AvrPita3*, *AvrPi9*, *AvrPi54*, *AvrPiz-t*, and *TRF1* junctions were all performed using biological replicate 1, technical replicate A. PCR for *AvrPita1*, and *PTP2* junctions were performed using biological replicate 1, technical replicate C. PCR for the *Pwl4* junction was performed using biological replicate 2, technical replicate A.

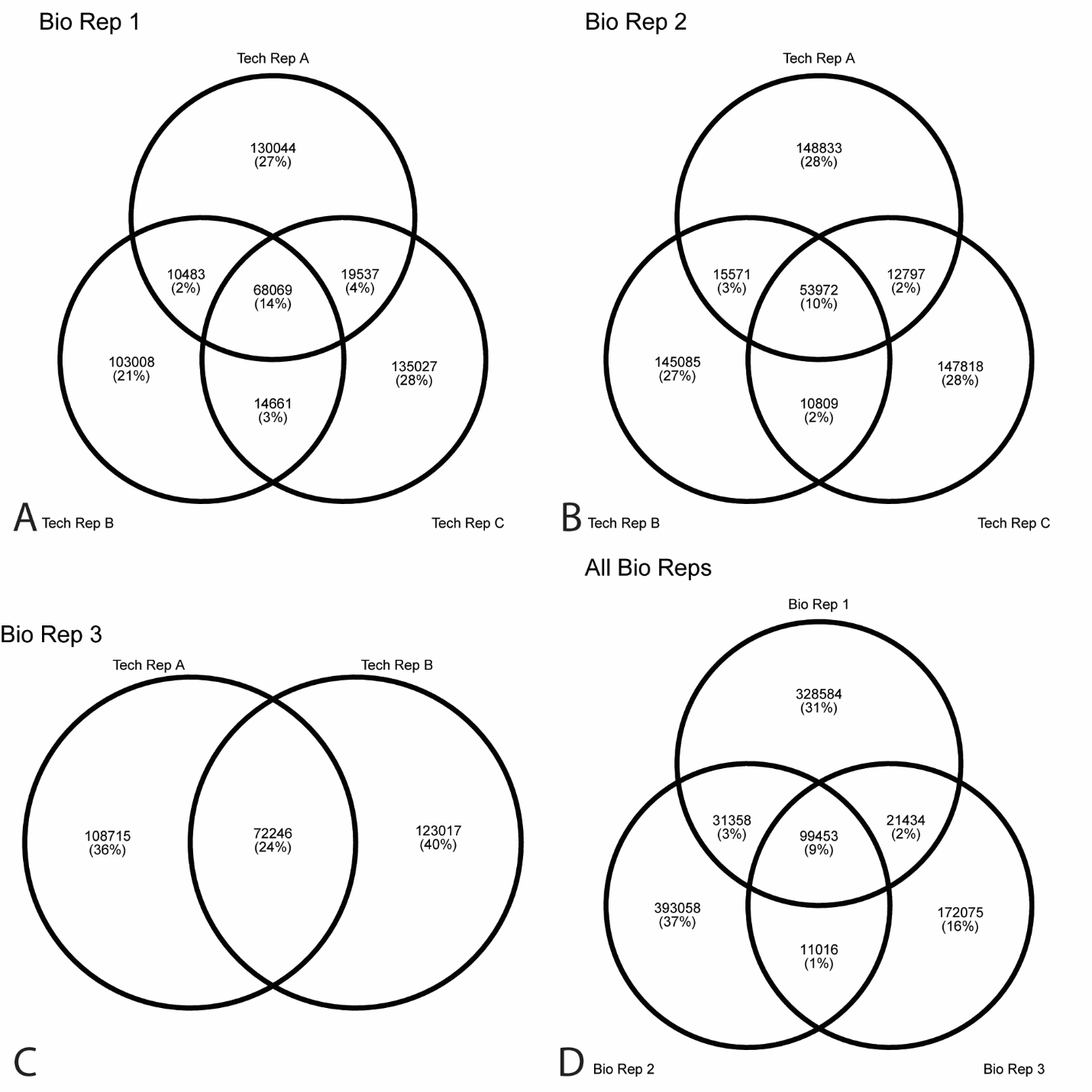

**Fig. S3.** Overlap in exact break points of eccDNA forming regions across samples. Venn diagrams showing the number of eccDNA forming regions sharing exact coordinates across technical replicates (**A-C**) and all biological replicates (**D**). EccDNA forming regions from all technical replicates for each biological replicate were merged before they were compared between biological replicates.

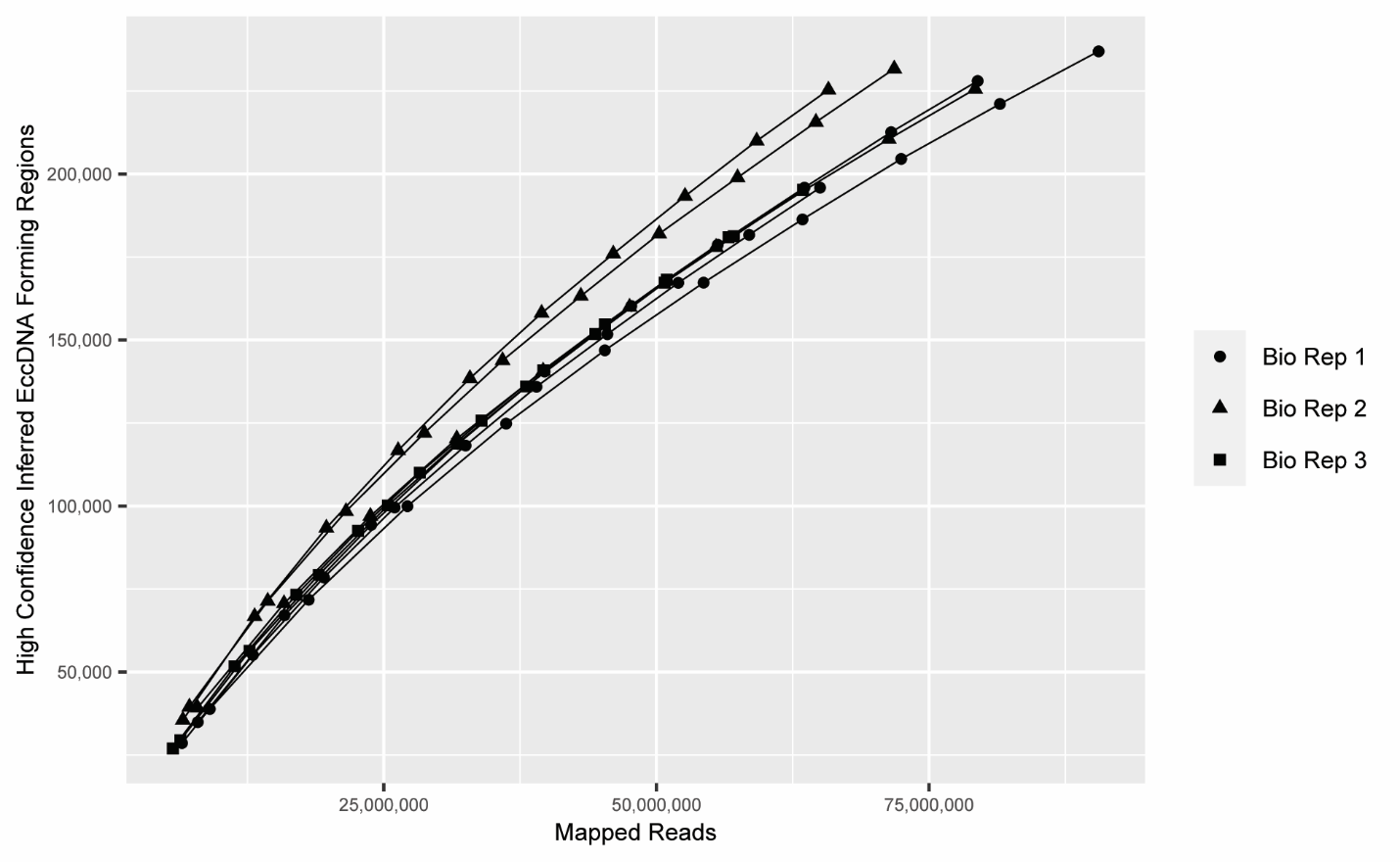

**Fig. S4.** Rarefaction analysis of sequencing coverage and eccDNA forming regions across all samples. Rarefaction curves showing the number of eccDNA forming regions called at each subset of total mapped reads for each sample. Each dot represents one subsample of mapped sequencing reads for one sequenced sample.

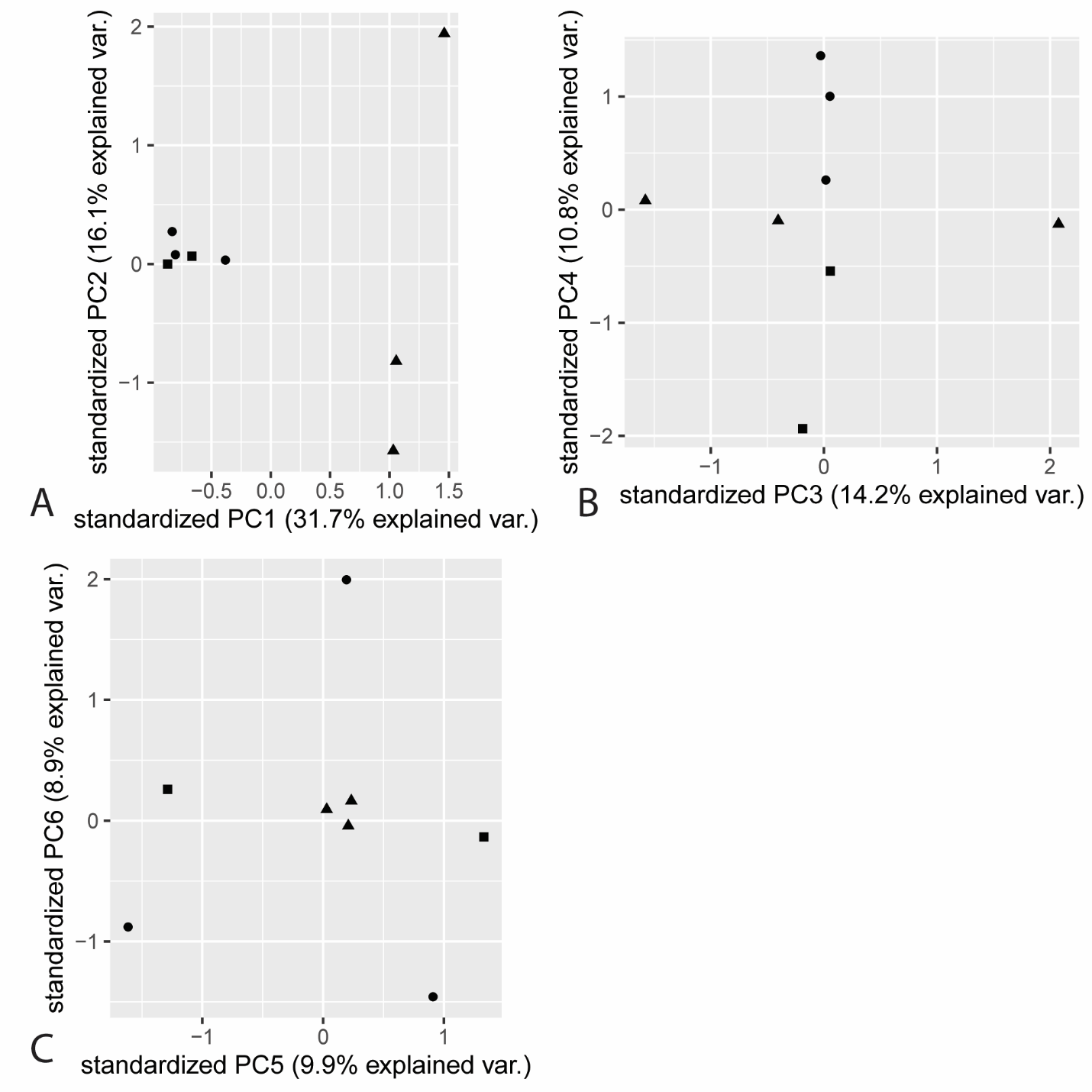

**Fig. S5.** Principal components analysis of sequencing coverage between samples. Biplots showing values of each sample for the first six principal components (PCs) generated from a principal components analysis performed using sequencing read coverage of all 10kbp bins across the *M. oryzae* Guy11 genome. Each dot represents one sample, and the shape of the dots represent the biological replicate each sample was taken from.

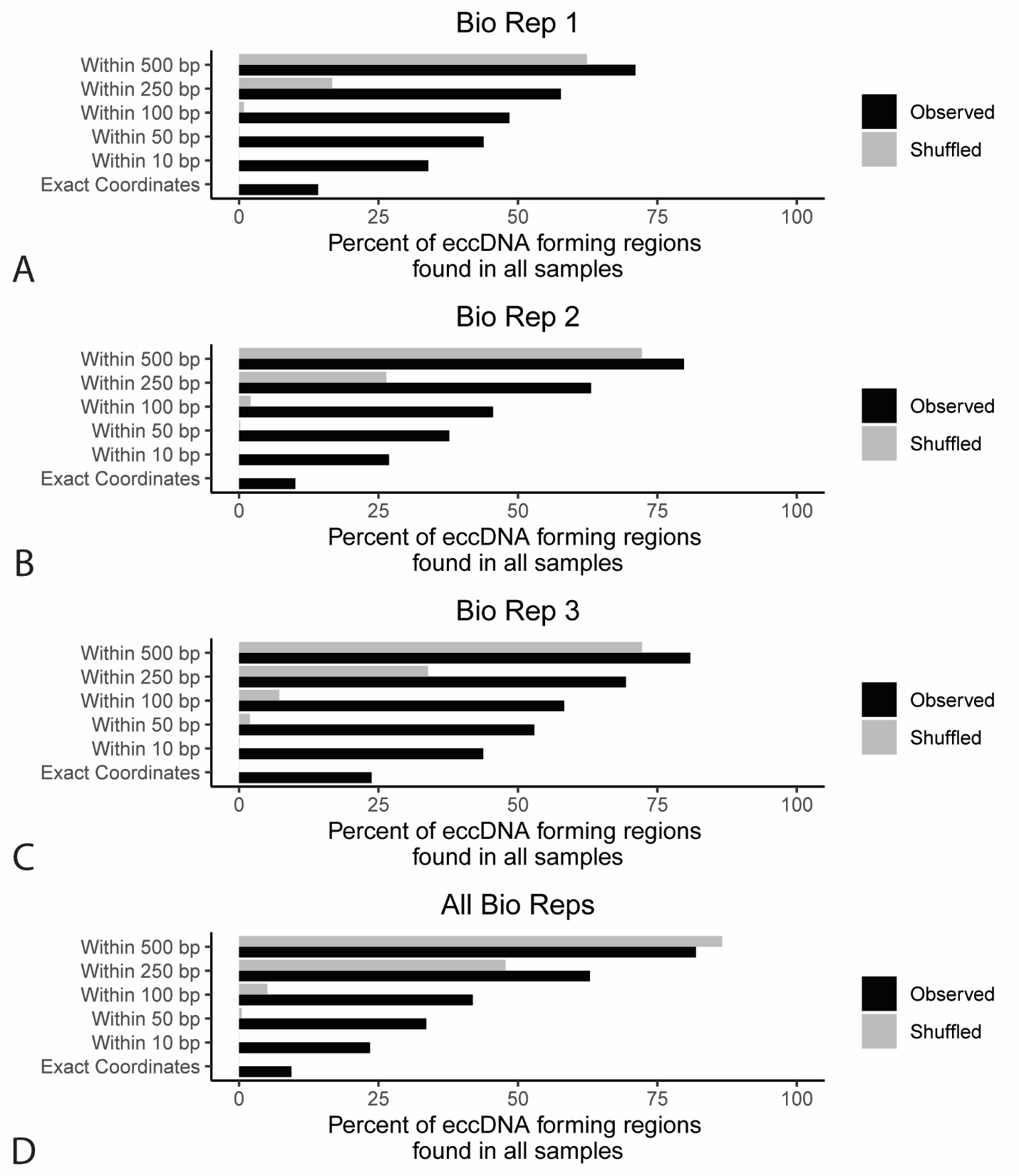

**Fig. S6.** Overlap in eccDNA forming regions across samples, with increasing tolerance for start and end coordinates. Histogram showing percentage of eccDNA forming regions found in all technical replicates for each biological replicate (**A-C**) as well as percentage of eccDNA forming regions found in all biological replicates (**D**). Percentages are shown for comparison of eccDNA forming regions based off exact coordinates as well as increasing levels of tolerance when comparing the start and end coordinates of the eccDNA forming regions. EccDNA forming regions from all technical replicates for each biological replicate were merged before they were compared between biological replicates. This observed data was compared to data obtained by randomly placing eccDNA forming regions throughout the genome for each sample. Percentages shown for these shuffled data points are the mean of 100 randomized trials. Standard deviations were too small to visualize meaningfully in the figure.

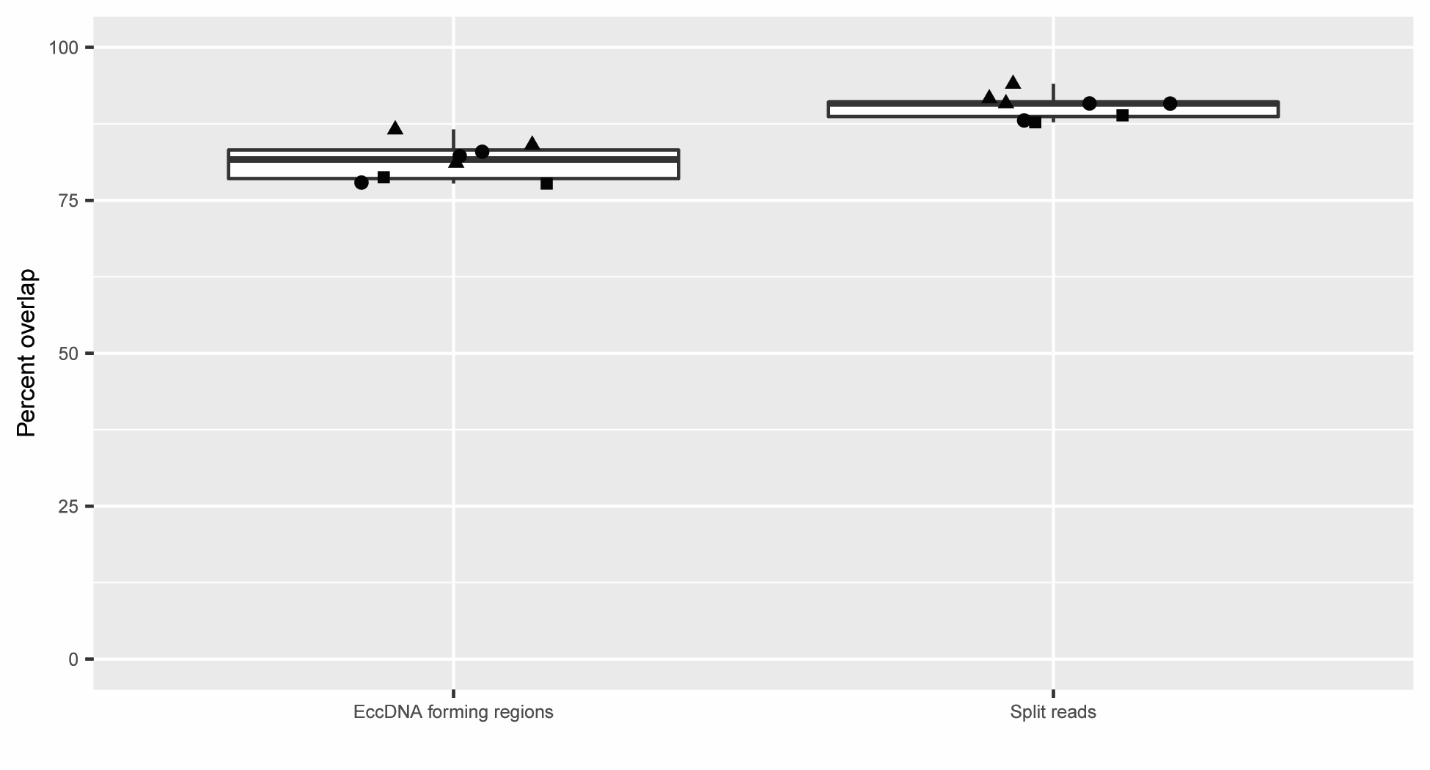

**Fig. S7**. Overlap between eccDNA forming regions called using PacBio sequencing data and Illumina sequencing data. Boxplot showing the percentage of eccDNA forming regions that were found in each sample using our PacBio sequencing data that were also represented in either eccDNA forming regions called using our Illumina sequencing data or split reads found in this data. Each point represents one sample, and the shape of the points represent the biological replicate that sample was taken from.

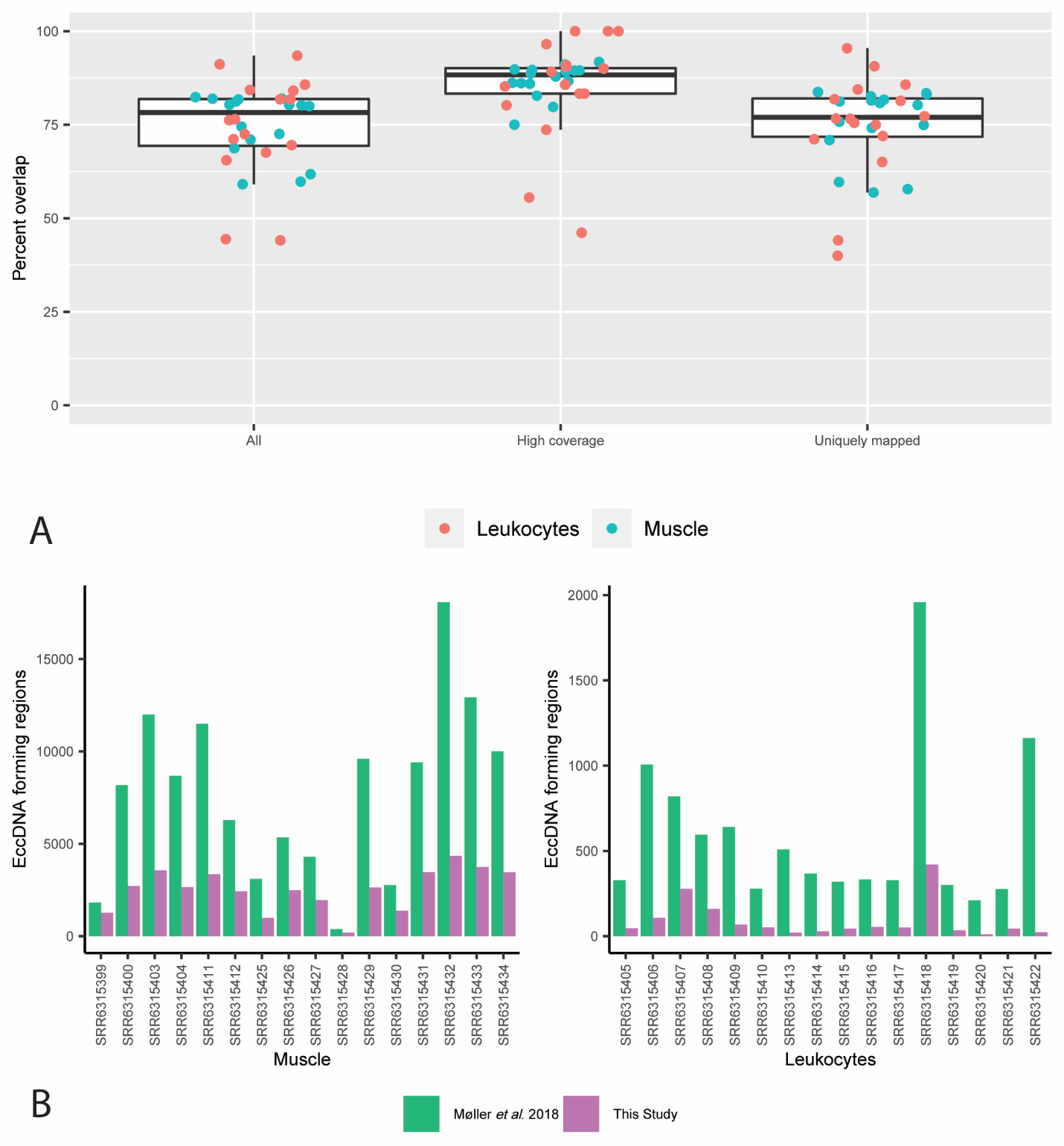

**Fig. S8.** Comparison between eccDNA forming regions in human samples called in this manuscript and in the original publication. **A.** Boxplot showing the percentage of eccDNA forming regions that were found using our pipeline that were also found in the published eccDNA forming regions for human samples. Each dot represents one sample. **B.** Bar plot showing counts of eccDNA forming regions generated from our pipeline compared to counts in published data. Sample IDs we taken from the Sequence Read Archive (SRA).

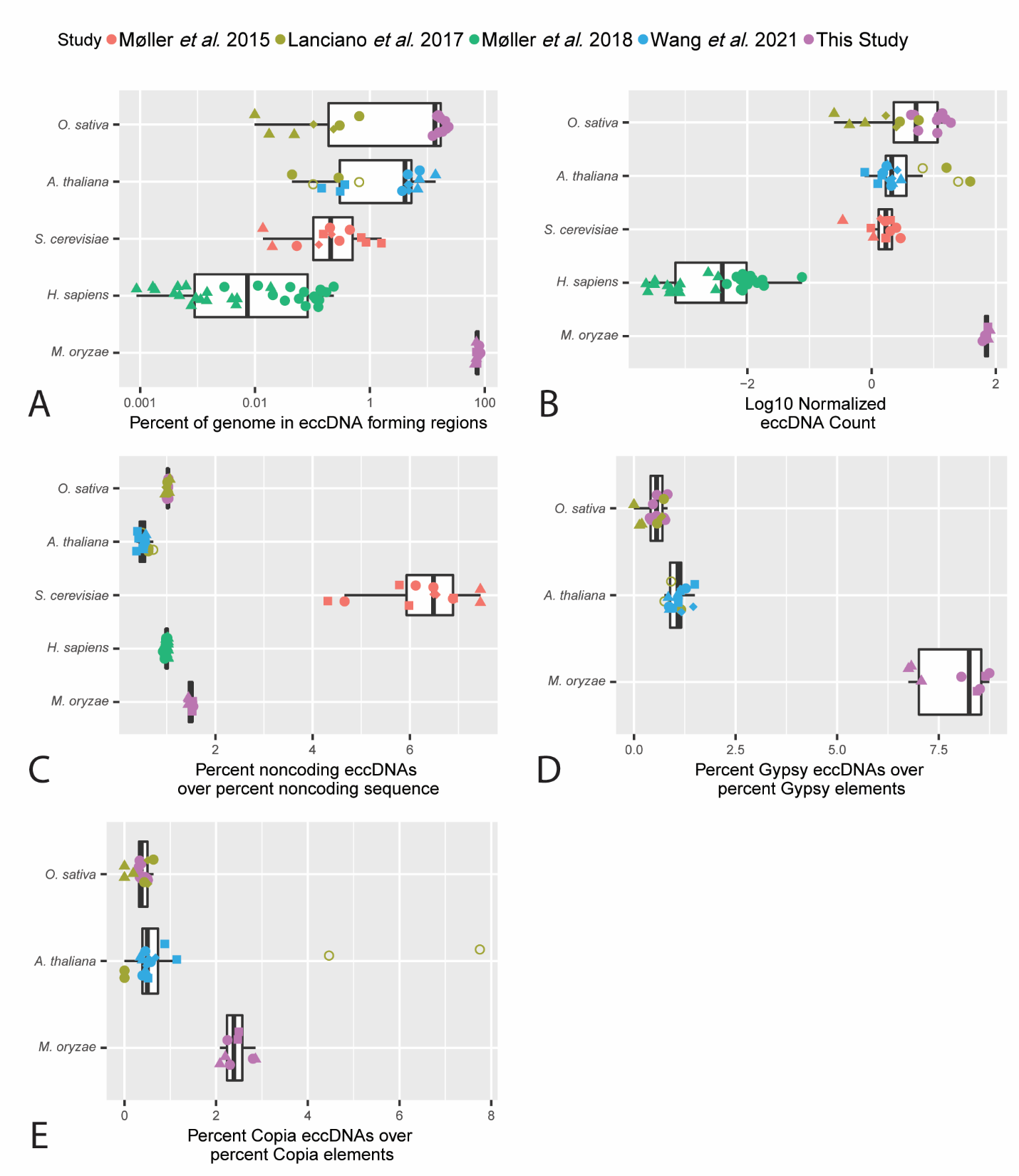

**Fig. S9.** Comparison of eccDNA forming regions between *M. oryzae* and other previously studied organisms. Box plots comparing **A.** the percentage of the genome found in eccDNA forming regions, **B.** log 10 count of eccDNA forming regions normalized to genome size and sequencing library size, **C.** percent of eccDNA forming regions that contain more than 50% noncoding sequences divided by percent of the genome made up of noncoding seqeuence, **D.** percent of eccDNA forming regions that contain more than 90% LTR/Gypsy retrotransposon sequence divide by percent of the genome made up of LTR/Gypsy retrotransposon sequence, **E.** percent of eccDNA forming regions that contain more than 90% LTR/Copia retrotransposon sequence divided by percent of the genome made up of LTR/Gypsy retrotransposon sequence across multiple organisms and studies. Each dot represents one sequenced sample. Shapes represent variations in sample type within the same organism. For *M. oryzae*, shapes correspond to which biological replicate each sample was taken from. For *Oryza sativa*, circles represent leaf samples, triangles represent callus samples and diamonds represent seed samples. For *Homo sapiens*, circles represent muscle samples and triangles represent leukocyte samples. For *Arabidopsis thaliana*, circles represent wild type flower samples, empty circles represent *epi12* mutant flower samples, squares represent root samples, diamonds represent leaf samples and triangles represent stem samples. For *Saccharomyces cerevisiae*, circles represent samples from the yeast deletion collection, squares represent samples from the yeast deletion collection treated with zeocin, triangles represent samples from *GAP1* circle carrying yeast, diamonds represent samples from clonal isogenic haploid S228C yeast. For the retrotransposon boxplots, *H. sapiens* samples were excluded due to a lack of active LTR/Gypsy and LTR/Copia retrotransposons in their genome [89] and *S. cerevisiae* samples were excluded due to a small number of eccDNA forming regions containing retrotransposon sequences.

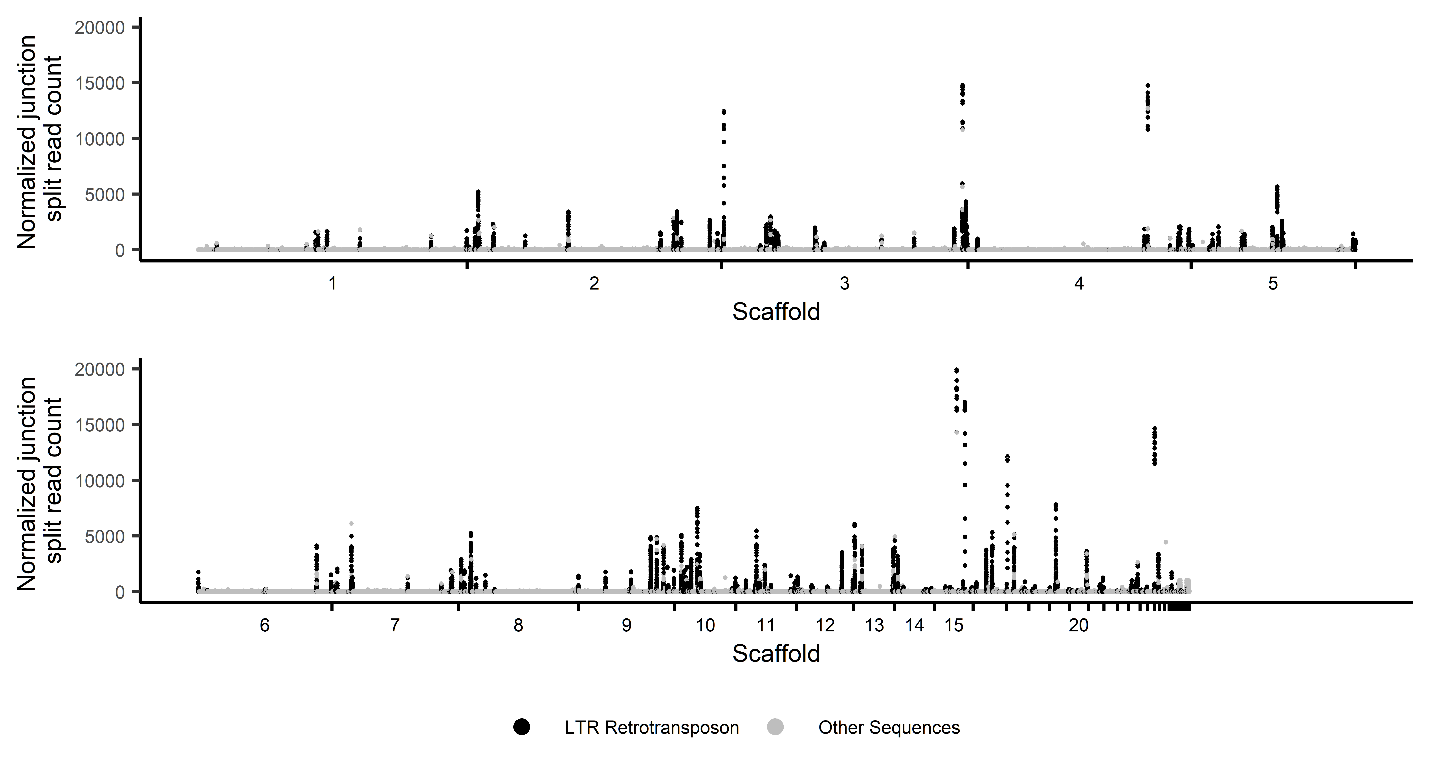

**Fig. S10.** EccDNA forming regions composed of more than 90% LTR retrotransposon sequence in *M. oryzae*. Manhattan plot showing the number of junction split reads per million averaged across biological replicates for all 100 bp bins with junction split read coverage greater than zero in the *M. oryzae* Guy11 genome. Each dot represents one of these bins. Bins made up of more than 90% LTR retrotransposon sequence are colored in black.

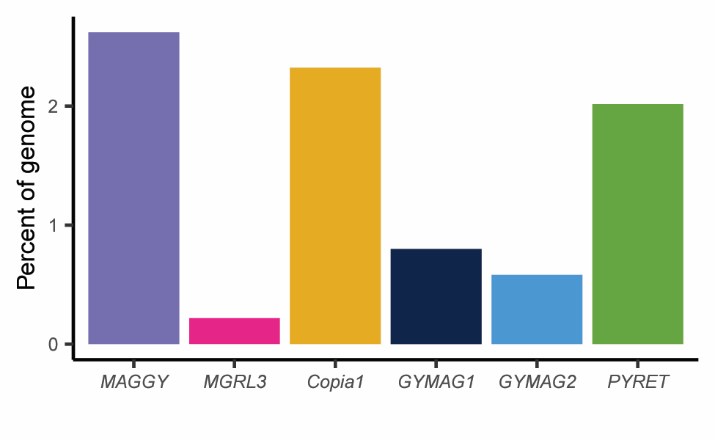

**Fig. S11.** Percentage of the *M. oryzae* Guy11 genome made up of each LTR retrotransposon.

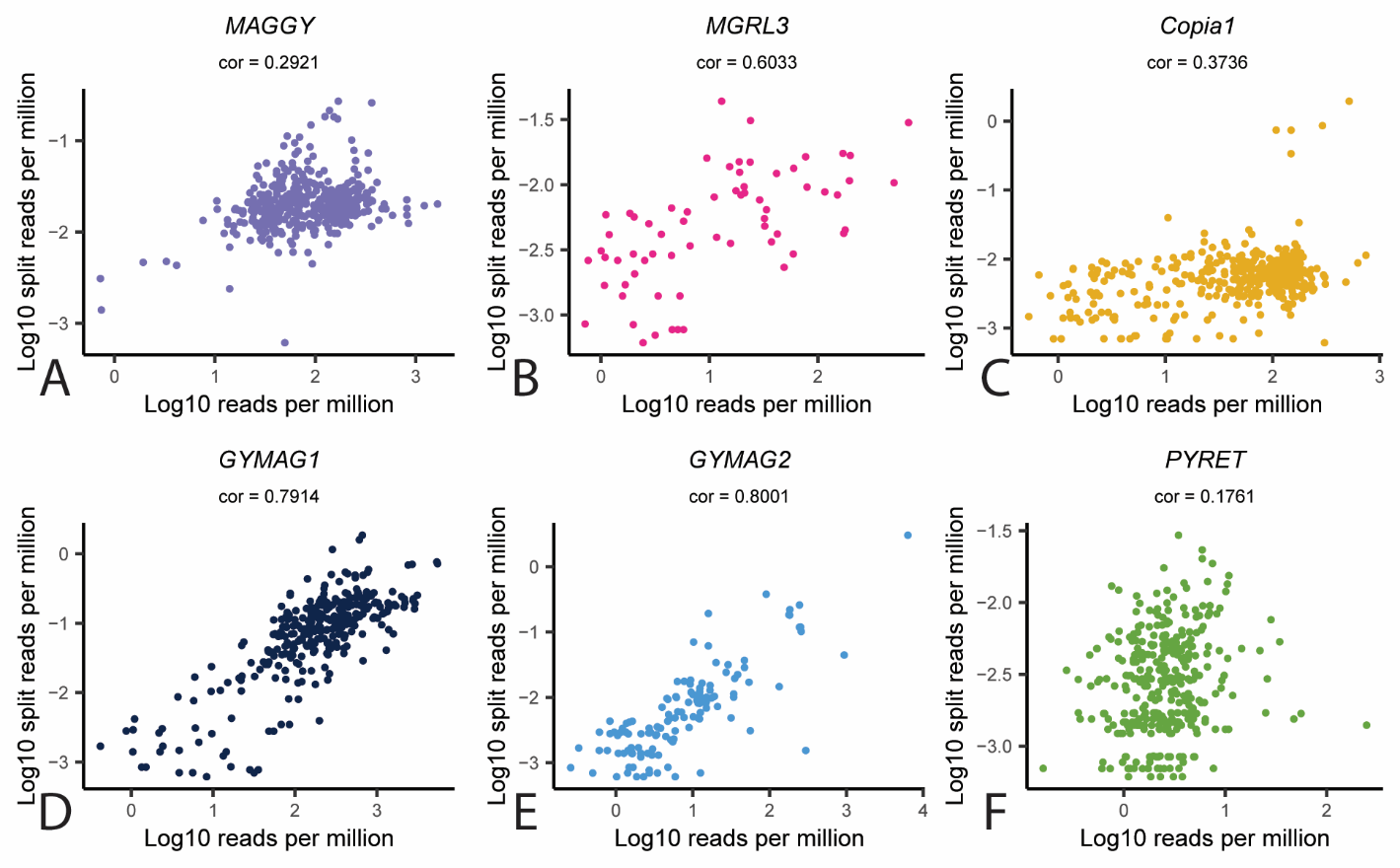

**Fig. S12.** Correlation between number of LTR-LTR split reads and sequencing reads in eccDNA sequencing samples for each LTR retrotransposon in *M. oryzae*. **A-F**. Scatter plots showing Pearson’s correlation coefficient between log 10 sequencing reads per million reads and log 10 LTR-LTR split reads per million sequencing reads, averaged across biological replicates. Each dot represents one annotated portion of the LTR region of an LTR retrotransposon.

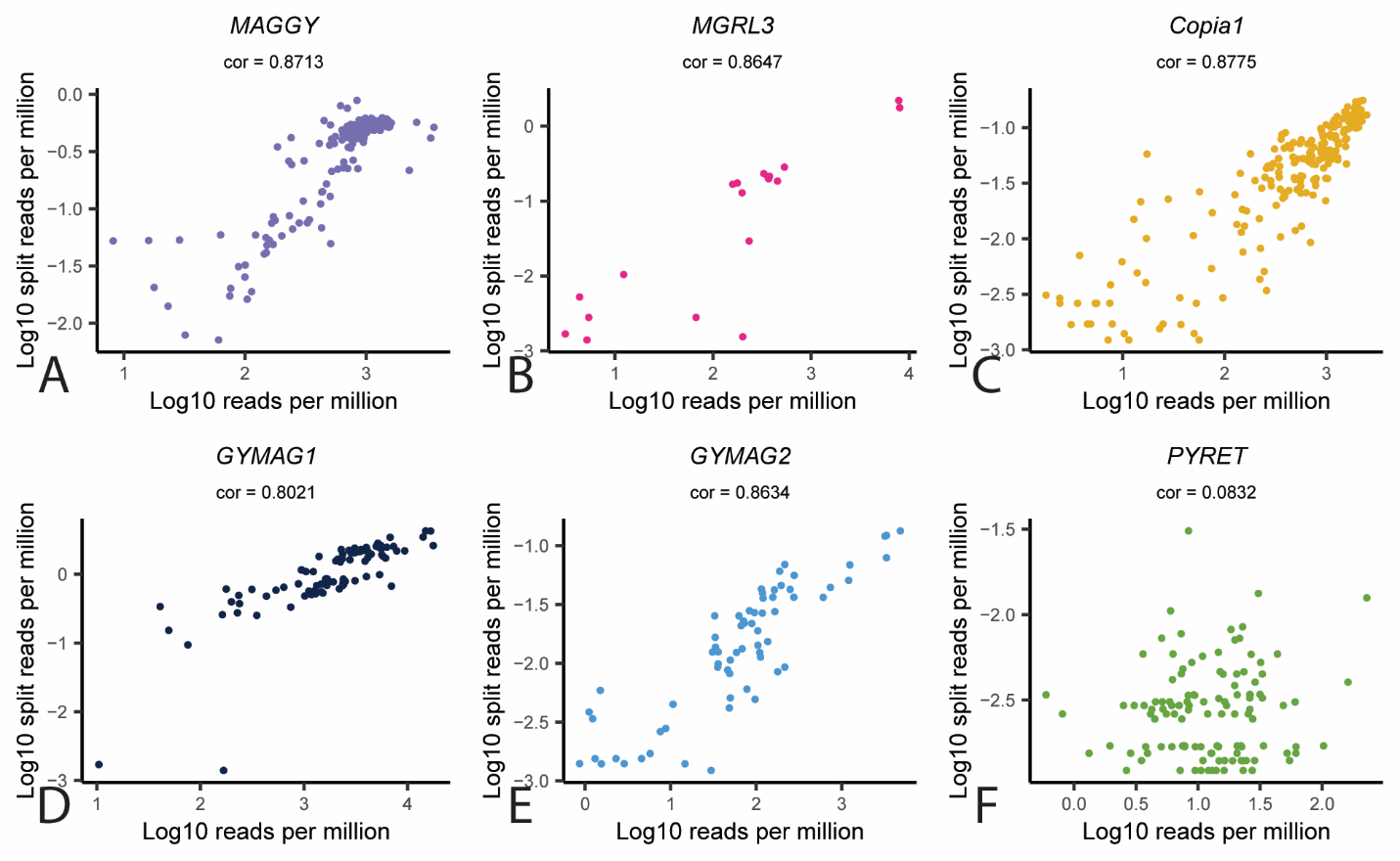

**Fig. S13.** Correlation between number of LTR-internal split reads and sequencing reads in eccDNA sequencing samples for each LTR-retrotransposon in *M. oryzae*. **A-F**. Scatter plots showing Pearson’s correlation between log 10 sequencing reads per million reads and log 10 LTR-internal split reads per million sequencing reads, averaged across biological replicates. Each dot represents one annotated portion of the internal region of an LTR retrotransposon.

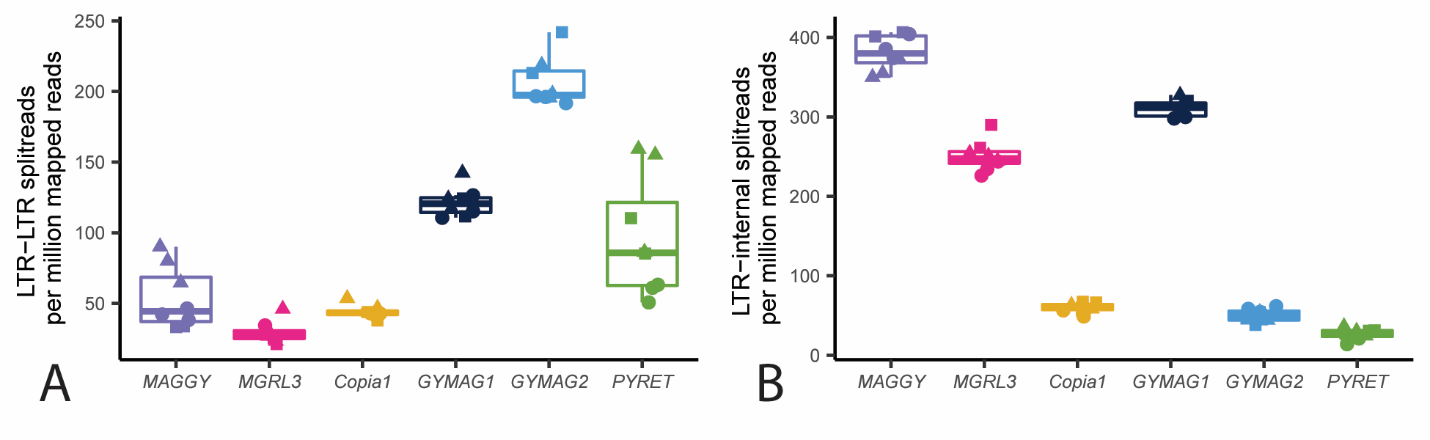

**Fig. S14.** Number of LTR-LTR split reads and LTR-internal split reads in eccDNA sequencing samples for each LTR retrotransposon in *M. oryzae.* **A.** Box plot showing identified LTR-LTR split reads per million reads mapped to each element for each LTR retrotransposon in the *M. oryzae* Guy11 genome. Each point represents one sample. **B.** Box plot showing identified LTR-internal split reads per million reads mapped to each element, for each LTR retrotransposon in the *M. oryzae* Guy11 genome. Each point represents one sample, and the shape of the points represent the biological replicate that sample was taken from.

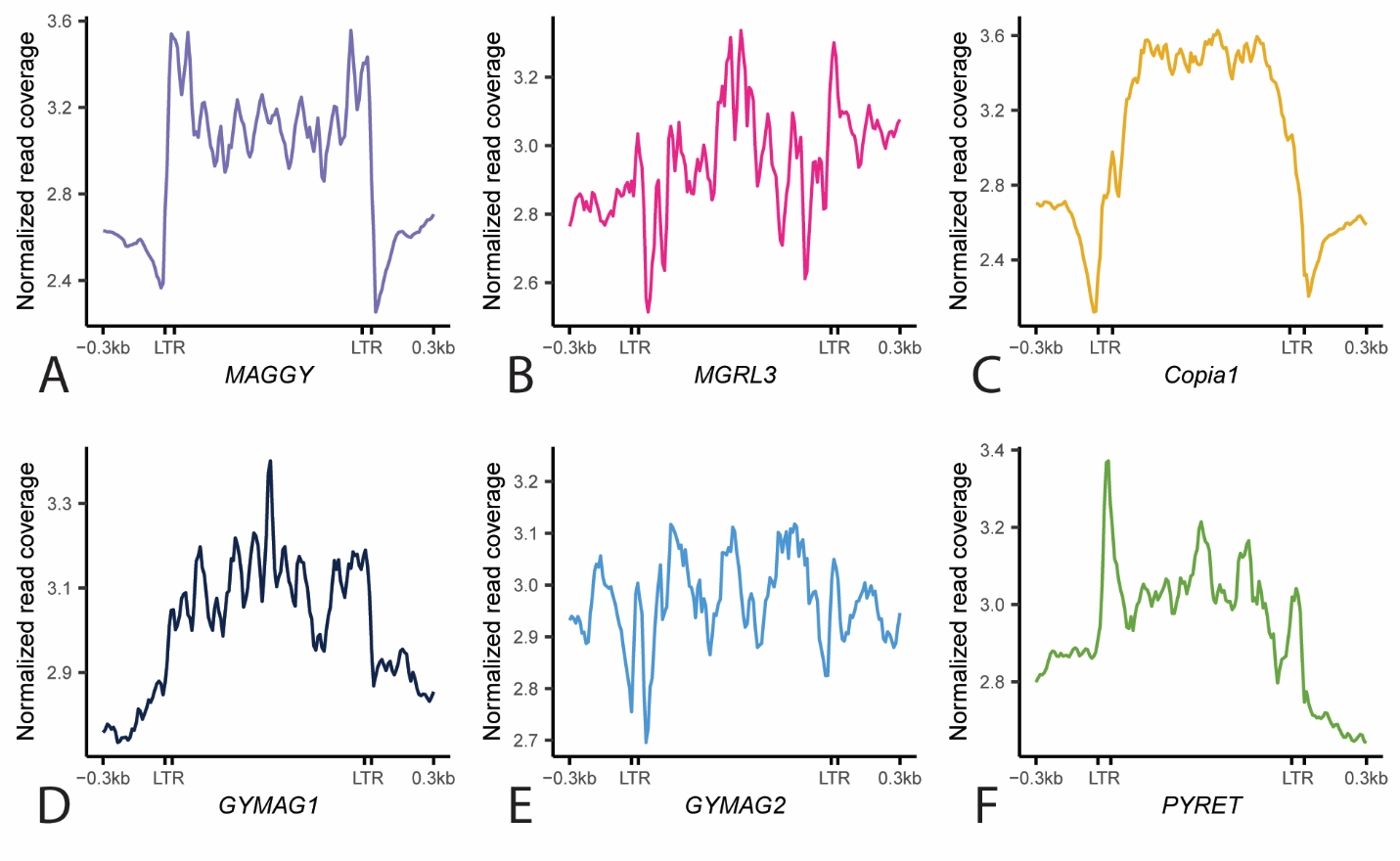

**Fig. S15**. Expected read coverage for LTR retrotransposons in *M. oryzae*. **A-F**. Profile plots showing observed whole genome sequencing read coverage for each LTR retrotransposon found in the *M. oryzae* Guy11 genome.

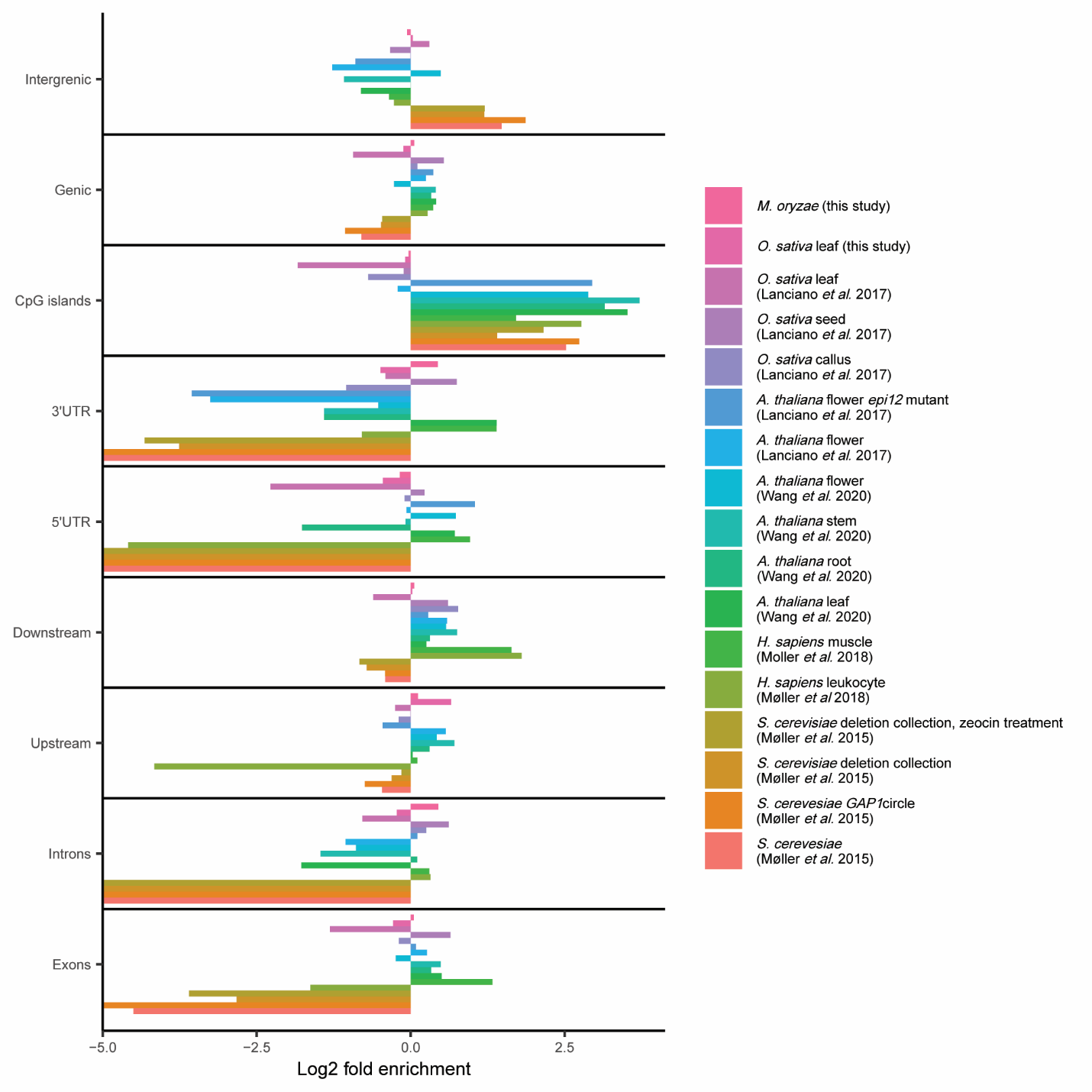

**Fig. S16.** MicroDNA enrichment and depletion in the genomes of various organisms. Bar plot showing observed enrichment of microDNAs across various regions of the genome across different previously sequenced organisms and sample types. Log2 fold enrichment of -5 represents samples where no microDNAs were found in that region. The presented data is an average of all sequenced samples of each type.

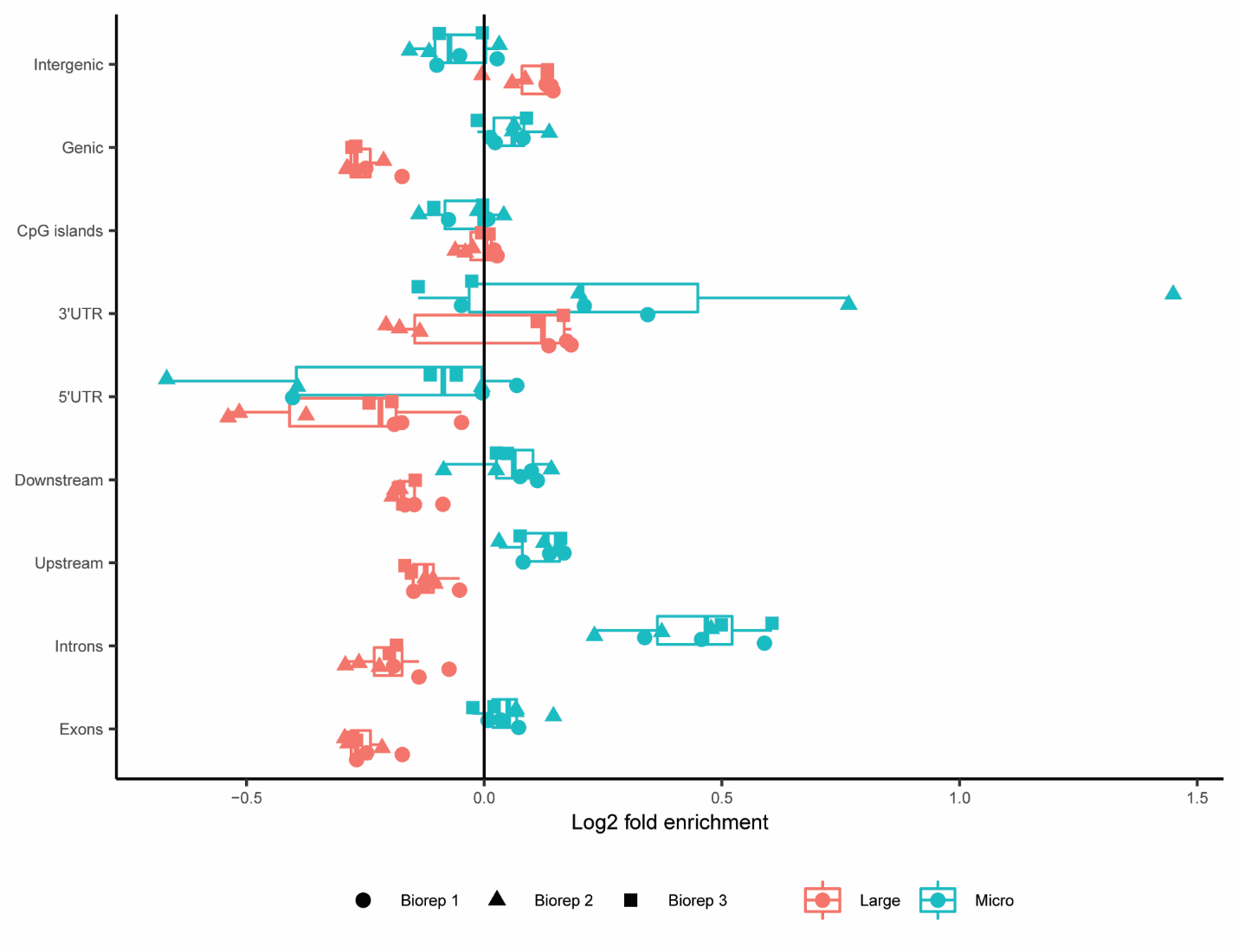

**Fig. S17.** Enrichment and depletion of microDNAs and large eccDNAs across various genomic regions in *M. oryzae*. Box plot showing observed enrichment of microDNAs and large eccDNAs across various regions of the genome. Each point represents one sample, and the shape of the points represent the biological replicate that sample was taken from.

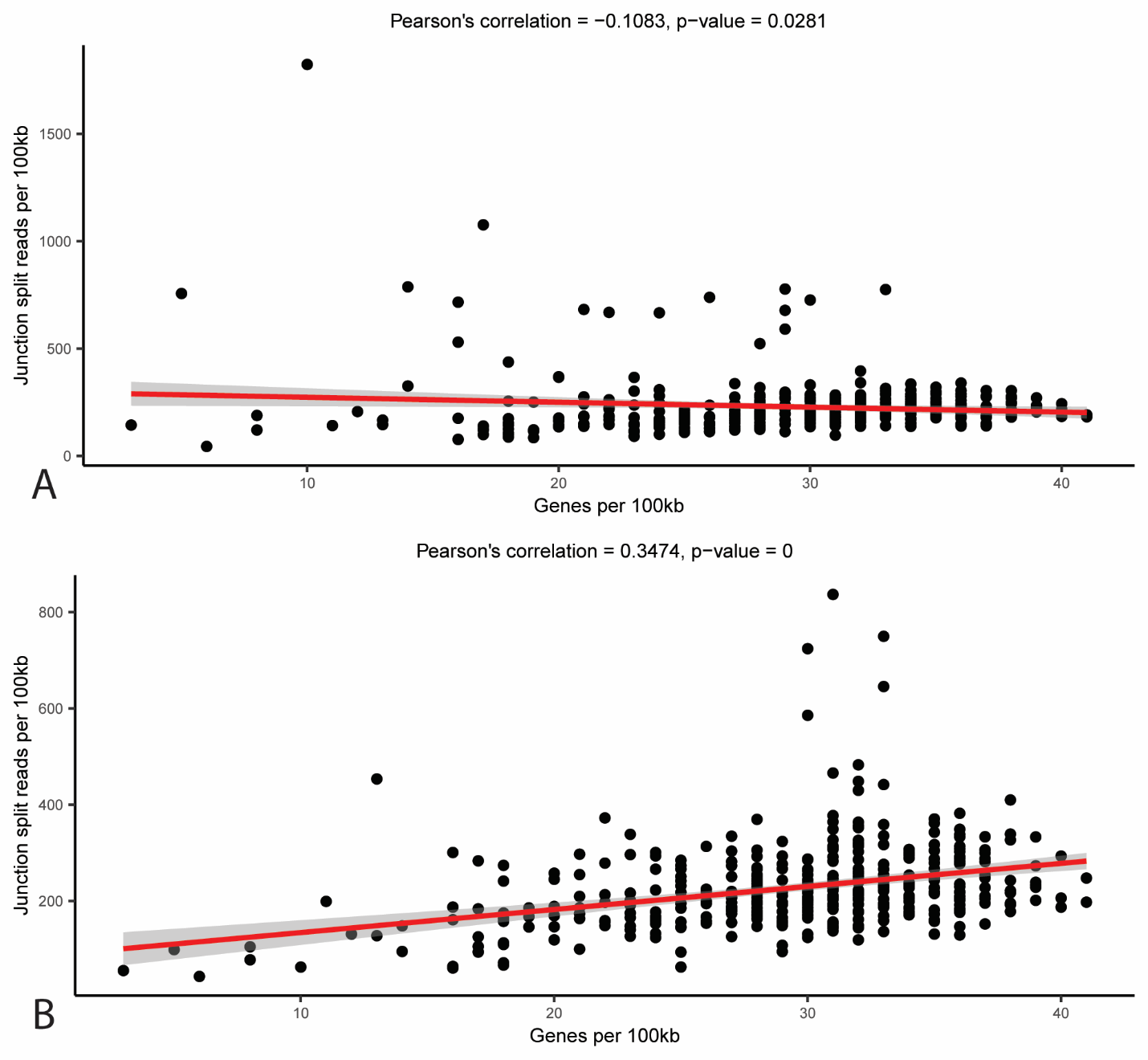

**Fig. S18.** Correlation between gene count and junction split read count across the *M. oryzae* genome. Scatter plot showing the number of genes and log 10 of the number of junction split reads per million per 100 kilobase pair bin in the *M. oryzae* Guy11 genome for **A.** large eccDNAs or **B.** microDNAs, averaged across biological replicates. The red line represents a linear regression line and the grey shadow represents 95% confidence intervals.

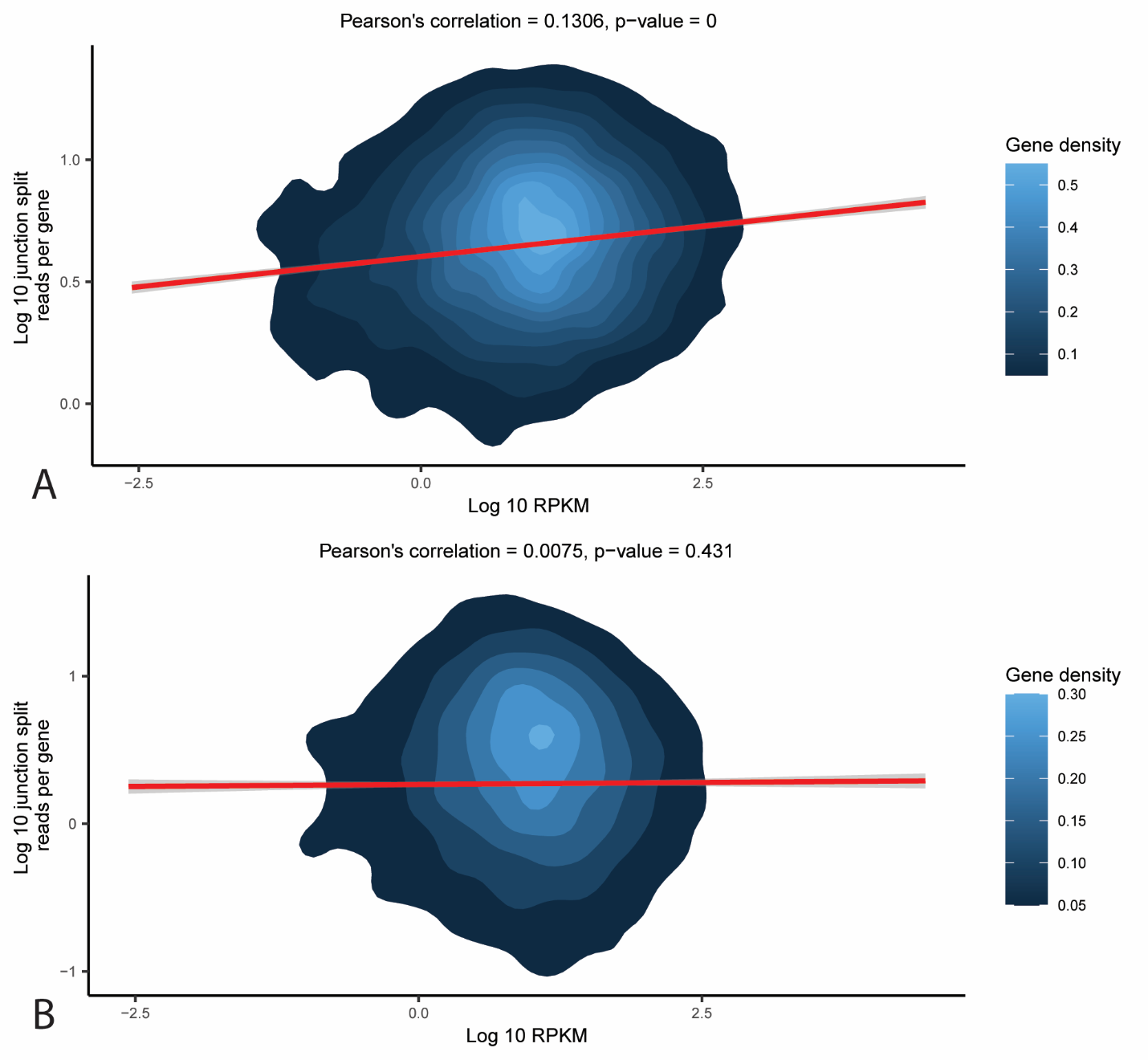

**Fig. S19.** Correlation between junction split read count and expression for *M. oryzae* genes. Two-dimensional density plot showing the log 10 of the reads per kilobase million averaged across multiple RNAseq samples and log 10 of the number of overlapping junction split reads per million for each gene for **A.** large eccDNAs and **B.** microDNAs, averaged across biological replicates. The red line represents a linear regression line and the grey shadow represents 95% confidence intervals.

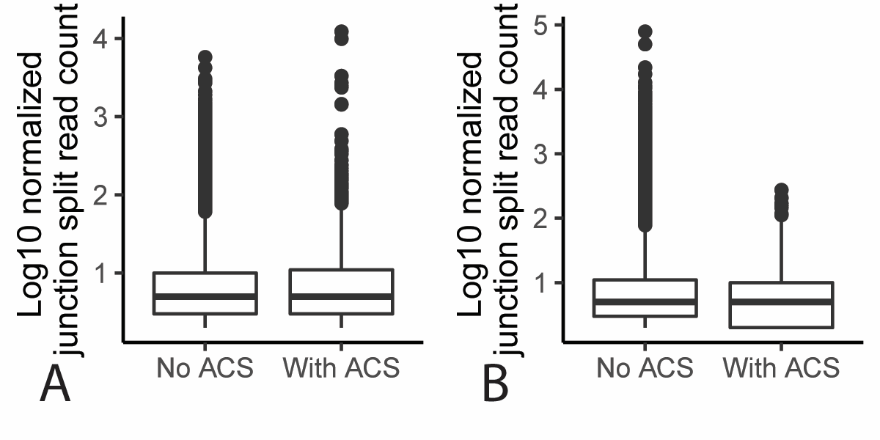

**Fig. S20.** Comparison of junction split read counts between eccDNA forming regions with and without an ACS. Box plot showing the log 10 of the number of junction split reads per million reads averaged across biological replicates for eccDNA forming regions that do and do not contain ACSs for **A.** large eccDNAs and **B.** microDNAs.

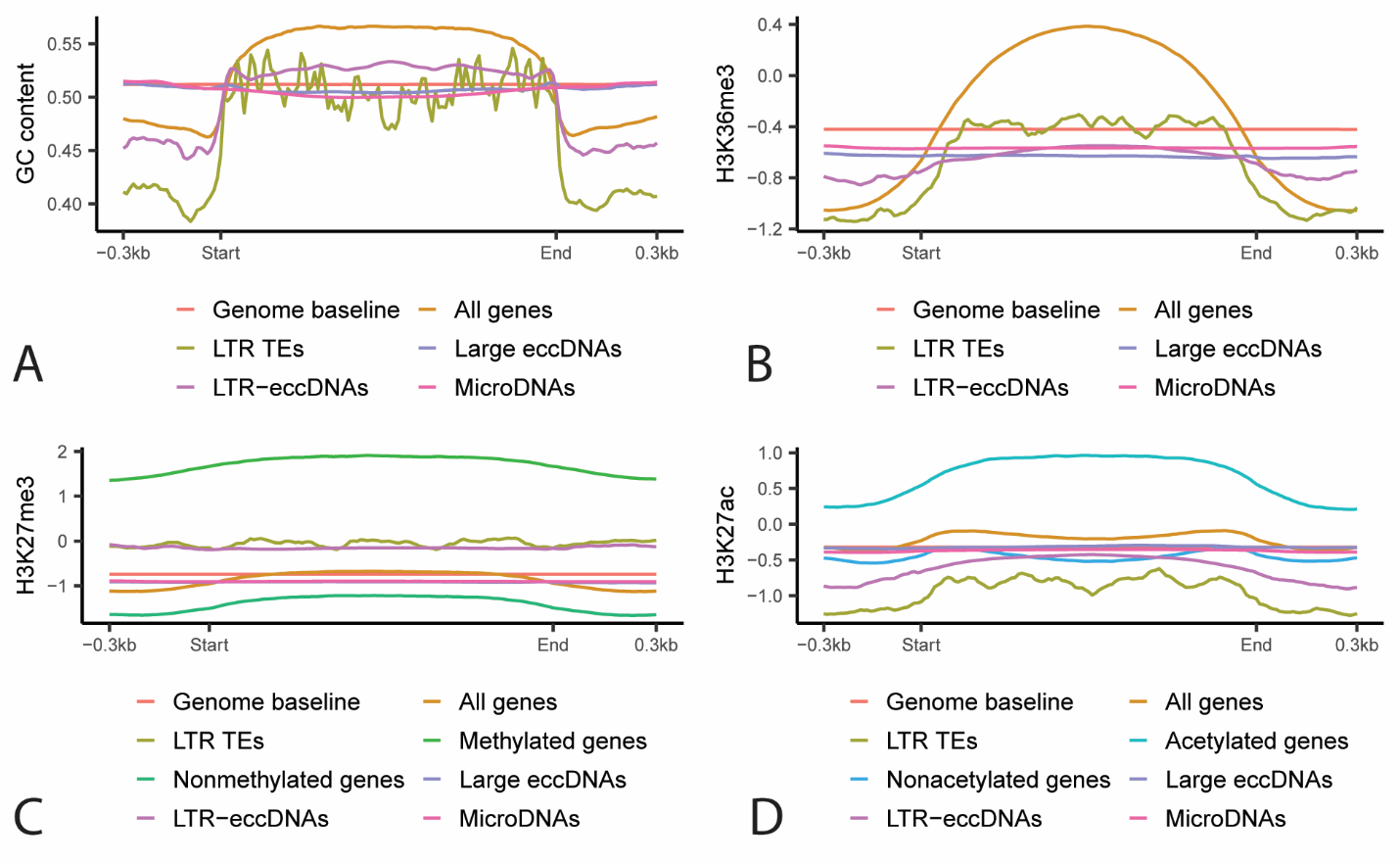

**Fig. S21.** GC content and chromatin marks of eccDNA forming regions in *M. oryzae*. Profile plots showing the average **A.** percent GC content, **B.** log2 ratio of read coverage for H3K36me3 chromatin immunoprecipitation and input control, **C.** log2 ratio of read coverage for H3K27me3 chromatin immunoprecipitation and input control and **D.** log2 ratio between read coverage for H3K27ac chromatin immunoprecipitation and input control for all *M. oryzae* genes, randomly selected regions of the genome, LTR retrotransposons, large eccDNAs, LTR-eccDNAs and microDNAs. Methylated and nonmethylated genes and acetylated and nonacetylated genes, as defined by Zhang *et al.*, are also represented in **C.** and **D.**, respectively.

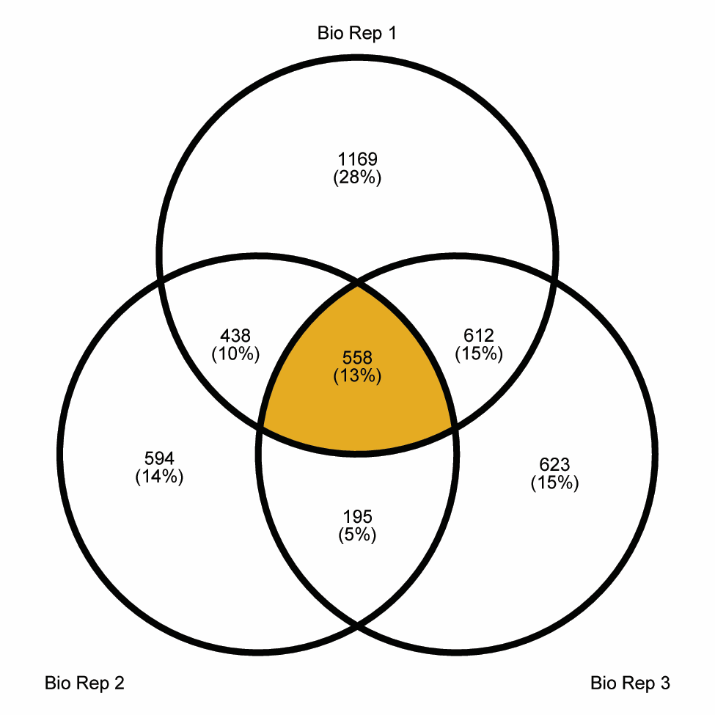

**Fig. S22.** Overlap between genes enriched on eccDNAs in biological replicates. Venn diagram showing overlap between genes in the top 33% for how often they were found fully encompassed by eccDNA forming regions in each biological replicate. Technical replicates for each biological replicates were normalized to the number of junction split reads in each sample then averaged. 558 genes found in the top 33% for all biological replicates were designated eccDNA-associated (colored in orange).

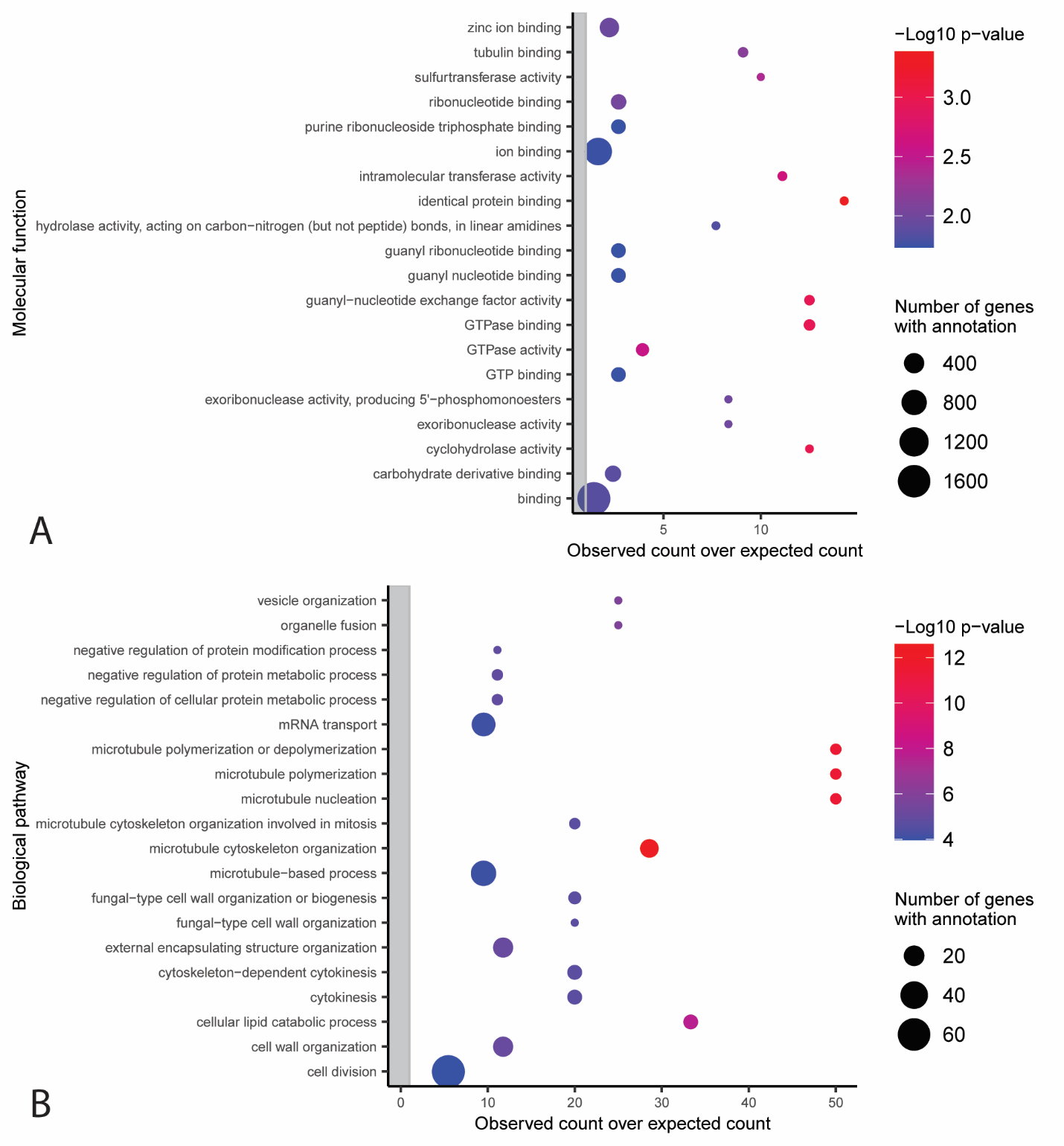
 **Fig. S23.** GO terms associated with eccDNA-associated genes. Functional categories in the **A.** molecular function and **B.** biological pathway Gene Ontology with an observed number of eccDNA-associated genes that is significantly different from the expected number with correction for gene length bias (Chi-square test, p < 0.05). The y-axis shows the different functional categories, and the x-axis represents the observed number of genes divided by the expected number of genes in this group. Dots outside of the grey rectangle represent functional categories that are observed more often than expected. The size of dots indicates the total number of genes in the *M. oryzae* genome that belong to each functional category. Only the 20 categories with the largest -log10 p-values are shown.

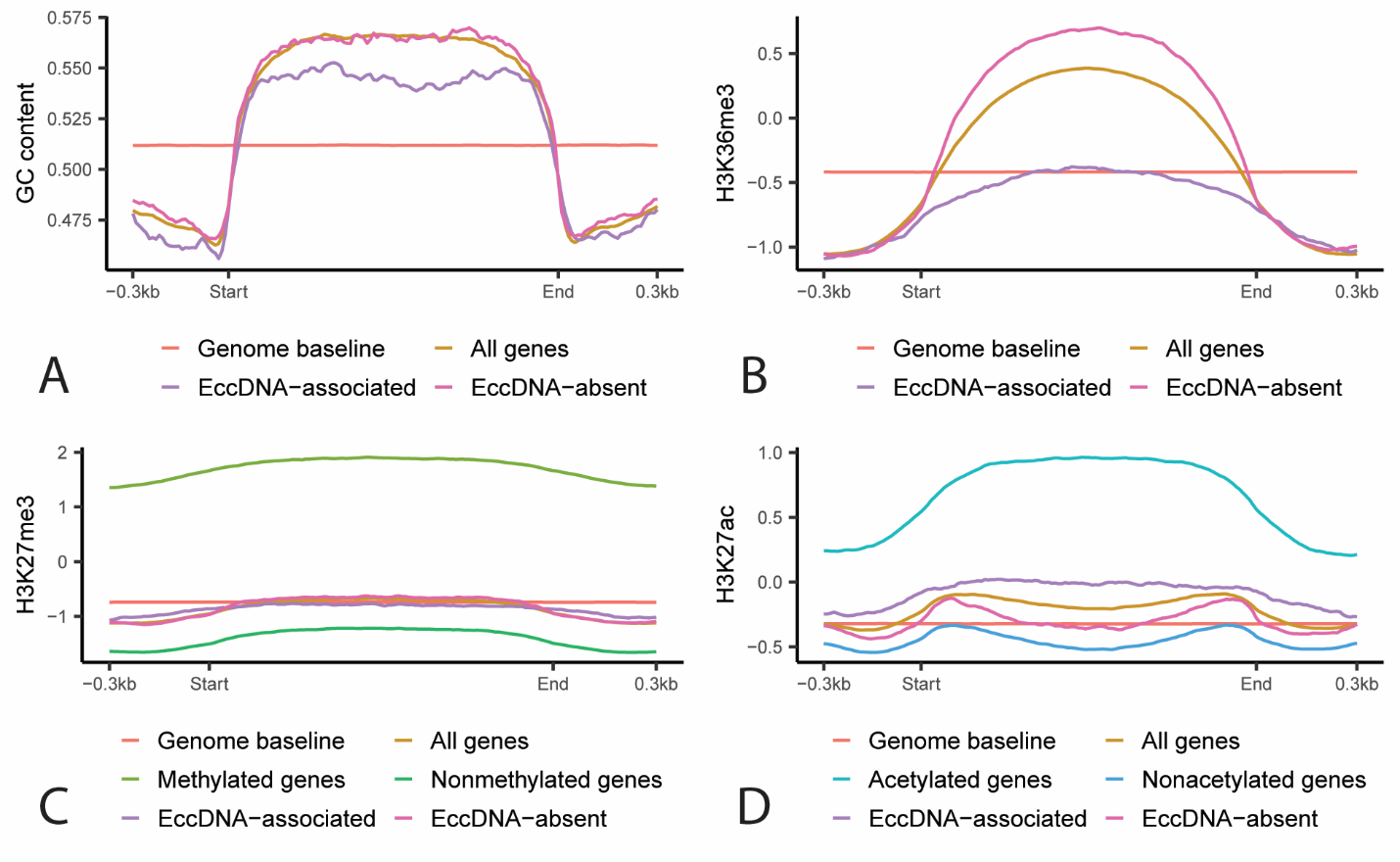

**Fig. S24.** GC content and chromatin marks of eccDNA-associated and eccDNA-absent genes in *M. oryzae*. Profile plots showing the average **A.** percent GC content, **B.** log2 ratio between read coverage for H3K36me3 chromatin immunoprecipitation and input control, **C.** log2 ratio between read coverage for H3K27me3 chromatin immunoprecipitation and input control and **D.** log2 ratio between read coverage for H3K27ac chromatin immunoprecipitation and input control for all *M. oryzae* genes, randomly selected regions of the genome, eccDNA-associated genes, and eccDNA-absent genes. Methylated and nonmethylated genes and acetylated and nonacetylated genes, as defined by Zhang *et al*., are also represented in **C.** and **D.**, respectively.

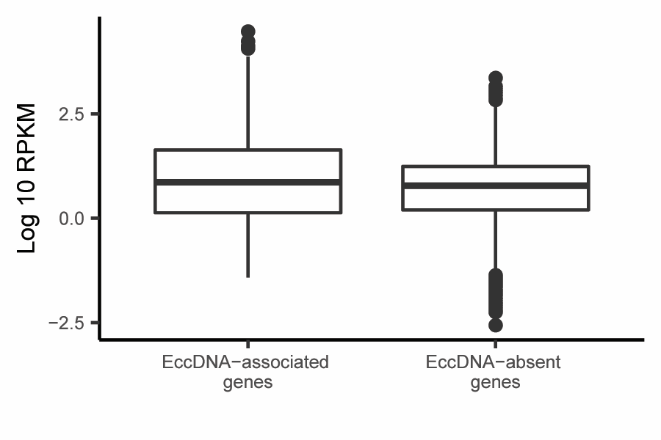

**Fig. S25.** Comparison of expression data between eccDNA-associated genes and eccDNA-absent genes in *M. oryzae*. Box plot showing the log 10 reads per kilobase million (RPKM) averaged across 12 previously published RNAseq samples for eccDNA-associated genes and eccDNA-absent genes.

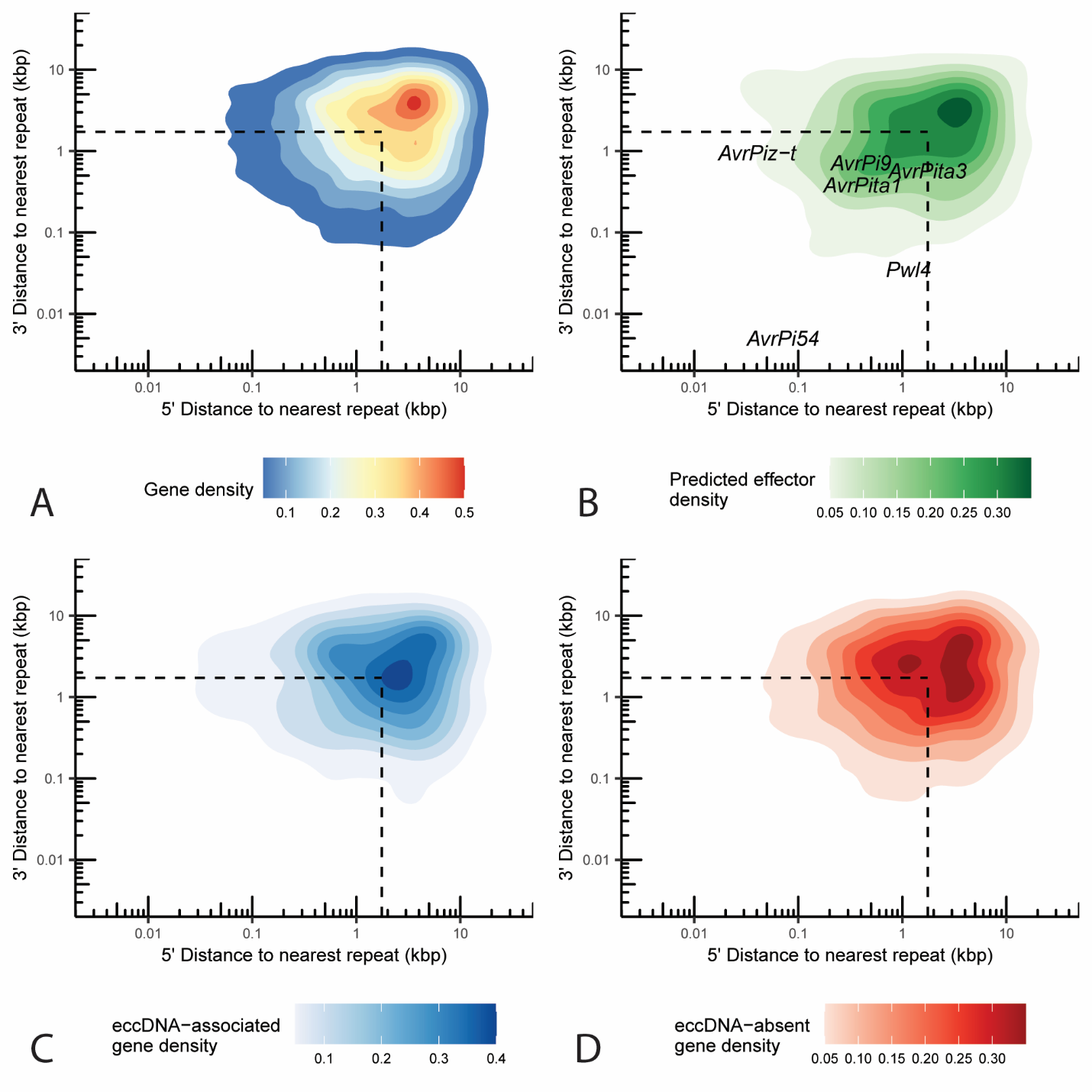

**Fig. S26.** Proximity of *M. oryzae* genes to repeats. Two-dimensional density plot representing the 5’ and 3’ distance to the nearest repeat in the *M. oryzae* Guy11 genome in kilobase pairs for each **A.** gene, **B.** predicted effector, **C.** eccDNA-associated genes, and **D.** eccDNA-absent genes. Known effectors are shown as text in **B.** Dashed lines represent median 5’ and 3’ distance to nearest gene.

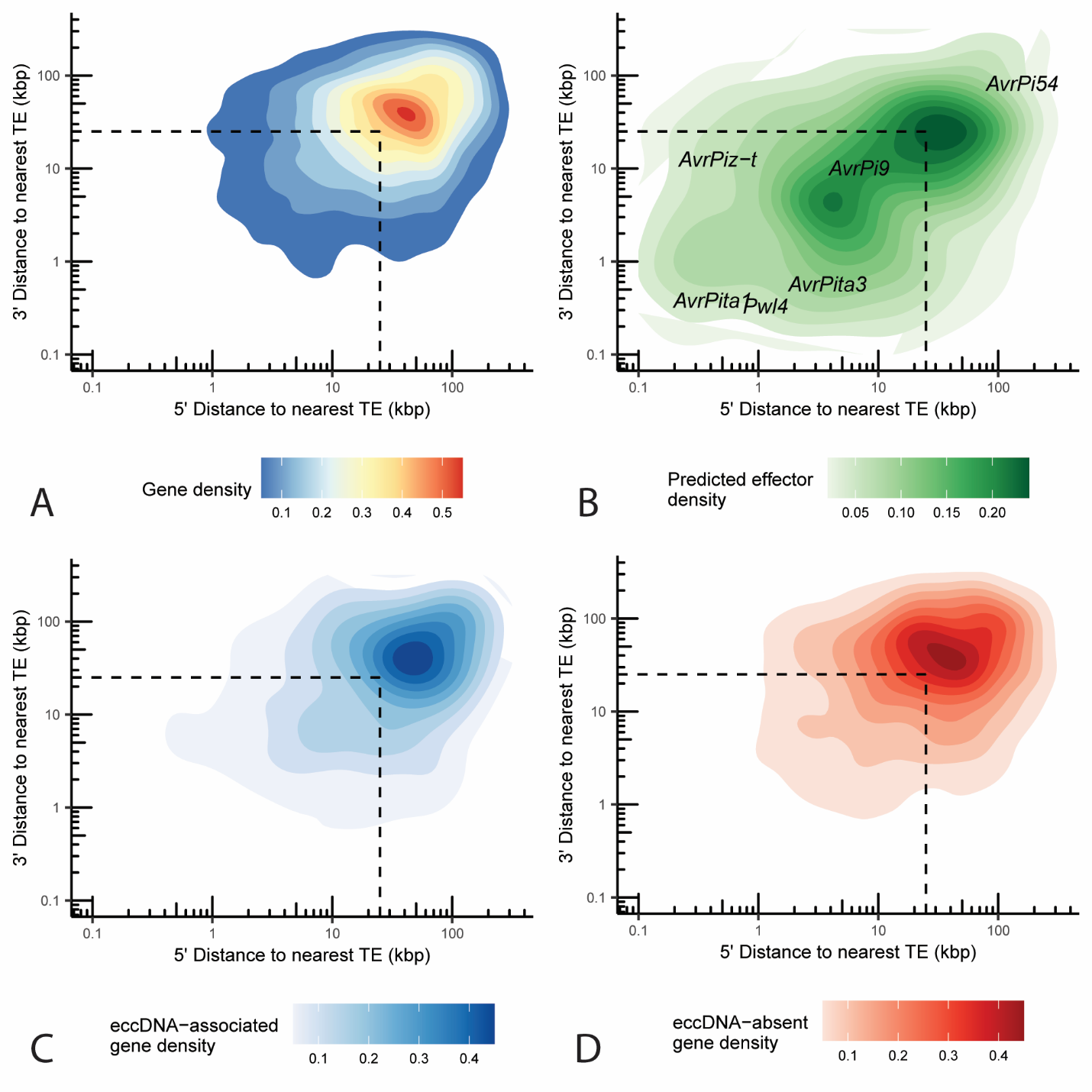

**Fig. S27.** Proximity of *M. oryzae* genes to TEs. Two-dimensional density plot representing the 5’ and 3’ distance to the nearest transposable element in the *M. oryzae* Guy11 genome in kilobase pairs for each **A.** gene, **B.** predicted effector, **C.** eccDNA-associated genes, and **D.** eccDNA-absent genes. Known effectors are shown as text in **B.** Dashed lines represent median 5’ and 3’ distance to nearest gene.

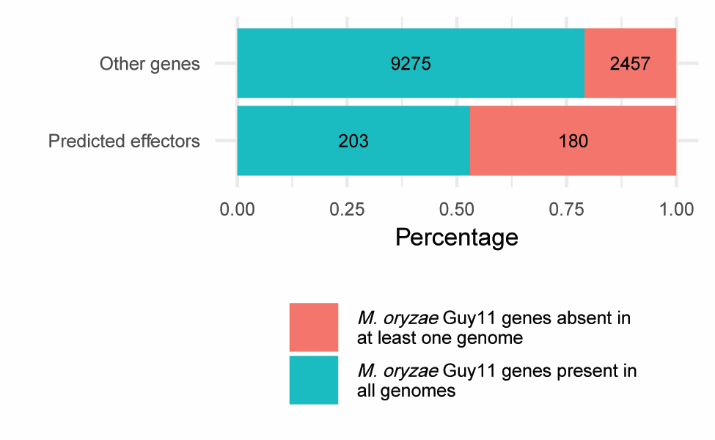

**Fig. S28.** Predicted effectors are prone to presence-absence variation in *M. oryzae*. Stacked bar plot showing the percentage of predicted effectors and all other genes in the *M. oryzae* Guy11 genome that had an ortholog in all other 162 *M. oryzae* genomes analyzed or not. Numbers indicate the number of genes in each category.

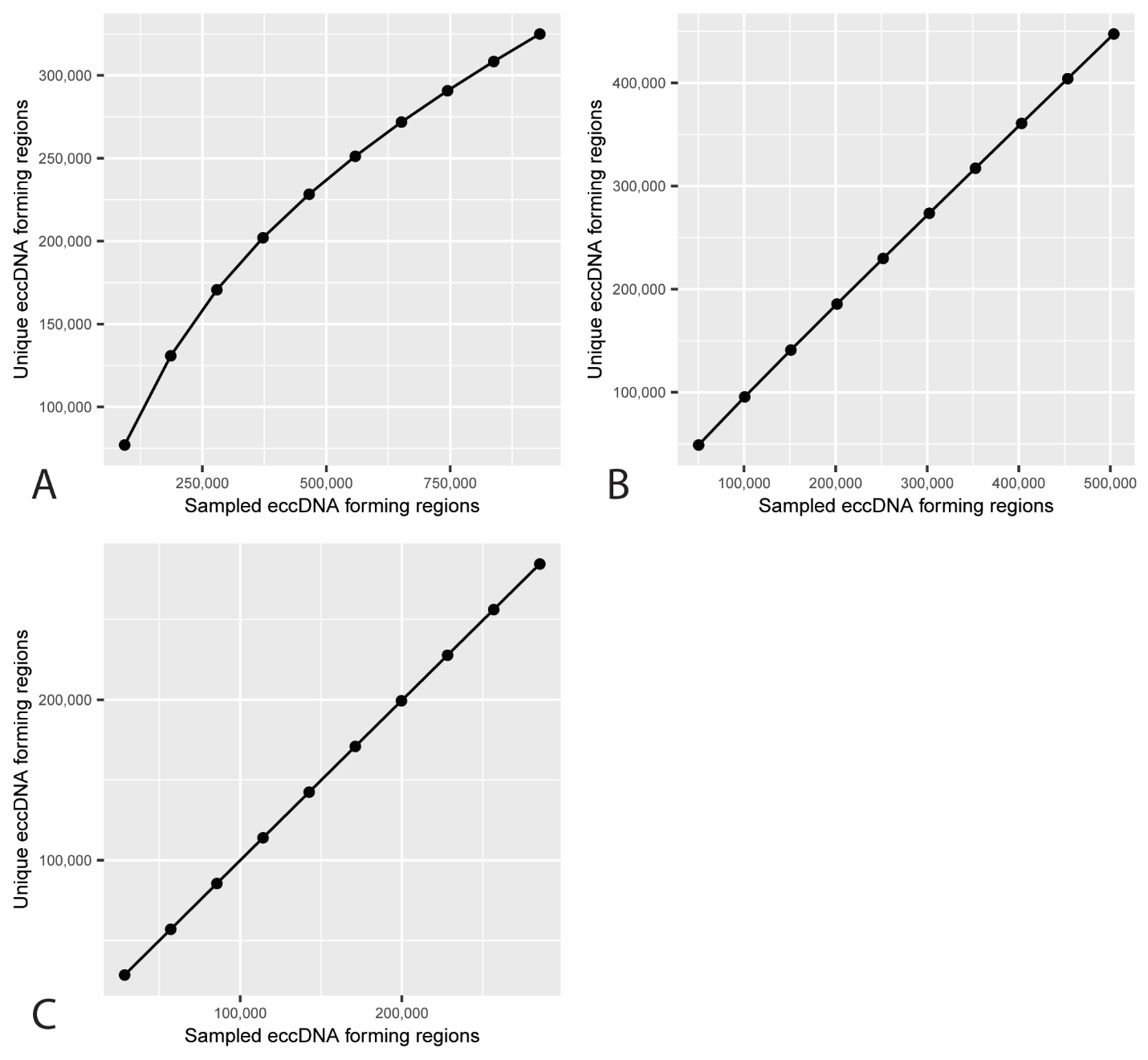

**Fig. S29.** Rarefaction curves for eccDNA forming regions in *M. oryzae*. Rarefaction analysis of the number of unique eccDNA forming regions at different subsamples of eccDNA forming regions across all samples for **A.** LTR-eccDNAs, **B.** large eccDNAs and **C.** microDNAs.

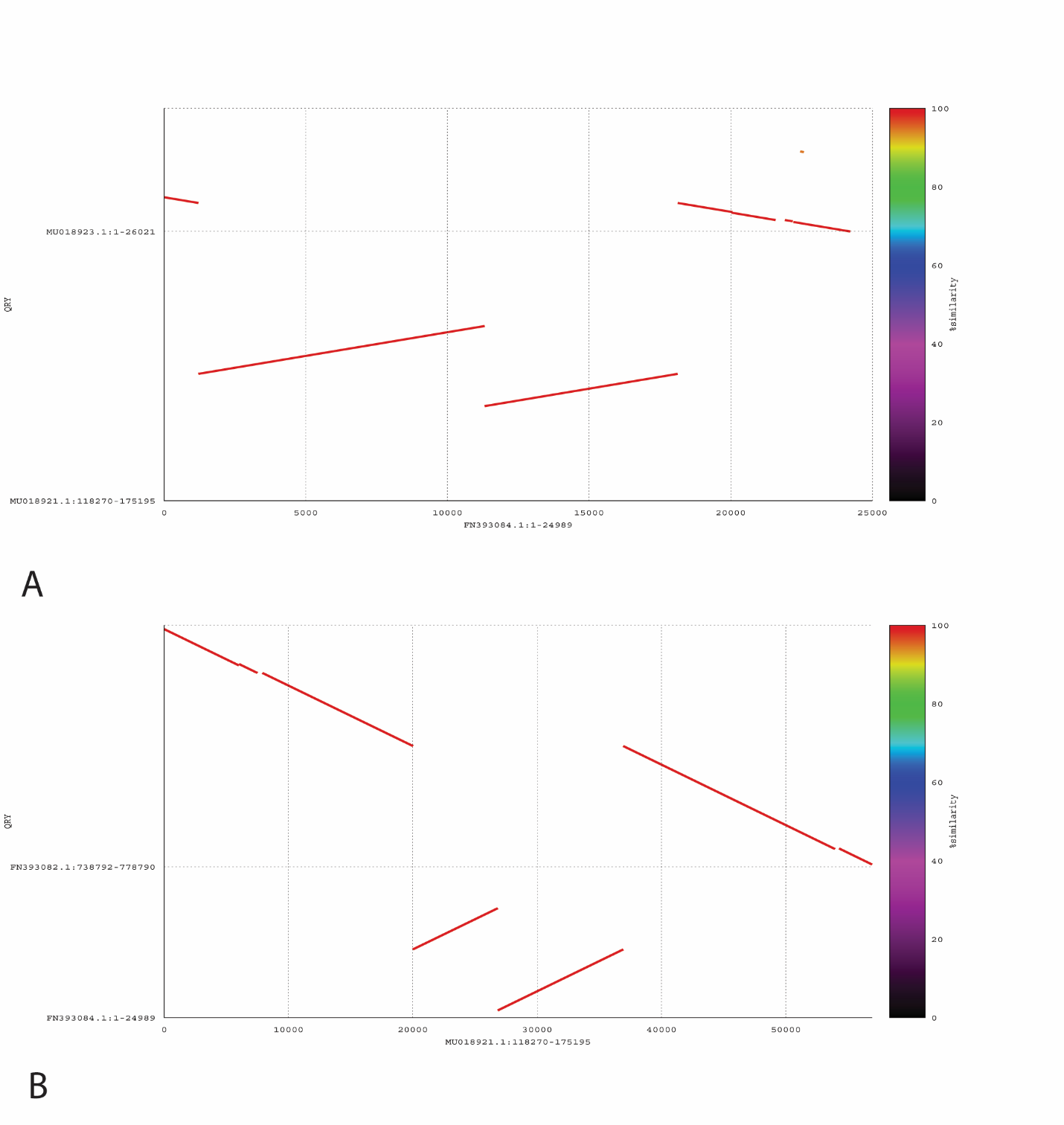

**Fig. S30.** Example of an eccDNA-mediated translocation in wine yeasts. Dot plot alignments between *S. cerevisiae* M22 and *S. cerevisiae* EC1118 genomes showing a DNA translocation likely caused by an eccDNA intermediate in yeast. **A.** A scaffold of the EC1118 genome aligns to two different scaffolds of the M22 genome. **B.** A scaffold of the M22 genome aligns to two different scaffolds of the EC1118 genome.

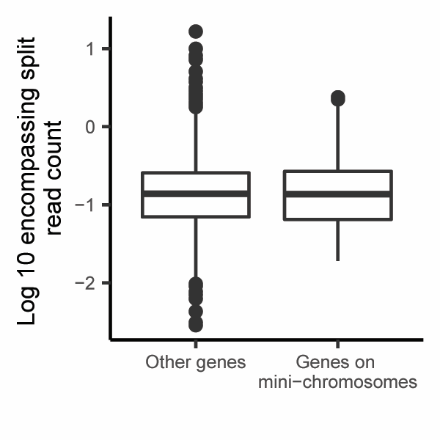

**Fig. S31.** Comparison of encompassing split read counts between genes found on mini-chromosomes in *M. oryzae* and other genes. Box plot showing the log 10 of the number of junction split reads per million reads averaged across biological replicates that fully encompass genes previously found on mini-chromosomes in other strains of *M. oryzae* and other genes.

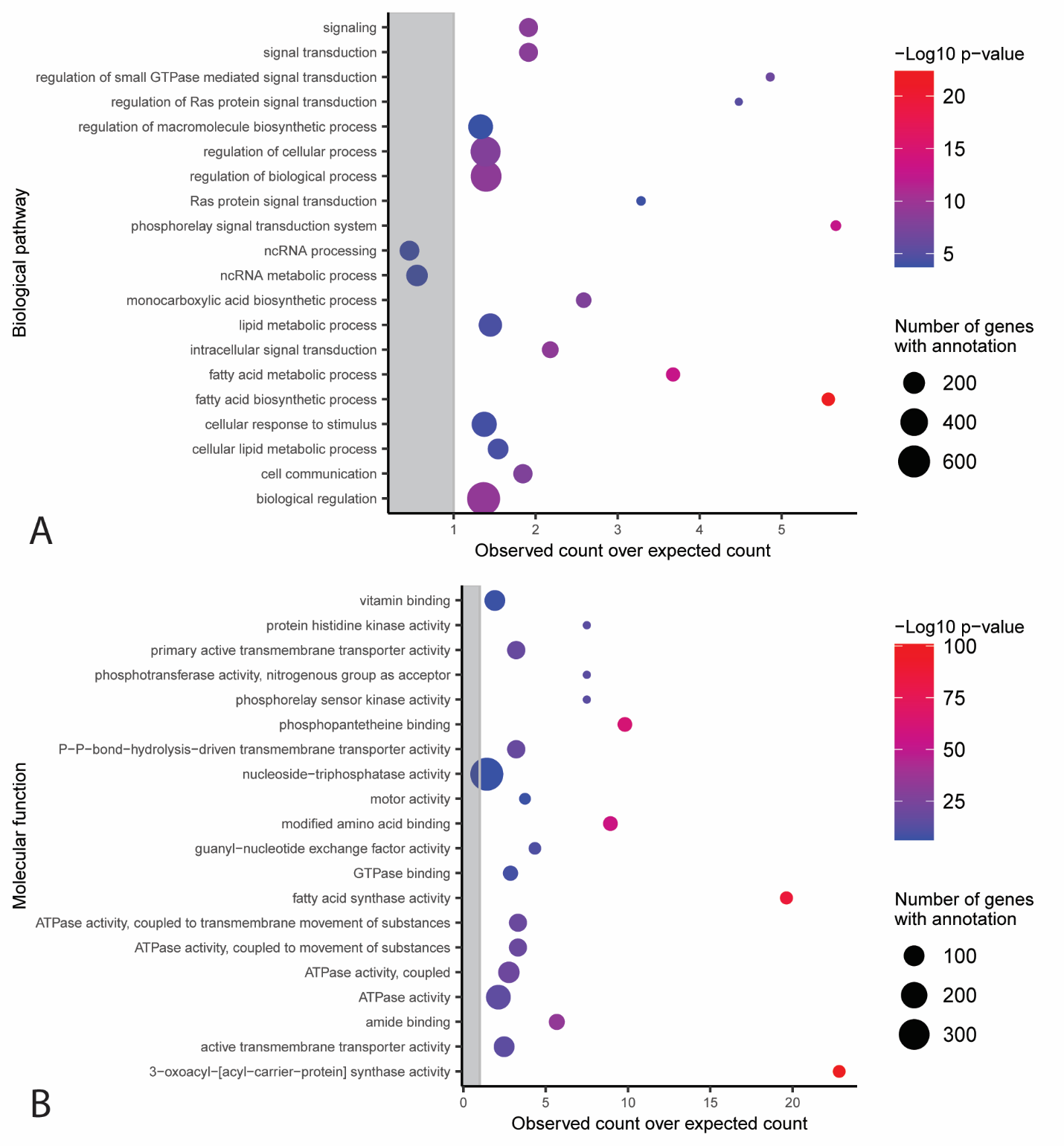

**Fig. S32.** GO terms associated with eccDNA-absent genes. Functional categories in the **A.** molecular function and **B.** biological pathway Gene Ontology with an observed number of eccDNA-absent genes that is significantly different from the expected number with correction for gene length bias (Chi-square test, p < 0.05). The y-axis shows the different functional categories, and the x-axis represents the observed number of genes divided by the expected number of genes in this group. Dots outside of the grey rectangle represent functional categories that are observed more often than expected. The size of dots indicates the total number of genes in the *M. oryzae* genome that belong to each functional category. Only the 20 categories with the largest -log10 p-values are shown.

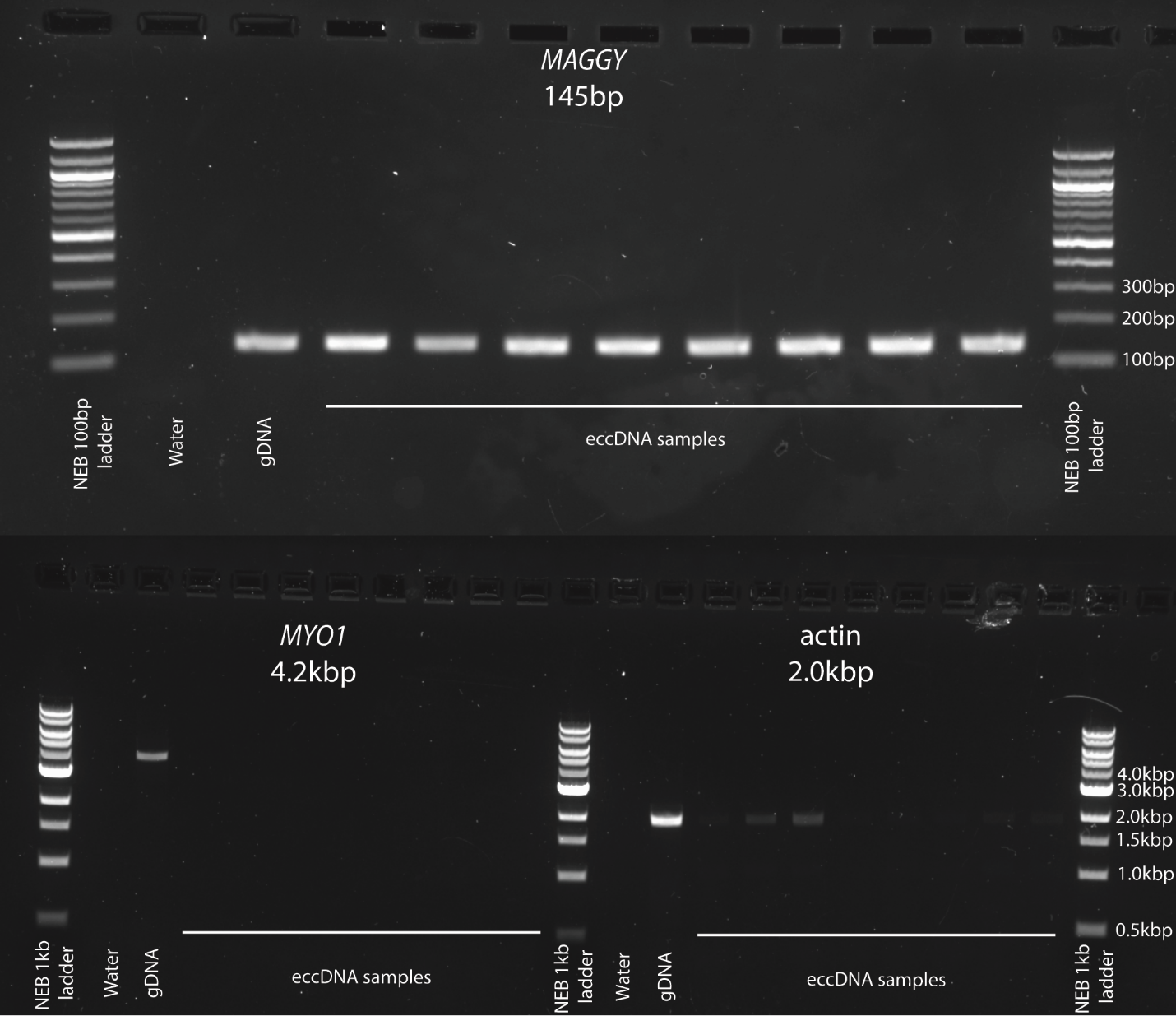

**Fig. S33.** PCR validation of eccDNA-absent genes. Features of interest are listed at the top of each group. One primer set was used per group and the expected product size is written below the feature name. A portion of the *MAGGY* LTR retrotransposon was used as a positive control for amplification. EccDNA samples were grouped by biological replicate and ordered within groups by technical replicate. All samples for each product were from the same PCR reaction.

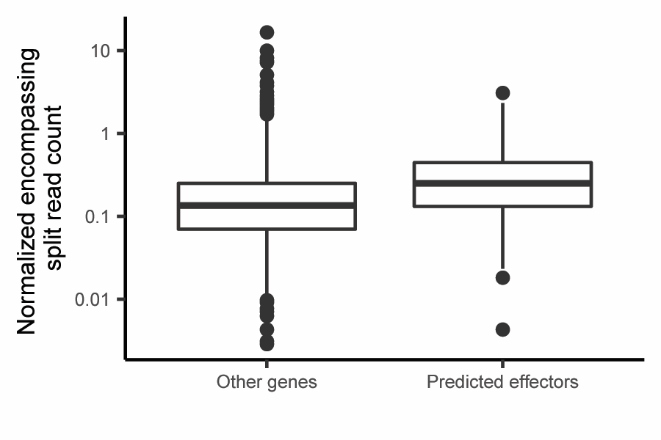

**Fig. S34.** Effectors are enriched in eccDNAs in M. oryzae. Box plot showing the number of fully encompassing junction split reads per million junction split reads averaged across biological replicates for predicted effectors compared to all other genes.

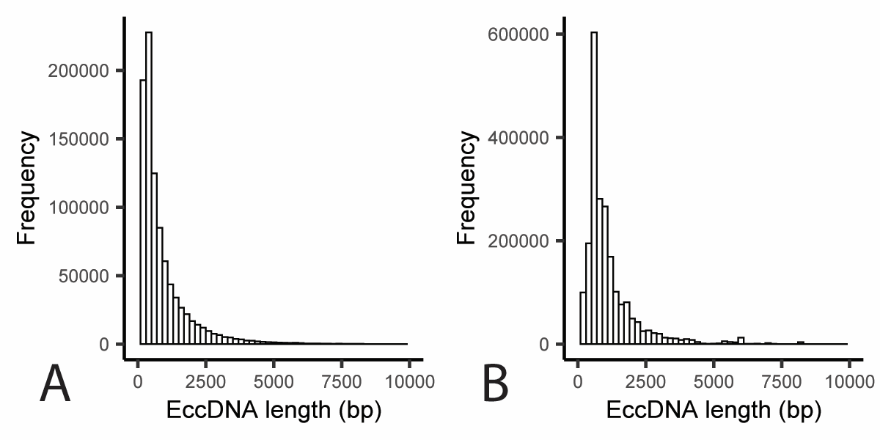

**Fig. S35.** Lengths of eccDNA forming regions in *M. oryzae*. Histograms showing the distribution of candidate eccDNA forming regions in *M. oryzae* for one sequenced sample. **A.** Length distribution of candidate eccDNA forming regions inferred from uniquely mapped reads. **B.** Length distribution of candidate eccDNA forming regions inferred from multi-mapping reads.
