## Additional File 2 for "The extrachromosomal circular DNAs of the rice blast pathogen *Magnaporthe oryzae* contain a wide variety of LTR retrotransposons, genes, and effectors"

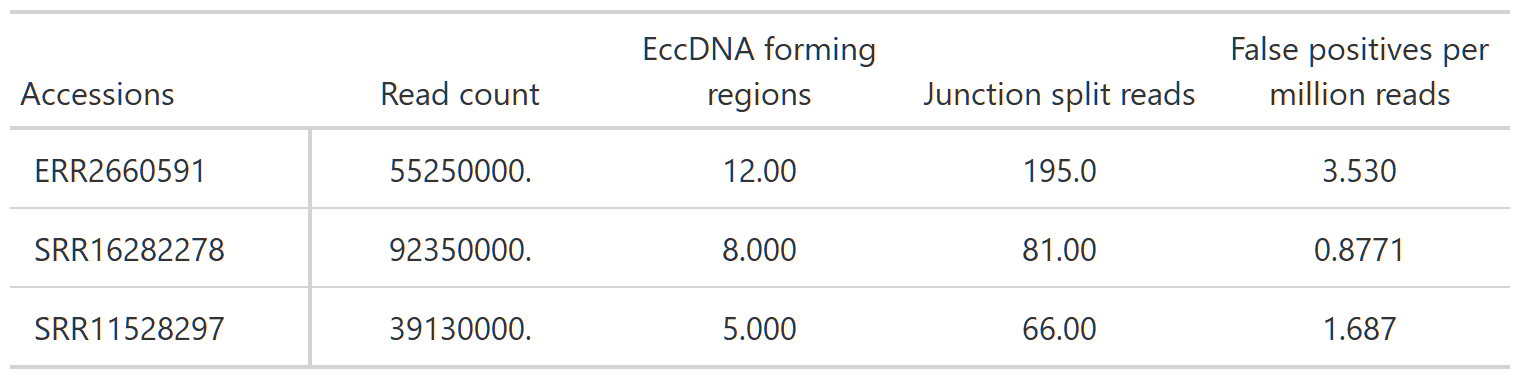

**Table S1.** Number of eccDNA forming regions called using whole genome sequencing data. Read count, eccDNA forming regions inferred, and number of junction split reads found using our pipeline on three previously published whole genome sequencing datasets for *M. oryzae*.

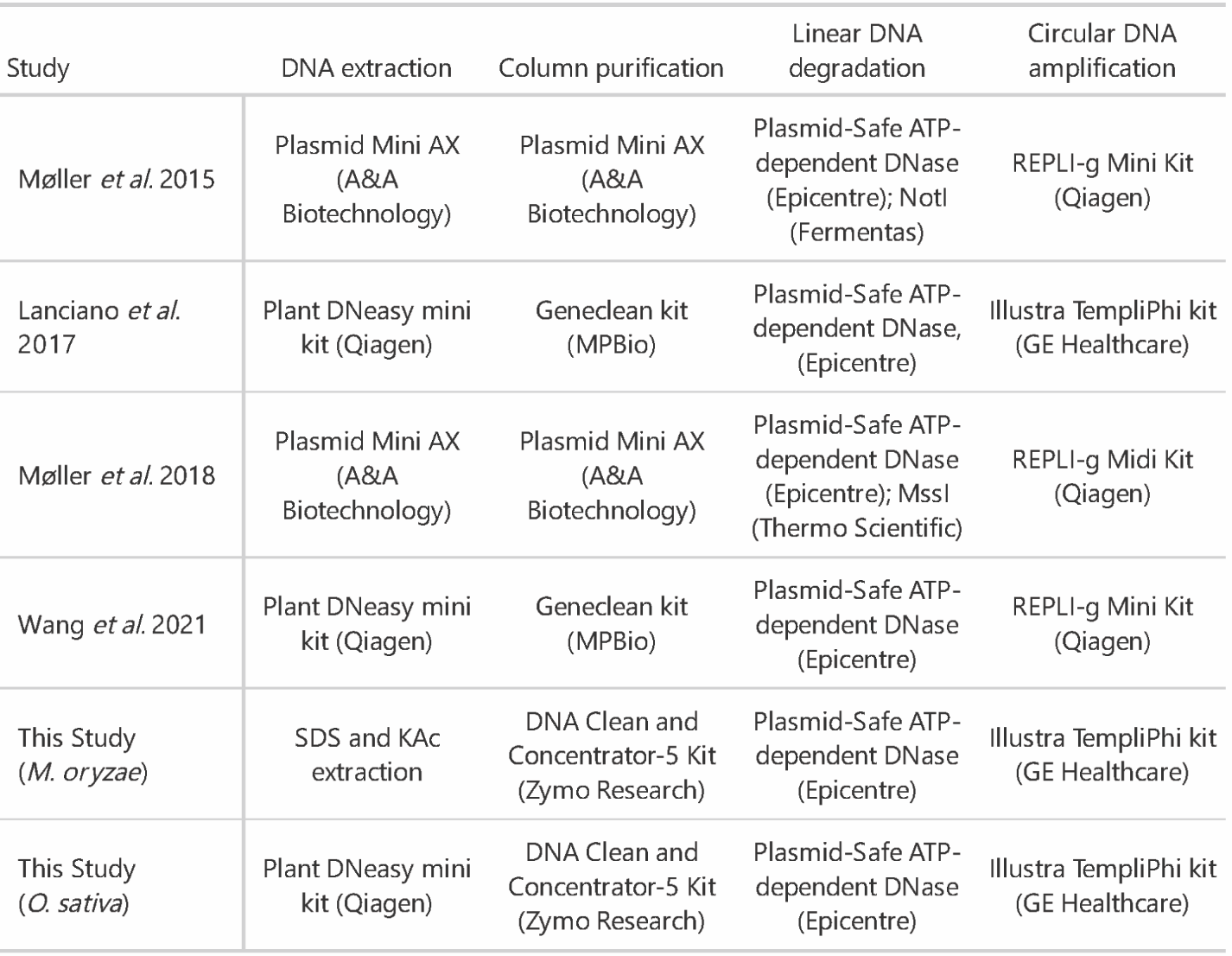

**Table S2.** Summary of protocols used to extract eccDNAs in studies analyzed in this manuscript. DNA extraction kit, column purification kit, linear DNA degradation enzymes and circular DNA amplification enzymes used for all studies whose data was used to compare the circularomes of the organisms discussed in this study.

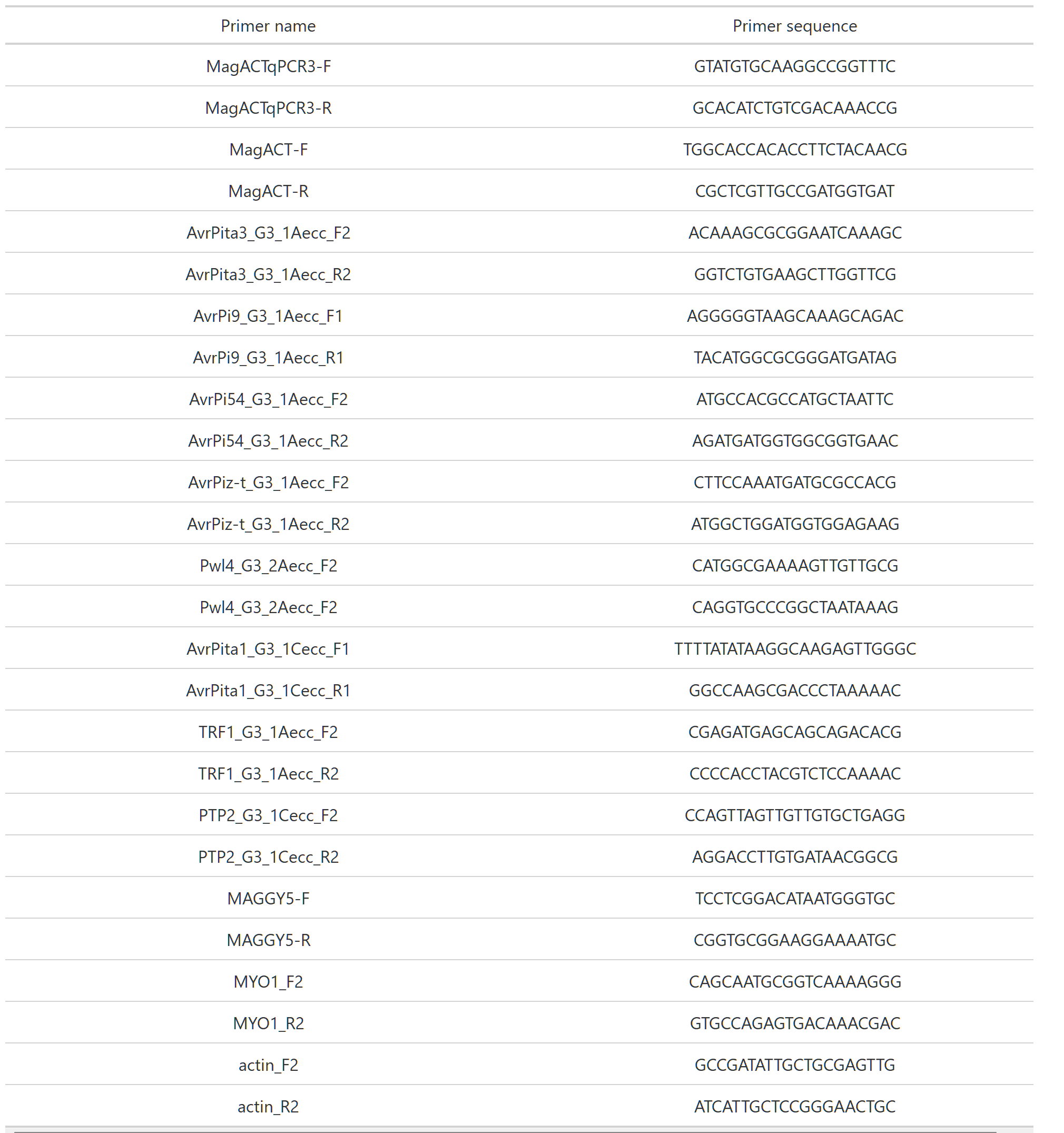

**Table S3.** Primers used for qPCR validation of linear DNA degradation and outward PCR validation of eccDNA forming regions.
